# Dlx2 reprograms the transcriptome and laminar position of glia-derived Ascl1-induced interneurons

**DOI:** 10.64898/2026.01.02.697435

**Authors:** Alexis Cooper, Araceli Garcia Mora, Gabriel Herrera-Oropeza, Ana Beltrán Arranz, Nicolás Marichal, Youran Shi, Sydney Leaman, Karla Lozano Gonzalez, Molly Strom, Hamid Fursham, François Guillemot, Benedikt Berninger

## Abstract

Direct lineage reprogramming of glial cells into neurons offers a promising strategy to repair diseased brain circuits, but engineering defined neuronal subtypes remains challenging. We found that a phospho-site-deficient Ascl1 variant, Ascl1SA6, but not wildtype Ascl1, induces hallmarks of parvalbumin fast-spiking interneurons, raising the question of how closely these induced neurons resemble canonical cortical interneurons and what transcriptional events underlie this process. Single-cell transcriptomic analysis revealed that Ascl1SA6-induced neurons only partially recapitulated canonical interneuron programs and failed to induce the transcription factor Dlx2 and its downstream targets. Co-expression of Dlx2 with Ascl1SA6 restored a more canonical interneuron-like transcriptome, including genes involved in migration, and resulted in neurons occupying laminar positions more typical of endogenous interneurons. These findings provide molecular insights into how Ascl1 posttranslational modifications regulate its transcriptional activity and demonstrate a strategy to engineer induced cortical interneurons that more closely resemble their native counterparts, offering a framework for layer-specific restoration of inhibitory circuits in neurological diseases.

## INTRODUCTION

*In situ* lineage reprogramming of brain-resident glia such as astrocytes and oligodendrocyte progenitor cells (OPCs) into induced neurons (iNs) has emerged as an experimental strategy towards generating new neurons in brain regions devoid of physiological neurogenesis ^1–3^. Such a strategy may allow for regeneration of neurons afflicted by disease or injury and pave the way towards new cell-based therapies for brain repair ^4,5^. Proof-of-principle studies have highlighted the feasibility of *in vivo* neuronal reprogramming across brain areas and glial subtypes ^5–10^.

Lineage reprogramming involves rewriting cell identity by enforcing new transcriptional programs, which is often achieved through forced expression of proneural transcription factors such as achaete scute complex like-1 (Ascl1) or neurogenin-2 among others that drive neural fate decisions during development ^11–14^. However, to fully harness lineage reprogramming for brain repair, a key challenge presents in generating iNs that could adequately replace degenerated or dysfunctional neurons. Ideally, iNs should closely resemble endogenous neurons in their molecular, cellular, and functional properties. Although developmental programs have informed current reprogramming factor cocktails ^5,8,10,15–18^, the molecular logic underlying the conversion of cortical glia into iNs with specific subtype and -class traits has not been elucidated due to the challenging nature of high-throughput isolation and transcriptomically profiling single cells undergoing neuronal reprogramming *in vivo*. Overcoming this challenge will elucidate how individual transcription factors shape fate conversion and subtype specifications while also revealing potential limitations to current approaches ^19^.

Given their role in neurological and neuropsychiatric disease ^20^, cortical interneurons have attracted particular interest as desired outcome of *in situ* reprogramming ^21,22^. In quest of a successful conversion strategy, we have recently discovered that employing a phospho-site-deficient mutant of Ascl1 (Ascl1SA6), previously shown to heighten its neurogenic activity ^23–26^, greatly enhances the induction of neurons with interneuron-specific hallmarks such as parvalbumin expression and fast-spiking, from glia of the early postnatal mouse cerebral cortex^21^.

Here, using high-throughput single-cell and single-nucleus RNA sequencing of over 60,000 cells, we examined transcriptional changes during early stages of glia-to-iNs conversion as well as maturation and circuit integration of iNs. We found that Ascl1SA6 outperformed wildtype Ascl1 in activating cognate neurogenesis-associated gene targets in proliferative glia *in vivo*. Astroglial transcriptomes were more effectively remodeled than those of OPCs, highlighting their greater reprogramming competence. Yet, despite its ability to activate genes associated with specific subclasses of cortical interneurons, Ascl1SA6 failed to fully induce the expression of the Dlx gene family, disclosing an incomplete interneuron program. Co-expressing Dlx2 triggered broad activation of canonical cortical interneuron genes. Intriguingly, Dlx2 co-expression redirected laminar iN position from enrichment in cortical layer 1 to more canonical positions in layers 2/3, indicating that Dlx2 not only reshapes the transcriptome of iNs but also their overall cellular behavior and biology. These findings break ground for closing the gap between induced and endogenous neurons for future application of induced neurogenesis in brain repair. Our study provides a resource available for browsing at Cellxgene.

## RESULTS

### Phospho-site-deficient Ascl1 outperforms wildtype Ascl1 in remodeling glial transcriptomes toward neurogenesis

To decipher the molecular underpinnings of why phospho-site-deficient Ascl1 (referred here to as Ascl1SA6) is more potent than wildtype Ascl1 (referred here to as “Ascl1”) at inducing glia-to-neuron conversion ^21,27^, we co-injected retroviruses encoding either Ascl1 or Ascl1SA6 (with (+) or without (-) anti-apoptotic factor Bcl2 ^16^) into the somatosensory cortex of postnatal day 5 (P5) mice (Fig. 1a, Extended Data Fig. 1a). At 4 days post-injection (dpi), Ascl1/Bcl2 co-transduced cells largely retained glial morphologies, whereas a subset of Ascl1SA6/Bcl2 co-expressing cells already displayed more neuron-like morphologies (Fig. 1b). To delineate the molecular programs and transcriptional transitions underlying these initial stages of glia-to-neuron reprogramming, we performed single-cell RNA sequencing of retrovirus-transduced cells isolated from cortices ^28,29^ injected with these reprogramming factor combinations or a viral vector encoding only a red fluorescent protein (Control) at 4 dpi (Fig. 1a; Extended Data Fig. 1a).

**Figure 1.**
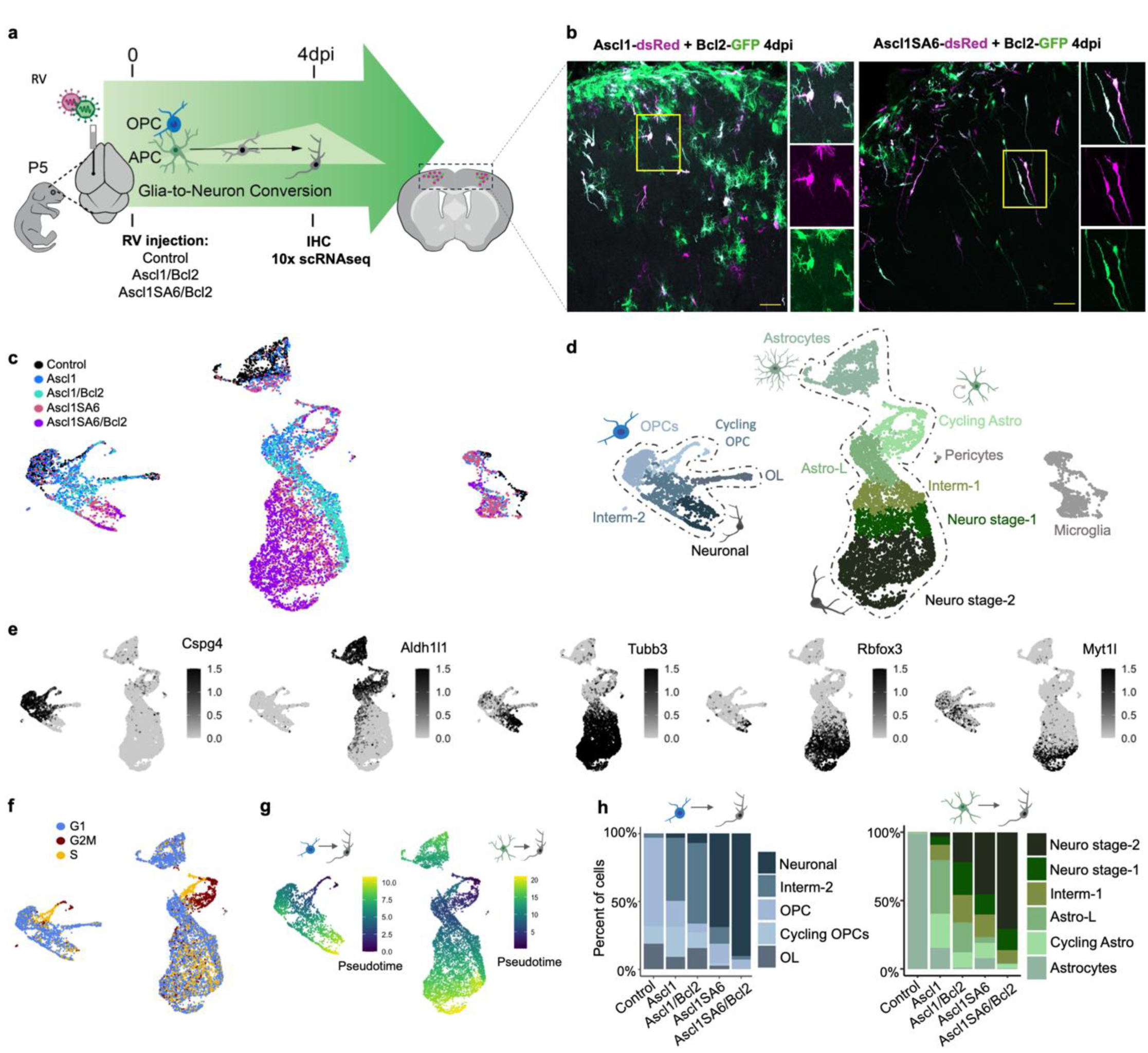
Phospho-deficient Ascl1 (Ascl1SA6) more effectively remodels glial transcriptomes toward neurogenesis than wildtype Ascl1. **a,** Experimental design. Retroviral constructs encoding Ascl1 or Ascl1SA6 with and without Bcl2 were injected into the somatosensory cortex of wild-type (C57BL/6J) mice at postnatal day 5 (P5). Reporter-positive cells were isolated at 4 days post-injection (dpi) for 10x Genomics single-cell RNA sequencing**. b,** Confocal images at 4 dpi showing co-transduced cells (magenta: Ascl1 or Ascl1SA6; green: Bcl2; white: merge). Cells expressing Ascl1SA6/Bcl2 displayed early neuronal morphologies, while Ascl1/Bcl2 cells retained glial features**. c,** UMAP visualization of retrovirally transduced cells colored by condition on merged UMAP embedding. Ascl1SA6/Bcl2-transduced cells show clear enrichment in neuron-like regions, whereas Ascl1/Bcl2 cells remain largely glial. Cell numbers per condition: Control (n=1,524), Ascl1 (n=2,391), Ascl1/Bcl2 (n=1,387), Ascl1SA6 (n=2,764), Ascl1SA6/Bcl2 (n=1,755). For visualization, UMAPs were down-sampled to equal cell numbers (1,000/condition). **d,** UMAP colored by annotated cell identity showing merged single-cell transcriptomes. Clusters include astrocytes-to-neuron, OPCs-to-neuron, microglia, and pericytes. **e,** UMAP feature plots colored in black by gene expression, highlighting lineage and neurogenic markers across all cells. **f,** UMAP embedding of single-cell transcriptomes dataset at 4dpi colored by cell cycle phase (G1, S, G2/M). **g,** Cells in UMAP projection colored by pseudotime in astrocyte-to-neuron clusters (right) reprogramming and OPC-to-neuron clusters (left). **h,** stacked bar plots showing cell type proportions across conditions, Left OPC-derived and right Astrocyte-derived cells. Ascl1SA6/Bcl2 shifted astrocyte- and OPC-derived populations toward neuron-like states more efficiently than Ascl1/Bcl2.

We obtained transcriptional profiles from 27,234 high-quality single cells and detected retroviral transcripts in the majority of these (Extended Data Fig.1b; Supplementary Fig. 1a). Cells expressing distinct reprogramming factor combinations (i.e. Ascl1 vs Ascl1SA6 +/- Bcl2) occupied overlapping but distinct clusters in a low dimensional (Uniform Manifold Approximation and Projection; UMAP)) embedding of their transcriptomic profiles, suggesting that Ascl1 and Ascl1SA6 might initiate early diverging transcriptional trajectories across glial populations (Fig. 1c-e; Extended Data Fig.1c, Supplementary Fig. 2a). In the absence of reprogramming factors (Control), most retrovirus-transduced cells were identified as astrocytes (∼47%) and oligodendrocyte lineage cells (∼41%), which included both proliferative (Cycling OPC and Cycling Astro) and differentiated (OL and Astrocytes) subpopulations (Extended Data Fig.1c; Supplementary Fig. 2b; Supplementary Table 1a).

To resolve transcriptomic dynamics of fate conversions, we applied pseudotime inference ^30^ to astrocyte- and OPC-derived cells, anchored in the mitotic subset of the respective glial populations (Fig. 1f,g). Retroviral expression of Ascl1 or Ascl1SA6 +/- Bcl2, reshaped the transcriptional landscape of astrocyte- and OPC-derived clusters, which transitioned through intermediate states toward neuronal stages (Fig 1d). These were characterized by the sequential induction of neural progenitor genes (*Hey1, Hes5, Notch1*), markers of early neurogenesis (*Dll1, Sox11, Tubb3*), and transcripts of more mature neurons (*Rbfox3, Srrm4, Myt1l*) (Fig. 1e; Supplementary Fig. 2a). Neuronal progression was most pronounced in Ascl1SA6-expressing cells, whereas Ascl1-expressing populations largely retained glial identities (Fig. 1h; Supplementary Fig. 2b). Within OPC-derived populations, Ascl1 downregulated radial glial genes (*Vim, Aldoc, Sox9*) while maintaining OPC differentiation genes (*Sox10, Cspg4*) (Extended Data Figs. 1d–f). Microglia were sporadically transduced under all conditions but did not upregulate neuronal markers. Sub-clustering analyses confirmed that microglia-specific genes were expressed similarly across all conditions (Supplementary Figs. 3a-c). Overall, Ascl1SA6, especially when co-expressed with Bcl2, markedly enhanced glia-to-neuron conversion in both astrocytes and OPCs compared to Ascl1 at 4dpi.

### Astrocytes exhibit greater competence in activating/suppressing canonical neural stem cell target genes of Ascl1 than OPCs

While both, proliferative astrocytes and OPCs can initiate neurogenic reprogramming, the molecular landscape provided by these two glial subtypes may dictate the specific transcriptional programs activated by Ascl1 and/or Ascl1SA6. Thus, we examined whether astrocytes and OPCs alter their transcriptomes by activating/suppressing similar or distinct target genes to those previously identified in ventral telencephalic neural stem cells (NSCs) during interneuron differentiation ^31,32^. Thus, we computed Ascl1 target module scores of up-and down-regulated genes in NSCs ^31^ and projected them onto the UMAP embedding of the above 4-day data set (Fig. 2a; Extended Data 2a). First, we analyzed whether astrocytes and OPCs differ in their overall regulation of these genes in response to wildtype and phospho-site-deficient Ascl1. Reassuringly, we found that Ascl1SA6 was more potent than Ascl1 in regulating *bona fide* targets (Extended Data 2b). However, Ascl1SA6 was significantly more effective in up- and down-regulating these targets in astrocytes as compared to OPCs (Fig. 2b,c; Extended Data 2c; Supplementary Table 2). Adding Bcl2 enhanced target regulation both by wildtype and phospho-site-deficient Ascl1 (Fig. 2b,c; Extended Data 2c). Previous work in ventral telencephalic NSCs showed that subsets of Ascl1 target genes follow distinct dynamics regarding onset and maintenance of gene activation/repression, allowing to group these targets into distinct clusters (Extended Data 2a) ^31^. Here, we projected module scores of these clusters onto the UMAP embedding and found that in astrocytes, target genes closely followed the dynamics observed in NSCs (Fig. 2d,e; Extended Data 2d; Supplementary Fig. 4-6). For instance, cluster 1 and 4 of upregulated genes exhibited a highly distinct module score distribution across pseudotime, consistent with cluster 1 genes being transiently expressed during neuronal reprogramming while cluster 4 becoming activated at a more advanced stage (Fig. 2d). While OPCs followed a similar trend, temporal dynamics of Ascl1 target gene expression were less salient (Fig. 2d,e; Supplementary Table 3).

**Figure 2.**
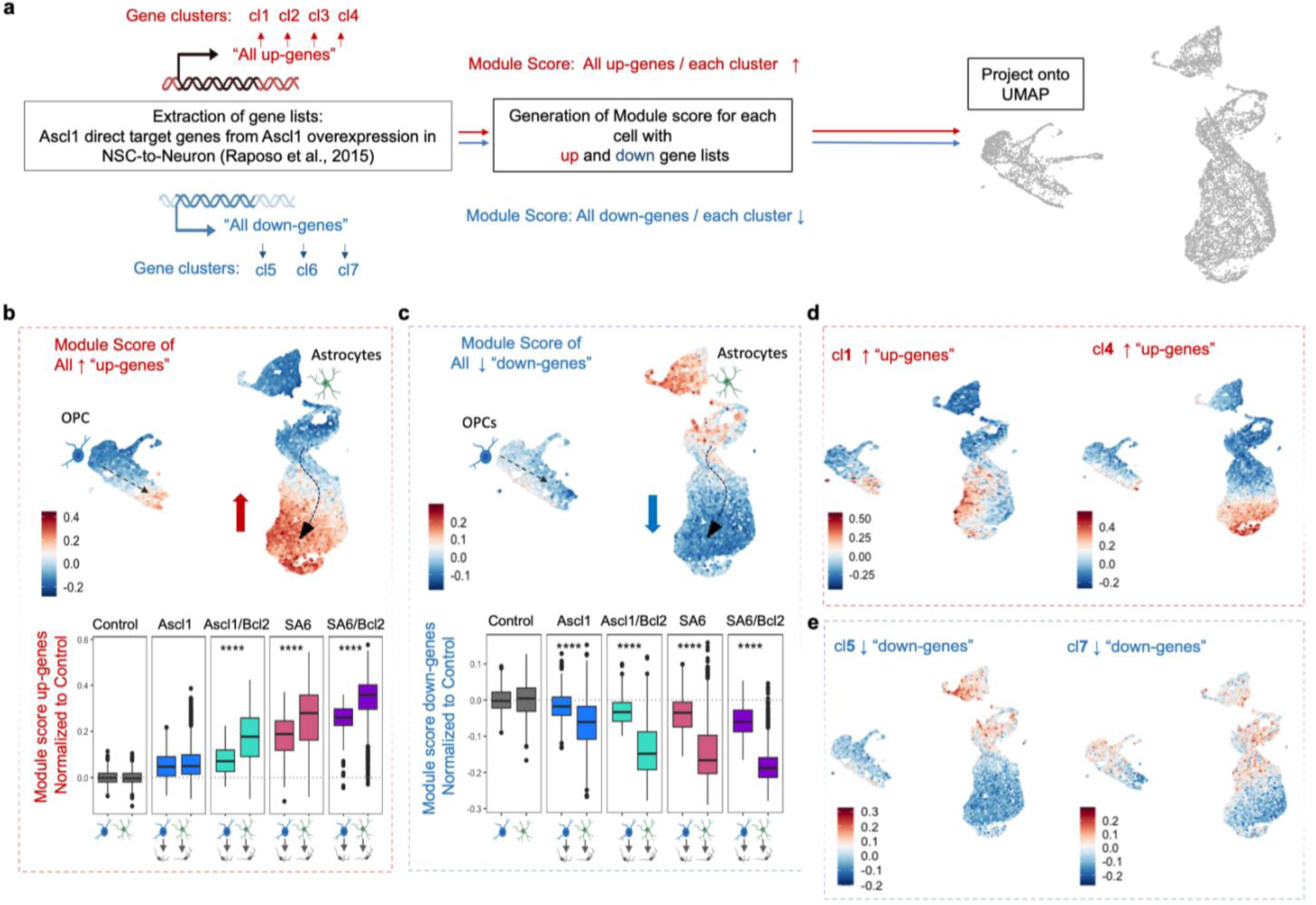
Astrocytes exhibit greater competence in activating/suppressing canonical neural stem cell target genes of Ascl1 than OPCs. **a,** Analysis workflow: Gene lists of up- and downregulated targets upon Ascl1 overexpression in neural stem cells (NSCs) were obtained from Raposo et al., 2015. Module scores were calculated for each dynamically regulated gene cluster as well as for the combined set of clusters for each category (“up” and “down”) and projected onto UMAP of condition (Control, Ascl1, Ascl1/Bcl2, Ascl1SA6 and Ascl1SA6/Bcl2). On right: Module scores projected onto UMAP of 4 dpi cells. **b,c**, UMAP projections of merged data with module scores for the combined upregulated clusters (**b**) and downregulated clusters (**c**). Boxplots show normalized Ascl1 target UP and DOWN module scores across OPC- (O2N) and astrocyte-derived (A2N) lineages per retroviral condition, normalized to Control. Astrocytes (A2N) consistently scored higher than OPCs (O2N), with significant differences as assessed by Wilcoxon rank-sum test: UP module: Ascl1 (ns, p = 0.522), Ascl1/Bcl2 (****, p = 4.09×10⁻³⁰), Ascl1SA6 (****, p = 7.18×10⁻³¹), Ascl1SA6/Bcl2 (****, p = 6.34×10⁻²²); DOWN module: Ascl1 (****, p = 5.86×10⁻⁶⁸), Ascl1/Bcl2 (****, p = 4.01×10⁻⁷²), Ascl1SA6 (****, p = 2.31×10⁻¹¹³), Ascl1SA6/Bcl2 (****, p = 2.07×10⁻⁴⁸). **d,** UMAP projection show module scores for upregulated clusters 1 and 4. **e,** UMAP projection show module scores for downregulated clusters 5 and 7.

These data indicate that, with respect to the identity and expression dynamics of Ascl1 target genes, astrocytes more closely resemble NSCs, providing an explanatory framework for why, in this experimental model, long-term neuronal reprogramming is more successful in astrocytes than in OPCs ^21^.

### Ascl1SA6 outperforms Ascl1 in overcoming roadblocks of astrocyte-to-neuron reprogramming but fails to induce Dlx gene activation

Given their heightened neuronal reprogramming competence, we next zoomed in on conversion of astrocytes into neuron-like cells at 4 dpi. Re-clustering and high-resolution UMAP embedding suggested stark differences in the transcriptomes between wildtype and phospho-site-deficient Ascl1 expressors (Fig. 3a,b; Extended Data Figs. 3a-c). Pseudo-bulk RNA expression analysis revealed that Ascl1SA6 expressors, particularly when co-expressing Bcl2, were more efficient in downregulating astrocyte fate-related genes as compared to Ascl1 expressors. Conversely, they also up-regulated neurogenesis-related genes (e.g., *Srrm4, Rbfox3*) more effectively (Fig. 3c). On a more global level, Gene Ontology (GO) terms showed that wildtype and phospho-site-deficient Ascl1 expressors differentially lost and gained hallmarks related to glio- and neurogenesis, respectively (Fig. 3d; Supplementary Table 4). This comparison across different reprogramming conditions furthermore highlighted the effect of Bcl2 promoting the acquisition of neuronal characteristics.

**Figure 3.**
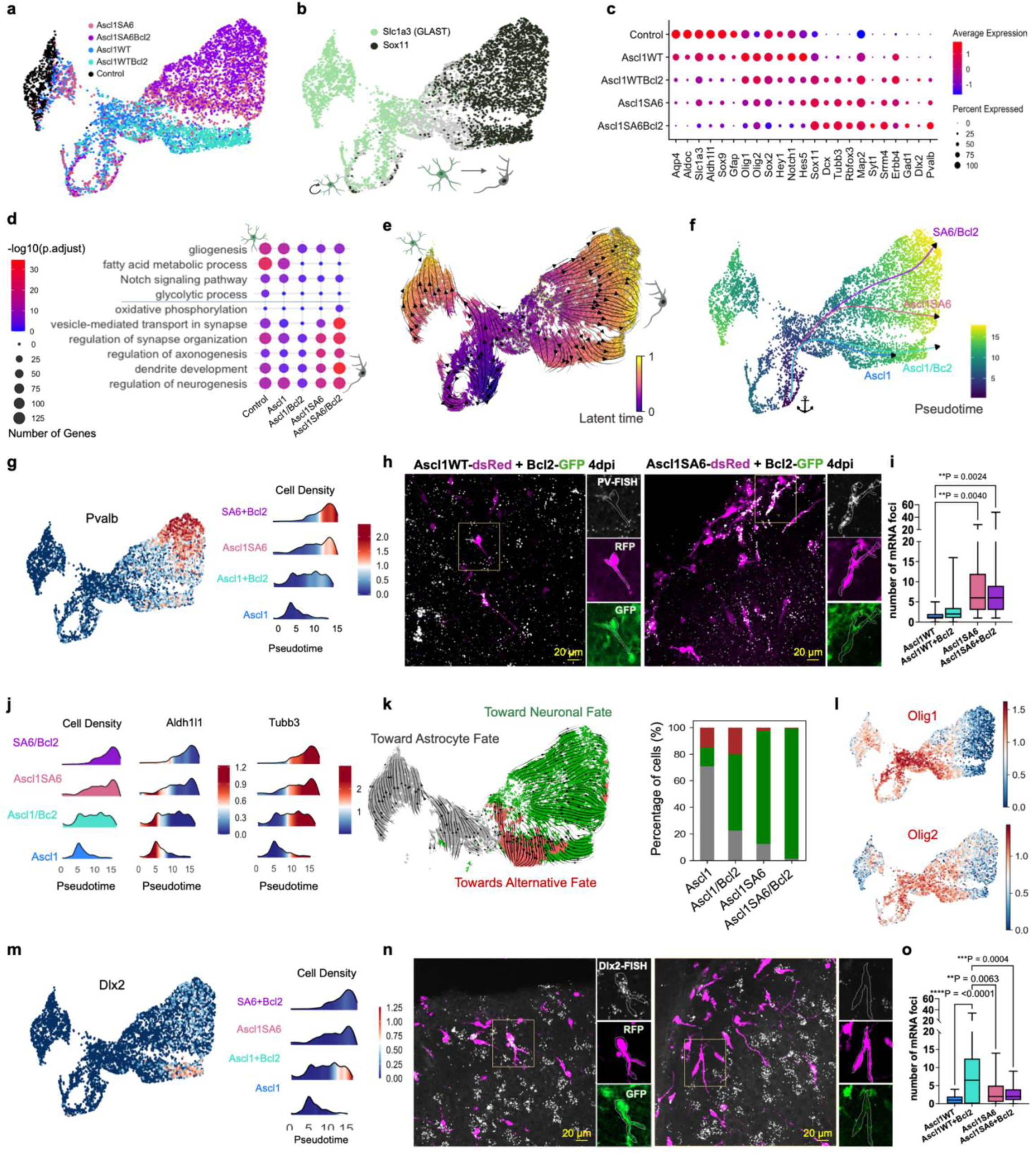
Ascl1SA6 outperforms Ascl1 in overcoming roadblocks of astrocyte-to-neuron reprogramming but fails to induce Dlx gene activation. **a,** UMAP embedding colored by five RV samples. Embedding shows transcriptomes of 5963 cells (633 Mscarlet Control, 1068 Ascl1, 1339 Ascl1/Bcl2, 1423 Ascl1SA6 and 1500 Ascl1/Bcl2 transduced cells. **b,** UMAP feature plot with dual expression of astrocyte marker Slc1a3 (GLAST) and neuronal marker Sox11, indicating transition from astrocytes to neurons. **c,** Dot plot showing the proportion of cells (dot size, % of total) and expression levels (color intensity) of selected marker genes across annotated cell type clusters. **d,** Bubble plot of enriched Gene Ontology (GO) terms for all 5 samples. Significance was determined using a Benjamini–Hochberg-adjusted hypergeometric test. **e,** RNA velocity-derived latent time shown as a gradient (blue to yellow), overlaid with unified velocity streams and individual cell vector arrows. Pseudotime endpoints correspond to astrocytes and neuronal stage 2. **f**, Pseudotime and only individual RV-specific lineages arising from cycling cells towards neuronal stages inferred using the Slingshot algorithm (Street et al., 2018) from the merged dataset. **g**, UMAP visualization of *Pvalb* gene expression (left); expression overlaid on density along Pseudotime (right). h, *Pvalb* expression is not observed in Ascl1/Bcl2-transduced cells (left) but is acquired in Ascl1SA6/Bcl2-transduced cells (right) as shown by RNAscope at 4 dpi. Scale bar, 20 µm. **i,** Quantification of *Pvalb* mRNA transcripts (RNAscope dot counts per cell) in Ascl1, Ascl1/Bcl2, Ascl1SA6, and Ascl1SA6/Bcl2-transduced cells. Values (mean ± SD): 1.7 (n = 21 cells), 3.5 (n = 18), 8.8 (n = 23), and 8.4 (n = 40), respectively, from 3 mice per group. Statistical analysis: one-way ANOVA with Tukey’s post hoc test. Ascl1 versus Ascl1/Bcl2: ****P=0.0001, Ascl1/Bcl2 versus Ascl1SA6: **P=0.0063 and Ascl1/Bcl2 versus Ascl1SA6/Bcl2: ***P=0.0004. (see Extended Data Table for Fig. 3). **j,** Density plots of cells along Pseudotime, grouped by sample demonstrating progression of cells. Expression levels of *Aldh1l1* (astrocytic) and *Tubb3* (neuronal) genes are colored within density plot along pseudotime. **k, Left:** RNA velocity stream plots and arrows of individual cells colored by fate trajectory: toward astrocyte fate (Astrocytes, Astrocyte-like), toward neuronal fate (Intermediate, Neuron-like 1 and 2), and towards alternative fate (Intermediate, Neuron-like 1 and 2). Right: corresponding proportions in a stacked bar plot (Control sample excluded). **l,** UMAP visualization of selected genes enriched in cells moving towards alternative fate. **m,** UMAP showing *Dlx2* gene expression (left); expression profile in density plot over pseudotime (right). n, *Dlx2* is expressed in Ascl1/Bcl2-transduced cells (left) and is acquired in Ascl1SA6/Bcl2-transduced cells (right), as revealed by RNAscope at 4 dpi. Scale bar, 20 µm **o,** Quantification of *Dlx2* mRNA transcripts (RNAscope dot counts per cell) in Ascl1, Ascl1/Bcl2, Ascl1SA6, and Ascl1SA6/Bcl2-transduced cells. Values (mean ± SD): 1.05 (n = 20), 8.1 (n = 18), 3.3 (n = 25), and 2.6 (n = 39), respectively, from 3 mice per group. Statistical analysis: one-way ANOVA with Tukey’s post hoc test, Ascl1 versus Ascl1SA6: **P=0.0024 and Ascl1 versus Ascl1SA6/Bcl2: **P=0.0040. (see Extended Data Table for Fig. 3).

These findings raised the question of whether Ascl1 and Ascl1SA6 expressors constitute different lineages. RNA velocity and latent time ^33^ analyses indicated that cells diverged from a cluster of cycling cells towards two main cellular fates, i.e., astrocyte or neuronal fate (Fig. 3e). To corroborate that cells dividing at 4dpi could still give rise to lineage-converted cells, and therefore validate the anchorage of lineages in the cycling cluster, we pulse-chased Ascl1SA6/Bcl2 expressing cells with EdU (5-ethynyl-2′-deoxyuridine) at 4dpi and found that 11.67% (±5.85%) indeed gave rise to induced neurons (iNs) at 2 weeks post injection (wpi) (Extended Data Figs. 3d-f). Applying pseudotime and trajectory analysis resulted in the inference of six lineages: two astrocyte lineages (Lin1/Lin2), potentially reflecting astrocyte heterogeneity, consisting predominantly of control and Ascl1 (+/- Bcl2) expressing cells; two further lineages (Lin3/Lin4) progressing towards neurogenesis mostly comprised of cells expressing Ascl1 (+/- Bcl2); finally, Ascl1SA6 expressors gave rise to two further neuronal lineages (Lin5/Lin6), largely segregated by their expression of Bcl2 (Fig. 3f; Extended Data Fig. 4a). A striking hallmark of the segregation between the neuronal lineages was the early-onset expression of *Pvalb* in Ascl1SA6 expressors, particularly those co-expressing Bcl2 (Figs. 3g-i; Supplementary Table 6). While such early onset expression is surprising, detection of *Pvalb* transcripts in these cells is consistent with our previous report that a significant portion of Ascl1SA6/Bcl2 expressing iNs were immunoreactive for Parvalbumin protein at 4wpi ^21^.

To further evaluate the rate of neurogenesis induced by the different reprogramming conditions, we plotted progression of cells along pseudotime. This analysis suggested that cells progressed at a distinct pace towards their respective neuronal endpoints, with Ascl1SA6/Bcl2 expressors progressing at the fastest and Ascl1 expressors at the slowest pace. Intriguingly, many Ascl1 expressors were still characterized by astroglial gene expression, hinting at reprogramming roadblocks that were more easily overcome by the Ascl1SA6 expressors (Fig. 3j). Upon closer inspection of the RNA velocity profile, we identified two vector streams pointing away from neuronal endpoints, which mostly consisted of wildtype Ascl1 expressors. One vector stream direction headed towards an astrocyte fate, while the other pointed to an alternative endpoint (Fig. 3k; Extended Data Fig. 4b). Cells heading towards this alternative endpoint were enriched in expression of OPC-associated genes (*Olig1, Olig2, Pdgfra, Sox10*) ^34^, possibly indicating partial reprogramming of astrocytes towards an OPC fate (Fig. 3l; Extended Data Fig. 4c; DEGs, Supplementary Table 5a). Regarding those cells moving towards a neuronal endpoint, what accounts for the apparent lineage differences between wildtype- and phospho-site-deficient Ascl1 expressors? To our surprise, a main differentiator between neuronal cells expressing wildtype vs phospho-site-deficient Ascl1 was the lack of expression of Dlx gene family members such as Dlx1/2 in Ascl1SA6 expressors (Fig. 3m; Extended Data Fig. 4d), a difference that was confirmed by single molecule RNA fluorescence *in situ* hybridization (Fig. 3n,o; Supplementary Table 7). Thus, our single cell transcriptome data indicates that Ascl1SA6 drives induced neurogenesis more efficiently than Ascl1 via divergent trajectories, notwithstanding the puzzling fact that an integral module of the genesis of cortical interneurons (i.e., expression of Dlx genes) fails to become activated.

### Co-expression of Dlx2 induces canonical interneuron gene expression programs while suppressing Pvalb

Given that Ascl1SA6 (+/- Bcl2) failed to activate expression of Dlx gene family members, which are important upstream regulators of interneuron specification, we next asked whether co-expression of Dlx2 would promote a more canonical interneuron gene expression profile ^35^ (Fig.4a, Extended Data Fig. 4b). Indeed, at 4 days post-injection of retroviral vectors encoding Ascl1SA6 and Dlx2, most cells exhibited neuron-like morphologies (Fig. 4b), similar to those previously observed following Ascl1SA6/Bcl2 injection. However, single-cell RNAseq revealed that Ascl1SA6-Dlx2 co-expressors displayed marked differences in gene expression (Fig. 4a,c-d), including well-characterized Dlx2 transcriptional targets important in regulation of interneuron migration (e.g., *Zeb2, Arx*) as well as other members of the Dlx gene family ^36,37^ (Fig. 4d; Extended Data Fig. 5f). Additionally, while Ascl1SA6/Bcl2 and Ascl1SA6-Dlx2 expressors shared high levels of *Gad1* expression, *Gad2* transcripts were virtually exclusive to the latter (Fig. 4e). Moreover, consistent with its repressive role of Olig transcription factors ^38,39^, Ascl1SA6-Dlx2 co-expression showed a sharper decline of *Olig1* and *Olig2* transcripts in cells undergoing astrocyte-to-neuron conversion (Fig. 4f). This data supports the notion that Dlx2 expression supports Ascl1SA6 in the activation a key genes involved in interneuron specification and identity ^35^. Interestingly, in contrast to Ascl1SA6/Bcl2 expressors, we observed that Pvalb expression was virtually absent from Ascl1SA6-Dlx2 expressors (Fig. 4g; Extended Data Fig. 5g). Even when we expressed Bcl2 alongside Ascl1SA6 and Dlx2, *Pvalb* transcripts remained undetected (Fig. 4h), suggesting that the action of Dlx2 might delay or suppress Pvalb expression, or instruct distinct molecular programs. While this data shows that Dlx2 drastically reshapes the transcriptional landscape induced by Ascl1SA6, when expressed on its own, Dlx2 failed to induce neurogenesis despite inducing endogenous Ascl1 expression (Extended Data Figs. 5a-f; Supplementary Figs. 7).

**Figure 4.**
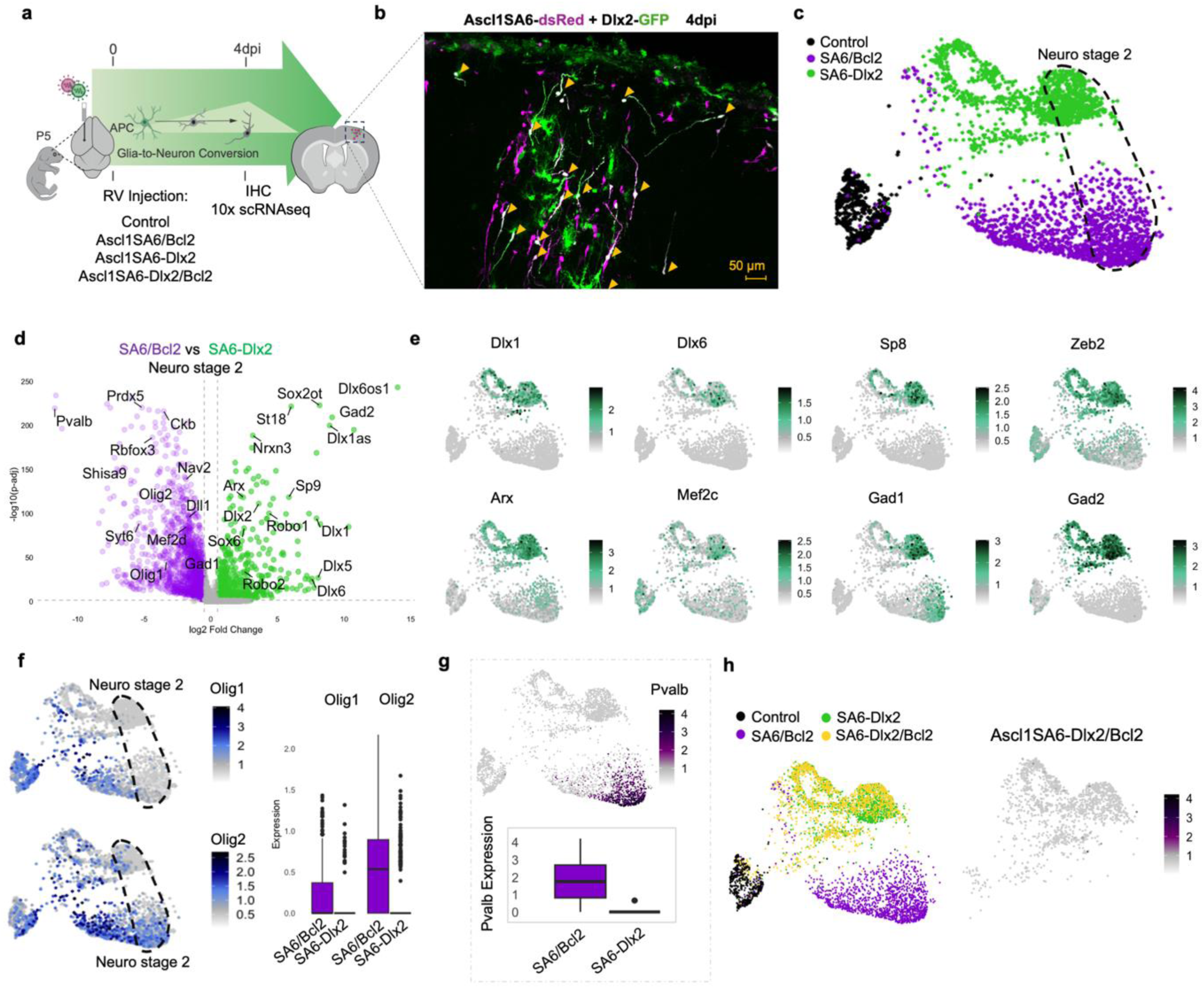
Co-expression of Dlx2 induces canonical interneuron gene expression programs while suppressing Pvalb. **a,** Experimental timeline: retroviral (RV) injection at day 0 (P5), followed by immunohistochemistry (IHC) and 10x Genomics single-cell RNA sequencing (scRNA-seq) at 4 days post-injection (dpi). **b,** Confocal images at 4 dpi showing co-transduced cells (magenta, *Ascl1SA6*; green, *Dlx2*; white, merge). Yellow arrows show cells expressing Ascl1SA6/Dlx2, exhibited early neuronal morphologies. **c,** UMAP embedding of 4383 single-cell transcriptomes (632 Control, 1,000 each for Ascl1SA6/Bcl2, and Ascl1SA6-Dlx2, down-sampled) colored by RV condition. The dotted line outlines Neuro stage 2 clusters. **d,** Volcano plot showing differentially expressed genes (FDR < 0.05) from a pseudo-bulk comparison between Ascl1SA6/Bcl2 Neuro stage 2 and Ascl1SA6-Dlx2 Neuro stage 2 populations. **e,** UMAP feature plots of genes in the transcriptional cascade downstream of Dlx2. **f,** UMAP feature plots (left) and Boxplots (right) display the distribution of normalized expression values for Olig1 and Olig2 across Neuro stage 2 cells in each condition (Control, Ascl1SA6/Bcl2, Ascl1SA6-Dlx2). The central horizontal line in each box represents the median expression. The box spans the interquartile range (IQR; 25th to 75th percentile), and whiskers extend to 1.5× the IQR. **g,** UMAP (top) and boxplots (bottom) showing *Pvalb* expression in each condition (Control, Ascl1SA6/Bcl2, Ascl1SA6-Dlx2). The central horizontal line in each box represents the median expression. The box spans the interquartile range (IQR; 25th to 75th percentile), and whiskers extend to 1.5× the IQR. **h,** UMAP embedding of 5,273 cells (632 Control, 1,370 Ascl1SA6/Bcl2, 1,447 Ascl1SA6-Dlx2, and 1,464 Ascl1SA6-Dlx2/Bcl2), colored by RV condition (left). UMAP feature plots showing *Pvalb* suppression in Ascl1SA6-Dlx2/Bcl2 (right).

### Induced neurons mature and acquire a CGE-like identity

Next, we sought to determine the transcriptional profiles of reprogrammed cells as they mature *in vivo*. To this end, we isolated high quality nuclei (36,960) at 2 and 4 wpi and performed single-nuclei RNA-sequencing (snRNA-seq) (Fig. 5a; Extended Data Fig. 1a; Supplementary Fig. 1b). Annotation of the resulting clusters indicated that besides the expected glial (i.e., astrocytes, OPCs, oligodendrocytes, microglia) and induced neuronal clusters, we could detect additional clusters comprising pericytes as well as endogenous neuron populations (Extended Data Fig. 6a,b; Supplementary Fig. 8, see also METHODS: Technical limitations of snRNAseq). Induced neuron transcriptomes formed a distinct cluster, were entirely absent from controls, and could only be detected in specific experimental groups, indicating their genuine emergence following reprogramming (Extended Data Fig. 6b).

**Figure 5.**
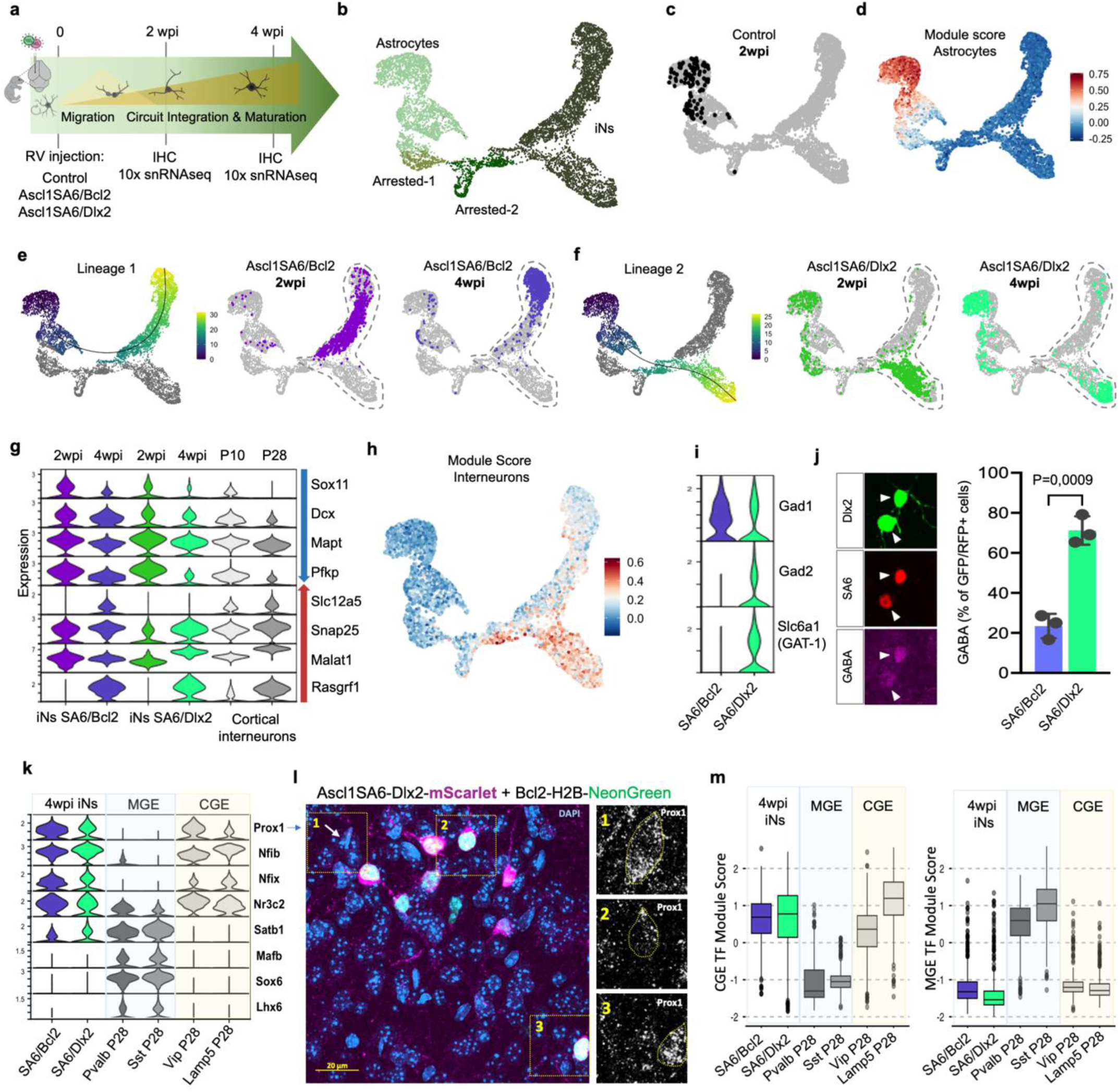
Induced neurons mature and acquire a CGE-like identity. **a,** Experimental workflow: retroviral (RV) injection at P5, followed by single-nucleus RNA sequencing (snRNA-seq, 10x Genomics) at 2- and 4-weeks post-injection (wpi). **b,** UMAP embedding of merged astrocyte-derived clusters colored by annotated cell type. **c,** Control cells in merged UMAP embedding. **d,** Module Score for cortical astrocytes ^34^. **e,f,** Separate pseudotime lineages and UMAP visualization of cells split by condition. Endpoints correspond to progressively maturing iNs at 4 wpi, dominated by Ascl1SA6/Bcl2 (**e**) and Ascl1SA6/Dlx2 (**f**) populations. **g,** Stacked violin plots show downregulation of immature neuronal genes (*Dcx, Mapt, Sox11, Pfkp*) and upregulation of maturation- and synapse-associated genes (*Slc12a5, Snap25, Malat1, Rasgrf1*) in comparison to endogenous interneurons at P10 and P28 ^41^. **h,** Module scores of P4 interneurons ^34^ plotted onto UMAP embedding. Data suggests Ascl1SA6/Dlx2 iNs align more closely with endogenous interneurons compared to Ascl1SA6/Bcl2 iNs. **i,** Expression of genes relevant to GABAergic neurotransmission: *Gad2* and *Slc6a1* are enriched in Ascl1SA6/Dlx2-iNs, whereas *Gad1* is expressed in both iN-populations. **j,** Confocal images showing GABA immunoreactivity (magenta) in Ascl1SA6/Dlx2-iNs at 4 wpi (left) and quantification of GABA+ transduced cells (right) (Ascl1SA6/Bcl2: 78 cells; Ascl1SA6/Dlx2: 171 cells; mean ± SD; n = 3 mice). Two-tailed unpaired Student’s t-test, ***P = 0.0009. **k,** Stacked violin plots of selected medial ganglionic eminence (MGE) vs caudal ganglionic eminence (CGE) transcription factors expressed in induced neurons compared to endogenous interneurons P28 ^41^ (left). UMAP feature plot of *Prox1* and *Lhx6* expression (right). **l,** left, Ascl1SA6-Dlx2-mScarlet + Bcl2-H2B-NeonGreen, DAPI in blue, mScarlet in magenta and merge is white right panels are maximum intensity projection single plane of magnification of 3 cells 1) endogenous Prox1+ cell 2) transduce cells with Ascl1SA6-Dlx2-mScarlet + Bcl2-H2B-NeonGreen with low Prox1 levels. 2) transduce cells with Ascl1SA6-Dlx2-mScarlet + Bcl2-H2B-NeonGreen with high Prox1 levels. 40x, white Prox1+ staining. **m,** Boxplots showing module scores generated from CGE (left) and MGE (right) transcription factors. The central horizontal line in each box represents the median expression. The box spans the interquartile range (IQR; 25th to 75th percentile), and whiskers extend to 1.5× the IQR.

Most iNs appeared to represent endpoints of transcriptional trajectories originating from astrocytes (Extended Data Fig. 6b), consistent with genetic fate mapping showing that over 70% of Ascl1SA6-Dlx2 iNs originated from astrocytic lineages (Extended Data Fig. 7a–c). In contrast, virtually none of the iNs could be traced back to an oligodendroglial origin (Extended Data Fig. 6b, 7d-f), despite evidence for early stages of OPC reprogramming at 4dpi (Fig. 1d). Intriguingly, a small but significant proportion of iNs appeared to have a pericyte origin (Extended Data Fig. 6b). However, we restricted our transcriptomic analysis to the major contributor of iNs, which were those derived from astrocytes (Fig. 5b-d). Pseudotime analysis indicated three lineages arising from astrocytes (two iN and one arrested/cycling lineage), clearly distinguished by the reprogramming factor combinations expressed in the cells along these lineages (Fig. 5e,f; Extended Data Fig. 8a). Importantly, most iNs (e.g., expressing *Rbfox3*) contained Ascl1SA6 in their reprogramming cocktail, while most Ascl1 expressors remained astroglial even when co-expressing Bcl2 (Fig. 5b-d, Extended Data Fig. 8b,c).

Intriguingly, Ascl1 and Dlx2 co-expressors failed to become iNs but remained in a seemingly arrested stage (Extended Data Fig. 8a-d). Thus, the main driver for successful astrocyte-to-neuron conversion in all our experiments is Ascl1SA6. Consistent with the differences we had observed already at 4dpi, Ascl1SA6/Bcl2 and Ascl1SA6/Dlx2 iNs gave rise to highly diverging lineages (Fig. 5e,f). Notably, iNs clearly matured over time as transcriptomes derived from 4-wpi nuclei accumulated at the pseudotime endpoints while most 2-wpi transcriptomes represented earlier stages (Fig. 5e,f). Accordingly, genes typically expressed by immature neurons and downregulated during neuronal maturation, such as *Sox11* and *Dcx*, were also downregulated in iNs between two to four weeks *in vivo*. Conversely, genes associated with neuronal maturation became upregulated (e.g., *Slc12a5*) (Fig. 5g; Extended Data Fig. 8e-h; Supplementary Fig. 9a). Remarkably, the Ascl1SA6/Dlx2 iNs exhibited a higher module score of developing interneurons (Fig. 5h; Extended Data Fig. 8i), in keeping with our earlier finding of expression of a more canonical interneuron program at 4dpi (Fig. 4d,e). These differences were highlighted by higher expression of *Gad2* and *Slc6a1* (GAT1) in Ascl1SA6/Dlx2 iNs, resulting in significantly higher GABA immunoreactivity as compared to Ascl1SA6/Bcl2 iNs (Fig. 5i,j).

Given that cortical interneurons originate from distinct germinal zones in the embryonic subpallium (medial ganglionic eminence, MGE vs caudal ganglionic eminence, CGE) ^40^, we next analyzed whether the transcription factor profiles of these iNs resemble more closely that of MGE- or CGE-derived interneurons. Intriguingly, both, Ascl1SA6/Bcl2 and Ascl1SA6/Dlx2 iNs displayed a transcription factor profile more akin to CGE-derived interneurons. This was underlined by the expression of *Prox1* and the conspicuous absence of *Lhx6* (Fig. 5k-m; Extended Data Figure 8j, k; Supplementary Fig. 9b,c; Supplementary Table 8).

### Ascl1SA6/Dlx2 iNs exhibit low-threshold spiking

Our previous work showed that a substantial portion of Ascl1SA6/Bcl2 iNs express hallmarks of fast-spiking interneurons (i.e., expression of parvalbumin and Kv3.1 protein as well as exhibiting action potential discharges above 100 Hz) ^21^. Given that co-expression of Dlx2 suppressed Pvalb, we next asked whether this homeobox transcription factor also prevented the expression of other hallmarks of fast-spiking interneurons. Indeed, we noted that while the gene encoding Kv3.1, critical for rapid hyperpolarization in fast-spiking interneurons, was significantly expressed in Ascl1SA6/Bcl2 expressors as early as 4dpi, Ascl1SA6/Dlx2 co-expressors transcribed markedly reduced levels of this potassium channel gene (Supplementary Fig. 10a). We next asked whether such early differences were reflected at later stages of iN maturation. Intriguingly, while Ascl1SA6/Bcl2 and Ascl1SA6/Dlx2 iNs transcribed many channel genes at similar levels, often similar to endogenous cortical interneurons ^41^, genes regulating key conductances in fast-spiking interneurons were expressed at significantly higher levels in Ascl1SA6/Bcl2 iNs as compared to Ascl1SA6/Dlx2 iNs (Fig. 6a,b; Extended Data Fig. 9a; Supplementary Table 9). Specifically, Ascl1SA6/Bcl2 iNs expressed significantly higher levels of *Scn1a* (Nav1.1), *Kcnc1* (Kv3.1), and *Hcn1* (HCN1), known to endow interneurons with fast-spiking capacity and highly expressed in FS/PV interneurons (Fig. 6a). Furthermore, these genes displayed maturation-dependent regulation in Ascl1SA6/Bcl2 iNs. In contrast, these genes were expressed at conspicuously low levels in Ascl1SA6/Dlx2 iNs (Fig. 6a; Extended Data Fig. 9a). Thus, we predicted that Ascl1SA6/Dlx2 iNs should not exhibit classical fast spiking characterized by rapid afterhyperpolarization. Indeed, when we performed current-clamp recordings from Ascl1SA6/Dlx2 iNs in acute brain slices, we noted drastically reduced afterhyperpolarization following action potential discharge and a trend towards lower firing frequencies (Fig. 6c-g; Extended Data Fig. b-d; Supplementary Fig. 10b; Supplementary Table 9). Likewise, consistent with the lower expression of *Scna1*, Ascl1SA6/Dlx2 iNs exhibited diminished action potential amplitudes as compared to Ascl1SA6/Bcl2 iNs (Extended Data Fig. b; Supplementary Table 9). In contrast, several of these iNs exhibited burst-spiking and rebound-spiking following hyperpolarization, suggesting that the electrophysiological properties of Ascl1SA6/Dlx2 iNs are determined to a significant degree by T-type calcium channel conductance (Fig. 6c-g; Extended Data Fig. 9b-d; Supplementary Fig. 10b). Indeed, *Cacna1g* encoding the T-type channel Cav3.1 was present in both iN populations, albeit at slightly higher levels in Ascl1SA6/Dlx2 iNs, and was down-regulated as iNs mature (Fig. 6b). This pattern of channel gene expression thus rendered Ascl1SA6/Dlx2 iNs more similar to low-threshold spiking interneurons ^42^.

**Figure 6.**
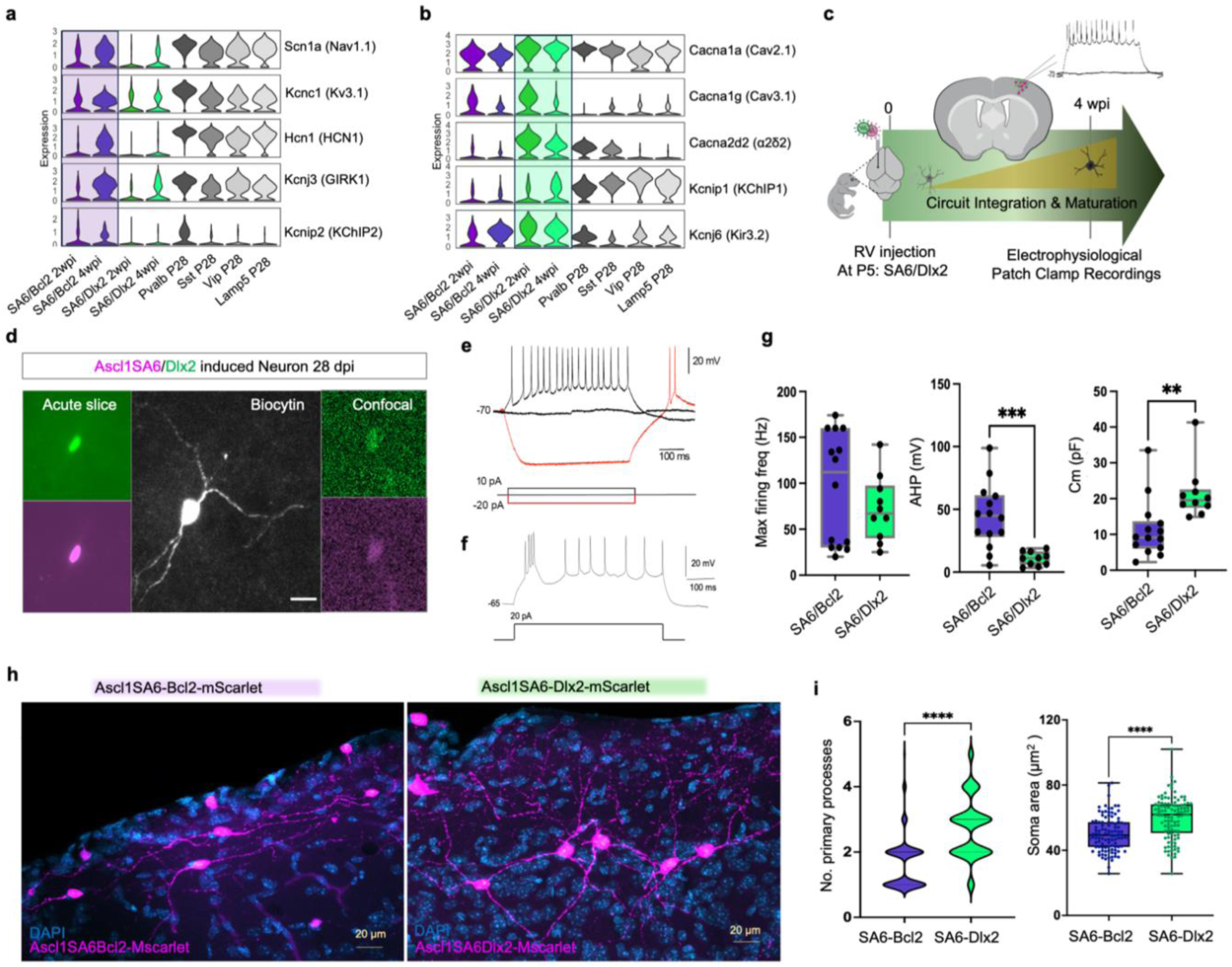
Ascl1SA6/Dlx2-induced neurons (iNs) exhibit low-threshold spiking and enhanced morphological maturation. **a,** Stacked violin plots of channels upregulated in Ascl1SA6/Bcl2-iNs at 2 and 4 weeks post-injection (wpi) compared to P28 endogenous cortical interneurons ^41^. **b,** Stacked violin plots of channels upregulated in Ascl1SA6/Dlx2-iNs at 2 and 4 wpi compared to P28 interneurons. **c,** Experimental timeline: retroviral (RV) injection at day 0, followed by whole-cell patch-clamp at 4 wpi. **d,** Left: Biocytin-filled Ascl1SA6/Dlx2-iN in acute slice. Insets: GFP (Dlx2) and mScarlet (Ascl1SA6) colocalization in acute slice (left) and after staining (right). Scale bar, 20 µm. **e,** Representative patch-clamp recording from Ascl1SA6/Dlx2-iN showing repetitive action potentials (10/10 cells) and rebound spiking after hyperpolarization (red; 5/10 cells). **f,** Bust firing and slow AHP Ascl1SA6/Dlx2-iN. **g,** Electrophysiological differences between Ascl1SA6/Bcl2 iNs and Ascl1SA6/Dlx2 iNs. Maximum firing frequency was not significantly different between the two groups (95 ± SEM Hz vs. 72 ± SEM Hz; p = 0.26). Bcl2 iNs showed significantly larger afterhyperpolarizations (AHPs; 44.7 ± SEM mV, n = 14 vs. 10.8 ± SEM, n = 10; p = 0.0002, Welch’s t-test); Membrane capacitance (Cm): Ascl1SA6/Dlx2 iNs exhibited higher Cm (21.5 ± SEM pF, n = 10 vs. 11.3 ± SEM pF, n = 14; p = 0.005. Values for Ascl1SA6/Bcl2 from ^21^. **h,** Confocal images at 4 wpi showing transduced cells in upper cortical layers (magenta: Ascl1SA6-Bcl2 or Ascl1SA6-Dlx2; blue: DAPI). Scale bar, 20 µm. **i,** Morphological parameters at 4 wpi. Dlx2 iNs displayed larger somata (82.96 ± 2.79 µm², n = 54 vs. 57.05 ± 1.94 µm², n = 80; p < 0.0001) and more primary processes (2.74 ± 0.11, n = 110 vs. 1.64 ± 0.05, n = 583; p < 0.0001). Data from 3 mice per group.

Similar to the divergence in channel gene expression, the two iNs populations transcribed genes for diverse classes of ionotropic and metabotropic glutamate and GABA receptors (Extended Data Fig. 9e). Consistent with this, Ascl1SA6/Dlx2 iNs (4/5) received spontaneous synaptic input as previously described for Ascl1SA6/Bcl2 iNs ^21^ (Extended Data Fig. 9f). Finally, while Ascl1SA6/Dlx2 iNs displayed similar input resistances as previously reported for Ascl1SA6/Bcl2 iNs, their membrane capacitance was significantly higher, indicative of an increased membrane surface (Fig. 6g). In line with this data, morphometric analysis revealed significantly larger somata of Ascl1SA6/Dlx2 iNs (Ascl1SA6/Dlx2: 82.96 ± 2.79 µm² vs. Ascl1SA6/Bcl2: 57.05 ± 1.94 µm²; p < 0.0001) and an increased number of primary processes (Ascl1SA6/Dlx2: 2.74 ± 0.11 vs. Ascl1SA6/Bcl2: 1.64 ± 0.05; p < 0.0001) (Fig. 6h,i; Supplementary Tables 10a-c).

### Dlx2 governs laminar position of induced neurons

In our previous work, we had noted that Ascl1SA6/Bcl2 iNs accumulated in layer 1, including those positive for parvalbumin, thus occupying partially atypical positions for cortical interneurons ^21^. Indeed, at 2 wpi 55.3% of Ascl1SA6-Bcl2 iNs were found to reside in layer 1, while only 22.9% localized to layers 2/3 (Fig. 7a-d; Supplementary Table 11), and they maintained such distribution over time (Fig. 7c, Extended Data Fig. 10a). While accumulation in layer 1 could be due to preferential retroviral targeting of glial cells in this layer, Ascl1SA6/Bcl2 iNs displayed swellings and branching of leading processes which are characteristic features of nucleokinesis of migrating interneurons (Fig. 7e). Thus, rather than being caused by a general motility defect, here we hypothesized that the absence of Dlx gene expression might account for their bias towards layer 1, and that co-expression of Dlx2 might promote a more canonical distribution (Fig. 7a). Indeed, early on Ascl1SA6/Bcl2 and Ascl1SA6/Dlx2 expressors differed in the expression of genes encoding transcriptional regulators of interneuronal migration programs (e.g., *Arx, Zeb2*) (Fig. 4d, e) as well as receptors of guidance cues (e.g., *Robo1* and *Robo2* among others) (Extended Data Fig. 10b) and these differences had persisted over time. Consistent with this differential gene expression, Ascl1SA6/Dlx2 co-expression induced a drastic shift towards layers 2/3, with only 26.3% remaining localized to layer 1 and 50.3% now populating layers 2/3 at 2wpi, and this laminar pattern of Ascl1SA6/Dlx2 iNs was retained by 4wpi (Fig. 7c, Extended Data Fig. 10a; Supplementary Table 11). Thus, Dlx2 co-expression alongside Ascl1SA6 does not only reshape the iN transcriptomic landscape towards more prototypical interneuronal gene expression but specifies their deployment within cortical circuits.

**Figure 7.**
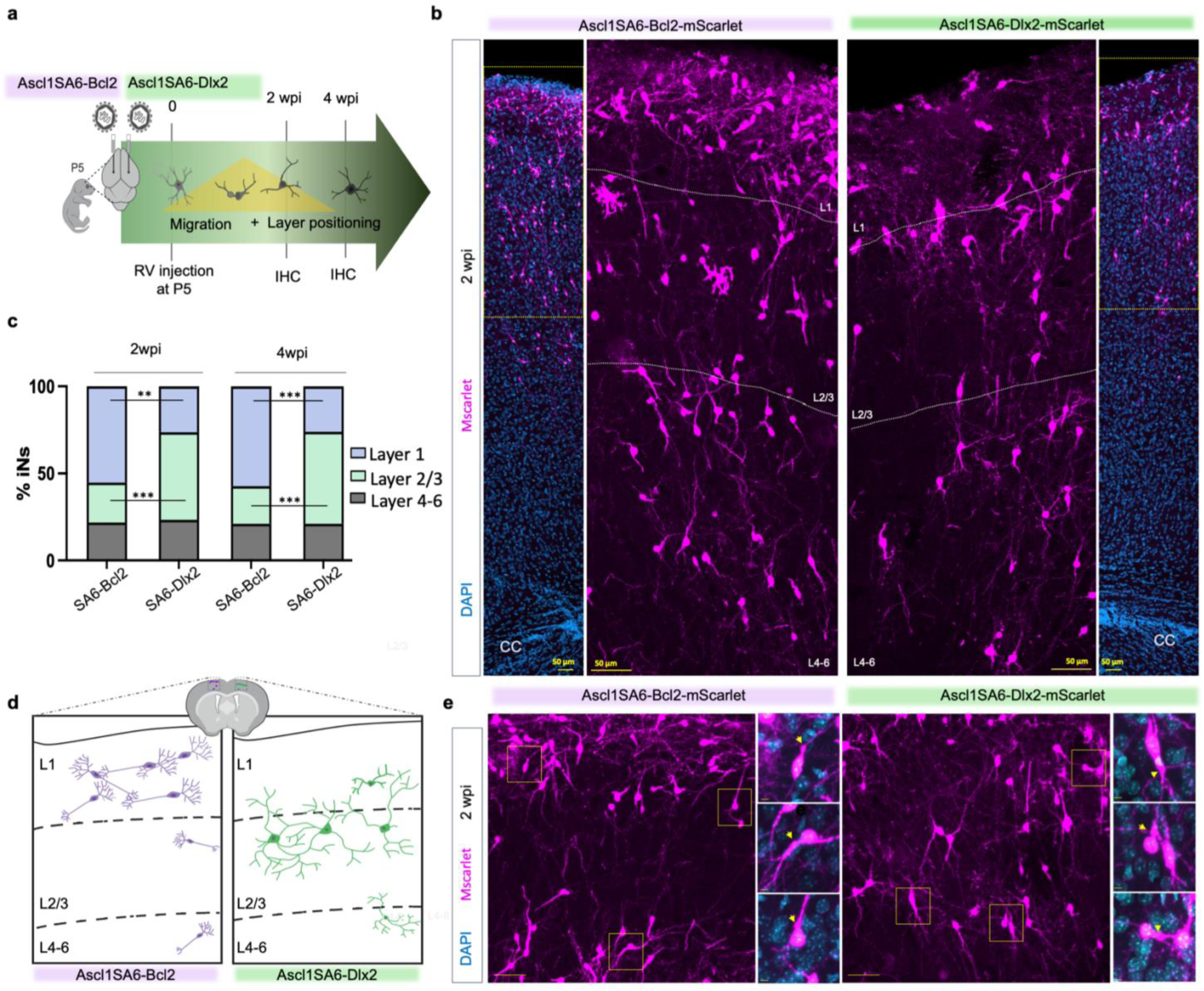
Dlx2 governs laminar positioning of induced neurons. **a,** Experimental workflow: retroviral (RV) injections at P5 and immunohistochemistry at 2 and 4 weeks post-injection (wpi). **b,** Distribution of Ascl1SA6-Bcl2-iNs (left) and Ascl1SA6-Dlx2-iNs (right) in the cerebral cortex at 2 wpi stained with DAPI (blue). Cortical layers are delineated by dotted lines (Layer 1, 2/3, 4–6**). c,** Stacked bar plots of laminar distribution for Ascl1SA6-Bcl2- and Ascl1SA6-Dlx2-iNs at 2 and 4 wpi. Data normalized per mouse.Ascl1SA6-Bcl2-iNs are significantly enriched in Layer 1 at 2 wpi (55.3% vs. 26.4%; t = 6.32, df = 4.98, p = 0.0015, **) and at 4 wpi (57.2% vs. 26.1%; t = 9.16, df = 4.54, p = 0.00043, ***). Conversely, Ascl1SA6-Dlx2-iNs preferentially localize to Layer 2/3 at 2 wpi (22.9% vs. 50.3%; t = –7.37, df = 5.00, p = 0.00073, ***) and at 4 wpi (21.7% vs. 52.8%; t = –9.66, df = 5.00, p = 0.00020, ***). No significant differences were observed in Layers 4-6 at either timepoint (21.8% vs. 23.3% at 2 wpi, t = –0.62, df = 4.69, p = 0.564, n.s.; 21.1% vs. 21.1% at 4 wpi, t = 0, df = 4.05, p = 1, n.s.). **d,** Schematic summarizing laminar positioning: Bcl2-iNs occupy Layer 1, while Dlx2-iNs are enriched in Layer 2/3. **e,** Confocal images of migrating neurons at 2 wpi, showing initiation of nucleokinesis marked by swelling and dynamic branching of leading processes. Representative Ascl1SA6-Bcl2-iNs (left) and Ascl1SA6-Dlx2-iNs (right). Insets (yellow) highlight swelling proximal to the nucleus.

## DISCUSSION

A key challenge to render *in vivo* glia-to-neuron reprogramming a viable approach for brain repair is to direct cells undergoing lineage conversion towards defined neuronal subtype and subclass identities. To this end, we aimed at uncovering the molecular logic that drives glia of the early postnatal mouse cortex towards acquiring interneuron identity. Based on our previous work demonstrating successful acquisition of subtype and subclass-specific interneuron hallmarks ^21^, here we set out to determine the transcriptional consequences of expressing the proneural gene Ascl1 and its phospho-site-deficient mutant Ascl1SA6 on postnatal cortical astrocytes and OPCs. Using single cell transcriptomics, we found that Ascl1SA6 not only clearly outperforms wildtype Ascl1 in inducing glia-to-neuron reprogramming, in line with observations that posttranslational modifications greatly impact Ascl1’s neurogenic activity ^21,23–26,43^, but strongly promotes the expression of a subset of genes that characterize specific interneuron subclasses such as *Pvalb* and *Kcnc1*. However, Click or tap here to enter text.Ascl1SA6 failed to induce Dlx gene family members, thus falling short in important aspects of forebrain interneuron specification ^35^. The addition of Dlx2 not only rendered astrocyte-derived transcriptomes more akin to authentic interneurons but also shifted iN distribution towards more canonical laminar locations within cortex ^46^. Thus, our work uncovers key aspects of the molecular logic underpinning lineage conversion towards endogenous neuronal identities.

Ascl1 has been shown to act as an on-target pioneer factor ^31,44^, thereby facilitating reprogramming through the opening of closed chromatin ^45,47^. However, pioneer factor-induced reprogramming often leads to the generation of cells that remain in an immature or hybrid state^48^. In line with this, we found that transcriptomes of Ascl1 expressors did not advance beyond immature neuronal stages or failed to exhibit clear signs of reprogramming altogether. In contrast, Ascl1SA6 was more effective in overcoming reprogramming hurdles and activating neurogenic programs in both astrocytes and OPCs. This data then indicates that wildtype Ascl1 undergoes phosphorylation in cortical glia ^25^, which may account for the widespread reprogramming failure in Ascl1 expressors. Recent work suggests that Ascl1SA6’ heightened neurogenic activity cannot be ascribed to enhanced stability alone, the strikingly different potency of Ascl1SA6 may be accounted for by additional mechanisms such as different binding to chromatin, more efficient nucleosome displacement, and/or differential recruitment of co-factors and chromatin remodelers ^26^.

In our experimental paradigm, i.e., uninjured early postnatal cortex, we noted a drastic difference in the induction of neurogenesis-related gene expression profiles of astrocytes vs OPCs. This was reflected by the fact that Ascl1SA6 induced more of Ascl1’s cognate neurogenic gene targets in astrocytes versus OPCs by 4 dpi. The temporal dynamics of Ascl1 target activation closely resembled those observed in ventral telencephalic NSCs, indicating that gene expression dynamics followed intrinsic mechanisms, possibly determined by chromatin remodeling ^31,44^. Moreover, at later time points, few if any OPC-derived iNs could be discerned, either in our transcriptome analyses or by genetic fate mapping ^21^. Lack of successful OPC reprogramming might arise from the context-dependent pioneering and transcriptional activity of Ascl1 ^26^, and/or from its activation of OPC-specific targets that may help maintain cells within the oligodendroglial lineage ^49–51^. Indeed, we observed that even some astrocytes partially activated oligodendroglial lineage genes in response to Ascl1. Furthermore, astrocytes may be more amenable to neuronal reprogramming, as they share the expression of genes, including transcription factors, with neurons generated within the same progenitor domains (Herrero-Navarro et al., 2021), and exhibit substantial injury-induced chromatin remodeling, including methylome changes, consistent with a transition toward an NSC-like state ^53^.

Here, we found that postnatal astrocytes expressed higher levels of *Sox2* and *Pax6* which may render them more reprogramming conducive, while OPCs were enriched in *Sox6* (Supplementary Fig. 6) which has been recently identified as an impediment to neuronal fate ^54,55^. Other proneural factors such as Neurog2 may be more effective than Ascl1 in activating neurogenic gene expression programs in OPCs ^56^. Finally, following lesion OPCs may remodel their chromatin to become more responsive to neurogenic cues ^5,10,54^.

Despite its enhanced neurogenic activity, Ascl1SA6 unexpectedly failed to induce expression of members of the Dlx gene family. The underlying mechanisms for this failure remain unclear. However, this failure has significant implications for iN specification. In fact, when co-expression of Dlx2 alongside Ascl1SA6 induced the full battery of interneuron differentiation promoting Dlx genes, resulting in an overall more prototypical program of interneuronal fate. This included key regulators of interneuron migration and maturation such as *Arx, Zeb2* and *Mef2c* as well as genes encoding the enzymes for GABA synthesis (*Gad1* and *Gad2*) ^35,57^. Consistent with its known roles, Dlx2 co-expression also promoted a more efficient down-regulation of *Olig1* and *Olig2* in line with their well-known mutually antagonistic roles ^39,58^, thereby helping to overcome reprogramming hurdles faced by iNs induced by Ascl1SA6. And yet, despite improving specification towards an interneuron identity, co-expression of Dlx2 totally abrogated *Pvalb* expression found in Ascl1SA6 iNs and resulted in a markedly reduced expression of *Kcnc1* (encoding Kv3.1). This raises the intriguing possibility that Dlx2 plays an active role in repressing the precocious induction of these genes, possibly being a contributing mechanism for the delayed surge of *Pvalb* in endogenous PV interneurons during their physiological maturation ^59^. Consistent with the lower expression of Kv3.1, Ascl1SA6/Dlx2 iNs showed a drastically reduced afterhyperpolarization as compared to iNs in the absence of Dlx2. Thus, in contrast to the latter, Ascl1SA6/Dlx2 iNs did not exhibit fast-spiking properties, instead their electrophysiological properties were dominated by the expression of T-type Ca^2+^ channels. Overall, Ascl1SA6/Dxl2 iNs often exhibited low-threshold burst spiking as well as rebound spiking when relieved from hyperpolarization, similar to what has been previously found in iNs induced in the epileptic hippocampus ^5^ following Ascl1 and Dlx2 co-expression and also found in a subclass of endogenous interneurons ^42,60,61^.

A particularly exciting observation was that co-expression of Dlx2 also resulted in iNs adopting a distinct pattern of distribution across cortical layers. Although iNs could be found in all layers, Ascl1SA6-Bcl2 iNs accumulated significantly in cortical layer 1, whereas their Dlx2 expressing counterparts became enriched in layers 2/3. Given that in general terms, their distribution appeared more similar to that of most endogenous cortical interneurons, this may suggest that Dlx2 regulates either migration, cell-cell interactions, or both in a more prototypical fashion. This is highlighted by differential expression of genes encoding transcription factors involved in regulation of migration as well as receptors involved in guiding cell migration as well as cell-to-cell signaling (e.g., ephrins, Eph receptors, Pcdh10, Lrps) ^62^. Thus, the acquisition of distinct laminar positions of iNs, dependent on the reprogramming factor cocktail employed, provides a handle to more rational approaches towards re-engineering diseased brain circuits via lineage reprogramming.

Finally, we also noted that both Ascl1 and Ascl1SA6 iNs were enriched for transcriptional regulators associated with CGE/LGE (caudal/lateral ganglionic eminence) rather than MGE (medial ganglionic eminence) origin. This distinction was particularly evident in the widespread expression of the CGE/LGE-associated gene *Prox1* ^63^ and the conspicuous absence of the MGE marker *Lhx6* ^64^. The prominent settling of iNs in layers 1 as well as 2/3 may be reflection of this more CGE-like identity ^65,66^. Future studies may address whether iNs can be steered to adopt more MGE-like identities.

In conclusion, our findings highlight the importance of uncovering the molecular logic and the associated critical reprogramming bottlenecks that drive lineage reprogramming of cortical glia toward neurons with subtype- and subclass-specific hallmarks, thereby paving the way for strategies to restore inhibitory circuits in neurological diseases.

## METHODS

### Plasmids and retroviruses

MMLV-based retroviral vectors were generated as previously described (Heinrich et al., 2011; Marichal et al., 2024). Briefly, coding sequences for Ascl1 and Ascl1SA6 were excised from pCIG2-Ascl1 and pCIG2-Ascl1SA6 parental vectors (Li et al., 2014) and subcloned into Gateway-compatible entry vectors containing attL recombination sites. These sequences were recombined into CAG promoter–driven retroviral backbones to generate pCAG-Ascl1-IRES-dsRed, pCAG-Ascl1SA6-IRES-dsRed, and a corresponding control vector pCAG-dsRed. For co-expression of the anti-apoptotic factor Bcl2, the previously described pMIG-Bcl2-IRES-GFP construct was used ^16^. The coding sequence for Dlx2 was obtained from published work ^5^ and inserted into the same retroviral backbone to generate pCAG-Dlx2-IRES-GFP. To enable bicistronic expression of transcription and survival factors, additional constructs were engineered linking factors via a self-cleaving T2A peptide. To facilitate visualization of transgene expression across experimental paradigms, we generated constructs expressing distinct fluorescent reporters either localized to the cytoplasm (non-nuclear) or fused to histone H2B for nuclear expression. In addition to Gateway Recombination Cloning, we PCR amplified relevant fragments and used NEBuilder to clone into a retroviral backbone plasmid. The sequences of all the resultant plasmids are in Supplementary Figure 11. The non-nuclear (cytoplasmic) constructs using Gateway cloning included pCAG-control-IRES-dsRed, pCAG-Ascl1-IRES-dsRed, pCAG-Ascl1SA6-IRES-dsRed, pMIG-Bcl2-IRES-GFP, pCAG-Dlx2-IRES-GFP, and pCAG-Ascl1SA6-T2A-Dlx2-IRES-dsRed, whereas non-nuclear constructs generated via the NEBuilder protocol included pCAG-Ascl1SA6-IRES-mScarlet, pCAG-Ascl1SA6-T2A-Bcl2-IRES-mScarlet, pCAG-Ascl1SA6-T2A-Dlx2*-IRES-mScarlet (Dlx2 plasmid was used from Addgene and pCAG-control-IRES-Mscarlet). Nuclear (H2B-tagged) constructs generated via the NEBuilder protocol included pCAG-control-H2B-Mscarlet, pCAG-Ascl1-H2B-Mscarlet, pCAG-Ascl1SA6-H2B-Mscarlet, pMIG-Bcl2-H2B-GFP, and pCAG-Dlx2-H2B-NeonGreen. Retroviral particles were produced using gpg helper-free packaging cells to generate VSV-G–pseudotyped retroviral particles ^67^. Viral supernatants were collected 48–72 h post-transfection, filtered through 0.45 μm membranes, and concentrated by ultracentrifugation. Pellets were resuspended in tris-buffered saline (TBS) and stored at −80°C until use. Viral titers for all preparations ranged between 10^7 and 10^9 transducing units/mL. All constructs were sequence-verified and tested for expression in HEK293T cells prior to viral production. We use separate and combine viral combinations for example Ascl1SA6/Dlx2 means separate viruses were injected and Ascl1SA6-Dlx2 implies both Ascl1SA6 and Dlx2 are combined in one plasmid vector. Find DNA sequences for plasmids in Supplementary Figure 12.

### Animals and animal procedures

All animal procedures were conducted in accordance with the European Directive 2010/63/EU on the protection of animals used for scientific purposes and the UK Animals (Scientific Procedures) Act (1986). Experimental protocols were approved by the Ethical Review Committee of King’s College London and the UK Home Office (project license numbers PD025E9BC and PP8849003; establishment license X24D82DFF). This project adhered to the UKRI principles of Responsible Research and Innovation (RRI). Mice were maintained on a 12:12 h light–dark cycle with food and water available ad libitum. Male and female C57BL/6J pups bred in-house from adult mice purchased from Charles River Laboratories (Walden, UK). Transgenic mouse lines were generated in-house. For this purpose, mice carrying a tamoxifen-inducible Cre recombinase under the Aldh1l1 locus (Aldh1l1-Cre/ERT2; JAX#031008; Srinivasan et al., 2006) or the NG2 promoter (NG2-CreERTM; JAX#008538; Zhu et al., 2011), were crossed with an EGFP reporter line (RCE:loxP; JAX#032037; Sousa et al., 2009) to generate double transgenic animals (mGFAP-Cre/RCE:loxP, Aldh1l1-Cre/ERT2/RCE:loxP, or NG2-CreERTM/RCE:loxP). In Aldh1l1-Cre/ERT2 and NG2-CreERTM/RCE:loxP mice, Cre recombination was induced by subcutaneous administration of tamoxifen (ApexBio Technology, #B5965) dissolved at 6 mg/mL in corn oil (Sigma-Aldrich, #C8267) with 10% ethanol. Pups received 60 μL per day from P2–P5 (Aldh1l1-Cre/ERT2) or on P2 and P5 (NG2-CreERTM). Retroviral injections targeting the somatosensory and visual cortical areas, either one or both hemispheres, were performed as described previously ^50^. Briefly, P5 pups were anesthetized, and 1 μL of viral suspension was injected using glass capillaries through a small skull incision. Injection coordinates relative to lambda were: +3 mm rostrocaudal, ±0.5 mm mediolateral, and −0.5 mm ventral. After injection, wounds were sealed with surgical glue (3M Vetbond, NC0304169), and pups were allowed to recover on a 37°C warming plate or chamber before being returned to the dam. Postoperative recovery was monitored daily for one week following surgery.

### Sample and Library Preparation for scRNA-seq and snRNA-seq

Single-cell RNA sequencing (scRNA-seq): Following viral injection, brains were collected 4 days post-injection (dpi). Brains were rapidly removed and placed in a mouse brain matrix, then sectioned into 0.5-mm coronal intervals. Cortical tissue proximal to the injection site was carefully dissected under a stereomicroscope. Single-cell suspensions were generated using the Adult Brain Dissociation Kit (Miltenyi Biotec, 130-107-677) and processed on a gentleMACS™ Octo Dissociator according to the manufacturer’s instructions and as described^28,29^.

Single-nucleus RNA sequencing (snRNA-seq): For nuclear transcriptomic profiling, injected brains were collected at 2 and 4 weeks post-injection (wpi). Single-nucleus suspensions were prepared following the 10x Genomics protocol: Nuclei Isolation from Adult Mouse Brain Tissue for Single Cell RNA Sequencing.

Fluorescence-Activated Cell Sorting (FACS): Fluorophore-positive cells or nuclei (GFP⁺/RFP⁺/NeonGreen/Mscarlet, either single or double positive) were isolated by FACS using a BD FACSAria™ III Cell Sorter operated with BD FACSDiva™ Software v6.1.3. Sorting was performed with a 100-μm nozzle for whole cells and a 70-μm nozzle for nuclei. Post-sort, cell and nuclei suspensions underwent quality control (QC) using the LUNA-FX7™ Automated Cell and Nuclei Counter to assess viability, concentration, and purity.

Library construction and sequencing: High-quality single-cell and single-nucleus suspensions were loaded onto the 10x Chromium Controller / Chromium X (10x Genomics) using Chromium GEM-X Single Cell 3′ Gene Expression kits, either v3 (PN-1000075) or v4 (PN-1000691), following the manufacturer’s user guide (Chromium Single Cell 3′ Reagents Kits v3.1/v4.1). cDNA synthesis and library preparation were carried out according to the 10x Genomics workflow. Libraries were quantified using an Agilent BioAnalyzer and pooled for sequencing on an Illumina NovaSeq X Plus or NovaSeq 6000 platform at the Francis Crick Institute (London, UK), targeting a minimum sequencing depth of 50,000 reads per cell. Raw sequencing data were processed using Cell Ranger (10x Genomics, Software version cellranger-7-9.0.0) for demultiplexing, alignment, and gene expression quantification. Workflow depicted in Extended data Figure 1a.

In addition to the data generated for this study, we incorporated previously published P10 and P28 endogenous interneuron datasets. We integrated data from two single-cell RNA-sequencing experiments of the ALM at these developmental stages conducted by Allaway and colleagues (GEO: GSE165233; sample accessions GSM5014305 and GSM5014307 ^41^). For downstream differential expression analysis, the raw data from this publication were processed and re-run through Cell Ranger under the same conditions as our single-nuclei RNA-sequencing pipeline. Within the integrated dataset, the four endogenous interneuron subtypes were annotated based on canonical marker expression: Pvalb, Sst, Vip, and Lamp5.

### Bioinformatic analyses of single-cell and single-nuclei RNA-sequencing

#### Quality control

Gene expression matrices were imported into R (v4.4.1) and processed using Seurat v5.3.0. Default parameters were applied unless otherwise specified. Each dataset was processed individually. Genes expressed in fewer than four cells were excluded, and cells with fewer than 200 detected genes were removed. Low-quality cells with high mitochondrial gene content were identified and filtered using the miQC-wrapper function, RunMiQC (SeuratWrappers package, v0.3.2). Putative doublets were detected with scDblFinder v1.18.0 and excluded. Cells with total gene counts exceeding five median absolute deviations (MADs) from the dataset median were also removed.

Filtered count matrices were log-normalized using NormalizeData and scaled with ScaleData. The top 2000 variable genes were identified using FindVariableFeatures, and linear dimensionality reduction was performed by principal component analysis (PCA) on these genes (RunPCA). To identify outliers, cells were clustered at low resolution (resolution = 0.1; FindClusters) using a shared nearest neighbor graph (FindNeighbors) on the first 50 principal components (PCs), and the Mahalanobis distance (stats package, v4.4.1) of each cell from its cluster centroid was calculated on the first 10 PCs. A 90% Chi-squared confidence threshold was applied to flag outliers, which were removed except for cells expressing reporter genes. Following outlier removal, variable feature selection, scaling, and PCA were recomputed to generate the final datasets for downstream analyses.

#### Data Processing

Datasets corresponding to the same sequencing strategy (scRNA-seq, 4 DPI; snRNA-seq, 2 WPI and 4 WPI) were merged into a single Seurat object, with data layers combined to treat the merged datasets as a unified dataset. The top 3,000 variable features of the merged datasets were identified, scaled, and used for principal component analysis (PCA). A nearest-neighbor graph was constructed using 25 PCs for the 4 DPI dataset and 20 PCs for the 2 WPI and 4 WPI datasets. Clusters were identified with a resolution of 3 (4 DPI) and 0.5 (2 WPI and 4 WPI), and two-dimensional UMAP embeddings were generated using 21 PCs (4 DPI) and 23 PCs (2 WPI and 4 WPI). Differential gene expression among clusters was assessed using the FindAllMarkers function, with thresholds of log₂ fold change > 0.25, detection in > 25% of cells, and adjusted p < 0.05. Clusters were annotated based on canonical marker expression.

#### Module Score

To quantify per-cell activity of defined gene sets, module scores were calculated using the AddModuleScore function (Seurat v4.3.0, R v4.2.3), with 5 control genes from the same expression bin for each gene in the module. Gene modules were defined based on curated literature lists of transcription factors and marker genes relevant to the cell types analyzed. Mouse MGE/CGE transcription factors were identified using differentially expressed genes (DEGs) from CGE and MGE P28 interneurons from the dataset by Allaway et al., 2021 ^41^, and cross-referenced with a curated list of mouse transcription factors available at https://esbl.nhlbi.nih.gov/Databases/KSBP2/Targets/Lists/TranscriptionFactors (accessed Feb 2025). All analyses were performed on the mm10 mouse genome assembly.

#### RNA Velocity

To determine the trajectory direction of cells during reprogramming, we employed RNA velocity. Seurat objects were converted into AnnData objects (H5AD format) using SeuratDisk (v0.0.0.9021) and downstream analyses were performed in a Python environment (3.14.2). Spliced/unspliced expression matrices were generated from Cell Ranger output BAM files using the Velocyto package (v0.17.17). The scVelo package (v0.3.3) was used to estimate RNA velocity vectors: first- and second-order moments were computed using 30 nearest neighbors and 30 PCs, velocities were estimated employing stochastic modeling, and a velocity graph containing cell transition probabilities was generated. To identify cells heading “towards alternative fate,” we selected cells deviating by more than 10% from the trajectory towards neuronal fate, specifically within clusters corresponding to intermediate and neuronal stages 1 and 2.

#### Lineage analysis

To delineate cellular trajectories and infer pseudotemporal dynamics within our reprogramming paradigm, we employed the Slingshot toolkit (v2.16.0). Cycling cells, identified via the RNA Velocity latent time root cell computation, were designated as the root population for trajectory inference. Lineage relationships were reconstructed using the principal graph method implemented in Slingshot, enabling cells to be ordered along continuous pseudotemporal axes. Branch points were delineated to interrogate potential lineage bifurcations and cell fate decisions. The resulting trajectory data, encompassing pseudotime values and principal graph embeddings, were extracted and exported for downstream analyses. Slingshot fitted a principal curve through the high-dimensional data, effectively capturing the progressive maturation and differentiation dynamics of the cells along the inferred trajectories.

#### GO analysis

Gene Ontology (GO) analysis of biological processes was performed on the differentially expressed genes for each induced neuron subtype, including both up- and downregulated genes, using the R package clusterProfiler (v4.16.0) in R v4.2.3. Analyses were corrected for multiple testing using the Benjamini-Hochberg false discovery rate (FDR) and restricted to the “GO:BP” category of the Gene Ontology database. For each analysis, active genes, defined as genes expressed in more than 25% of cells, consistent with the threshold used for differential expression, were used as the background gene set. GO term categories presented in Figs. 3c and Extended Data Fig. 10a were selected manually based on biological relevance. No additional statistical cutoff was applied beyond the FDR threshold for differential expression.

#### Technical limitations

Our single-nuclei RNA-seq dataset revealed additional clusters beyond the expected glial and pericyte populations, including endogenous neurons (both excitatory and inhibitory) (Extended Data Fig. 6a,b; Supplementary Fig. 8). The presence of these neurons was unexpected, as no endogenous neurons were detected at 4 dpi (Fig. 1d), likely reflecting incomplete FACS selectivity and contamination by unlabeled nuclei. Supporting this, reporter gene (mScarlet) transcripts were rarely detected in these neurons, in contrast to nuclei of induced neurons (Extended Data Fig. 6c), and transcriptomes from control and reprogramming factor-injected brains largely overlapped (Extended Data Fig. 6b). These observations do not impact the interpretation of iN-specific results but underscore the technical challenges associated with isolating pure reprogrammed populations for high-throughput single-cell transcriptomic analysis.

### Immunohistochemistry

Tissue preparation and immunostaining were performed as previously described ^21^. Briefly, animals were lethally anesthetized by intraperitoneal administration of ketamine (120 mg/kg; Zoetis) and medetomidine (1 mg/kg; Orion Pharma) prepared in 0.9% NaCl, and transcardially perfused with 0.9% NaCl followed by 4% paraformaldehyde (PFA; Sigma-Aldrich, Merck, Germany, P6148). Brains were harvested and postfixed overnight in 4% PFA at 4°C. Coronal sections of 40 μm thickness were prepared using a vibratome (Microm HM650V, Thermo Fisher Scientific, or Leica VT1000S) and stored at −20°C in a cryoprotective solution containing 20% glucose, 40% ethylene glycol, and 0.025% sodium azide. Using a free-floating procedure, sections were washed three times for 15 min with 1× phosphate-buffer saline (PBS) (50 mM Tris, 150 mM NaCl, pH 7.6) and incubated for 1.5 h in blocking solution containing 2.5% donkey serum, 2.5% goat serum, 0.3% Triton X-100, and 1× PBS. Sections were then incubated with primary antibodies diluted in blocking solution for 2–3 h at room temperature (RT) followed by overnight incubation at 4°C. After three washes with 1× PBS, sections were incubated with secondary antibodies diluted in blocking solution for 2 h at RT, washed once with 1× PBS, incubated with 5 μM DAPI in 1× PBS for 5 min at RT, and washed three times with 1× PBS. For mounting, sections were washed twice with 1× phosphate buffer (PB) (30 mM Na₂HPO₄·12H₂O, 33 mM NaH₂PO₄·2H₂O, pH 7.4), dried on 1mm thick microscope slides, and covered with coverslips using Mowiol supplemented with DABCO. The following primary antibodies were used: anti-Dcx (goat, 1:250, sc-8066), anti-Dcx (guinea pig, 1:500, AB2253), anti-GABA (rabbit, 1:300, A2052), anti-GFP (chicken, 1:1000, GFP-1020), anti-GFP (goat, 1:500, ab5450), anti-mCherry (chicken, 1:300, CPCA-mCherry), anti-NeuN (mouse IgG1, 1:500, MAB377), anti-RFP (rabbit, 1:500, 600401379S), anti-Sox11 (rabbit, 1:300, ab134107). anti-Lhx6 (mouse, 1:200, sc-271433), anti-PROX1 (goat, 1:200, AF2727). Secondary antibodies were made in donkey or goat and conjugated with the following: A405 (anti-mouse, 1:250, 715-475-151), A488 (anti-chicken, 1:500, 703-545-155), A488 (anti-goat, 1:200, ab150129), A488 (anti-mouse, 1:200, A21202), A568 (anti-rabbit, 1:500, A11011), A647 (anti-rabbit, 1:500, A31573), A647 (anti-mouse, 1:500, A31571), Cy3 (anti-chicken, 1:500, 703-165-155), Cy3 (anti-rabbit, 1:500, 711-165-152), Cy5 (anti-goat, 1:500, 705-175-147), Cy5 (anti-guinea pig, 1:500, 706-175-148), and A647 (anti-rat, 1:500, ab150155).

### 5-Ethynyl-2′-deoxyuridine incorporation assay

Two doses of EdU (ThermoFisher Scientific, A10044) were administered by intraperitoneal injection at 50 mg/kg (in 0.9% NaCl and 0.25% DMSO), 6 hours apart, 4 days after retrovirus delivery. Analysis was performed 2 days post-injection. Immunohistochemistry was performed as described above. After secondary antibody incubation, EdU incorporation was detected using the Click-iT™ EdU Cell Proliferation Kit for Imaging, Alexa Fluor™ 647 dye (Invitrogen, C10340). Slices were then incubated with 5 μM DAPI in 1× PBS for 5 min at room temperature and washed three times with 1× PBS. For mounting, sections were washed twice with 1× phosphate buffer (30 mM Na₂HPO₄·12H₂O, 33 mM NaH₂PO₄·2H₂O, pH 7.4), dried on 1 mm thick microscope slides, and coverslipped using Mowiol supplemented with DABCO. Primary antibodies used were anti-GFP (chicken, 1:1000, GFP-1020) and anti-RFP (rabbit, 1:500, 600401379S), with secondary antibodies conjugated to Alexa Fluor 488 (anti-chicken, 1:500, 703-545-155) and Alexa Fluor 568 (anti-rabbit, 1:500, A11011).

### Single-molecule fluorescence in situ hybridization (smFISH)

Single-molecule fluorescence in situ hybridization (smFISH) was performed using the RNAscope Multiplex Fluorescent V2 kit (Advanced Cell Diagnostics, ACD) in accordance with the manufacturer’s instructions. All solutions were prepared in ribonuclease-free diethyl pyrocarbonate (DEPC)–treated water (Sigma-Aldrich, Merck, Germany; D5758). Hybridizations were carried out with target-specific probes against Pvalb-C1 (#421931) and Dlx2-C3 (#555951). Control hybridizations were performed using a positive control probe (PN320881) and a negative control probe (PN320871). Tissue sections were placed in 0.05 M PBS and mounted in 0.1 M PB on SuperFrost Plus slides (J1800AMNZ, Menzel). Sections were air-dried overnight at room temperature (RT), followed by incubation at 40°C for 40 min in a HybEZ II oven (ACD). After rinsing with distilled water (dH₂O), sections were dehydrated sequentially in 50%, 70%, and 100% ethanol (EtOH) for 5 min each. Samples were then treated with hydrogen peroxide (RNAscope V2 kit, ACD) for 10 min at RT, washed three times (3–5 min each) in dH₂O. Antigen retrieval was performed using the RNAscope antigen retrieval solution (Multiplex Fluorescent V2 kit, ACD) at 90°C for 10–15 min, after which sections were rinsed in dH₂O, briefly dipped in 100% EtOH, and air-dried for approximately 5 min. Subsequently, sections were incubated with Protease III (RNAscope V2 kit, ACD) for 10–15 min at 40°C in a HybEZ II oven. After three washes in dH₂O (3–5 min each), hybridization was performed with the target probes for 2 hours at 40°C in the HybEZ II oven. Sections were then washed three times in RNAscope wash buffer (3–5 min each) and stored overnight in 5× saline-sodium citrate (SSC) buffer prepared according to ACD’s user manual. The following day, signal amplification was carried out sequentially with AMP1 (30 min, 40°C), AMP2 (30 min, 40°C), and AMP3 (15 min, 40°C) reagents (Multiplex Fluorescent V2 kit, ACD), with intermediate washes in wash buffer (3×, 3–5 min each). Fluorescent signal development was performed by incubation with horseradish peroxidase (HRP)–C1 or HRP–C3 (corresponding to the probe channel) for 15 min at 40°C, followed by labeling with Opal 690 dye (FP1497A, ACD) or Opal 570 (FP1488A, ACD) at 1:1000 dilution in Tyramide Signal Amplification buffer (#322809, ACD) for 30 min at 40°C, and subsequent incubation with HRP blocker (15 min at 40°C). All steps were performed in the HybEZ II oven with three intermediate washes in wash buffer (3–5 min each). Following hybridization, sections were counterstained with 5 μM DAPI in 1× TBS for 7 min at RT, rinsed in 0.1 M TBS, and processed for immunofluorescence. For this, sections were permeabilized in PBS containing 0.25% Triton X-100 for 20 min at RT, followed by blocking for 2 hours at RT in 0.3% Triton X-100, 3% bovine serum albumin (BSA; Scientific Laboratory Supplies, #A2153), and 10% normal donkey or goat serum (or 5% donkey and 5% goat serum) in 1× PBS. Primary antibodies were diluted in blocking buffer (0.3% Triton X-100, 3% BSA, 5% serum in PBS) and incubated for overnight + 24 hours at 4°C. After three washes in 1× PBS, sections were incubated with secondary antibodies for 2 hours at RT, rinsed, and mounted in ProLong Gold Antifade Mountant (Thermo Fisher Scientific, Waltham, MA, USA; P36930). The following antibodies were used: Primary antibodies: chicken anti-GFP (1:200) and rabbit anti-RFP (1:100). Secondary antibodies: donkey or goat anti-chicken Alexa Fluor 488 (1:200) and anti-rabbit Alexa Fluor 568 (1:250).

### Confocal microscopy and quantifications

Immunostainings and RNAscope signals were acquired using a Zeiss LSM 800 confocal microscope (Carl Zeiss Microscopy, Jena, Germany; Centre for Developmental Neurobiology, King’s College London) equipped with four solid-state lasers (405, 488, 561, and 633 nm) and 20× (NA 0.8) or 40× (NA 1.3) objectives. Serial optical Z-stacks were acquired at 0.3–2.13 μm intervals to capture the full thickness of the brain sections. Image acquisition was carried out using Zeiss ZEN software. For figure preparation, maximum intensity projections of the Z-stacks were generated using the respective software functions. Cell quantification was performed using ZEN or ImageJ v1.51v (National Institutes of Health, USA). Cell identification and counting were conducted by navigating through Z-stacks to ensure accurate visualization of transduced cells. For fate-mapping analyses, the following populations were quantified: (i) RFP/EGFP/Dcx or RFP/EGFP/NeuN triple-positive cells (fate-mapped induced neurons, iNs); (ii) RFP/Dcx or RFP/NeuN double-positive iNs; (iii) RFP/EGFP double-positive transduced cells; and (iv) RFP-only transduced cells. For Sox11 expression analyses RFP/Sox11 positive cells were quantified. The proportion of each population was expressed as a percentage of the total number of reporter- or double-reporter–positive transduced cells. Data are presented as mean ± SD, with quantifications performed from at least three to five brain sections per animal, across three independent biological replicates (minimum of nine sections in total). For RNAscope analyses, DsRed reporter fluorescence (transduced cells) or DAPI+ nuclei (endogenous neurons) were used to delineate regions of interest (ROIs). The number of mRNA transcripts was manually quantified in maximum projection images when puncta were clearly distinguishable or navigating through the Z-stack otherwise. Transcript counts were expressed as the total number of puncta per DsRed+ cell or endogenous neuron. Quantifications were performed on cells from three independent mice for each experiment.

### Electrophysiological recordings

#### Slice Preparation

Mice were anesthetized with isoflurane (Forane, AbbVie, IL, USA) and decapitated. Brains were rapidly removed and placed into ice-cold artificial cerebrospinal fluid (ACSF) containing (in mM): NaCl, 85; sucrose, 73; KCl, 2.5; NaHCO₃, 25; CaCl₂, 0.5; MgCl₂, 7; NaH₂PO₄, 1.25; and glucose, 10. The ACSF was continuously bubbled with 95% O₂ / 5% CO₂ to maintain pH 7.4. Coronal cortical slices (300 μm thick) were cut using a vibratome (VT1200 S, Leica Microsystems, Wetzlar, Germany) and transferred for recovery to standard ACSF (34°C, 10–15 min) composed of (in mM): NaCl, 125; KCl, 2.5; NaHCO₃, 25; CaCl₂, 2; MgCl₂, 1; NaH₂PO₄, 1.25; and glucose, 12 (pH 7.4). Slices were subsequently maintained in standard ACSF at room temperature (21 ± 2°C) for at least 1 hour prior to recording. For recordings, individual slices were transferred to a submerged recording chamber, continuously superfused (1–2 ml·min⁻¹) with standard ACSF, and mounted on an upright microscope (SliceScope Pro 6000 System, Scientifica, Uckfield, UK). Cells were visualized using Dodt gradient contrast optics with a 40× water immersion objective (NA 0.70) and a Hamamatsu Orca-Flash 4.0 camera (Hamamatsu, Japan). Retrovirally transduced cells expressing GFP or DsRed were identified via LED fluorescence excitation (CoolLED pE-100, Andover, UK) using appropriate excitation and emission filters.

#### Electrophysiology

Whole-cell patch-clamp recordings were performed at 30 ± 2°C using an in-line solution heater (Scientifica, Uckfield, UK). Recording pipettes (10–15 MΩ) were fabricated from borosilicate glass capillaries (1B150F-4, World Precision Instruments, Sarasota, FL, USA) using a horizontal micropipette puller (P-1000, Sutter Instruments, Novato, CA, USA). The pipette solution contained (in mM): K-gluconate, 125; NaCl, 5; Na₂ATP, 2; MgCl₂, 2; EGTA, 1; HEPES, 10; and biocytin, 10 (for post hoc morphological analysis); adjusted to pH 7.4 and 280 mOsm. Voltage- and current-clamp recordings were obtained using MultiClamp 700B amplifiers (Molecular Devices, San Jose, CA, USA), digitized via Digidata 1550B interface, and acquired at 20 kHz using pClamp 10 software (Molecular Devices). The same software was used for data analysis. Cells were selected based on neuronal morphology (round soma with neurite-like processes), while astrocyte-like cells were excluded. Inclusion criteria for analysis were as follows: (i) visual confirmation of GFP or DsRed fluorescence at the pipette tip; (ii) attachment of the labelled soma upon suction; (iii) seal resistance between 4–18 GΩ; (iv) initial series resistance < 40 MΩ, remaining stable (<20% change) throughout the recording. Whole-cell capacitance and series resistance were not compensated. During current-clamp recordings, the membrane potential was maintained at −70 mV or adjusted as necessary using a holding current. Passive and active membrane properties were assessed by applying series of hyperpolarizing and depolarizing current steps (2- or 10-pA increments, 500 ms). The instantaneous firing frequency (IFF) was calculated as the inverse of the interspike interval, derived from the time difference between adjacent action potential peaks. Maximum firing frequency was calculated by dividing the number of action potentials by the duration of the current step. Resting membrane potential (RMP) was measured immediately after break-in (I = 0 configuration). Input resistance (Rin) was determined from the voltage response to a −20 pA, 500-ms hyperpolarizing pulse using Ohm’s law (Vhold = −70 mV). Membrane capacitance (Cm) was calculated from the time constant of the membrane (τm = Rin × Cm), where τm was obtained by fitting a single exponential to the voltage response to a −20 pA, 500-ms step. Action potential properties were analysed from the first spike at rheobase. Spike amplitude was measured from threshold to peak; afterhyperpolarization (AHP) amplitude was measured from threshold to the negative peak during repolarization; and spike width was measured at half amplitude. Spontaneous excitatory postsynaptic currents (sEPSCs) were recorded for 60 s in gap-free mode at Vhold = −70 mV and analyzed using the Clampfit template search function (Molecular Devices). No correction was applied for liquid junction potential between the pipette and the ACSF.

#### Morphological Identification of Recorded Cells

Retrovirally transduced cells expressing DsRed, Mscarlet or GFP were first imaged in acute slices. During whole-cell recordings, cells were filled with biocytin through the patch pipette. Following recordings, slices were fixed by immersion in 4% paraformaldehyde (PFA) for 24 hours, rinsed in TBS, and blocked in 0.5% BSA in TBS for 1 hour. Sections were subsequently incubated in TBS containing 0.3% Triton X-100 and a streptavidin–fluorophore conjugate (1:400) for 2 hours, then mounted for confocal imaging.

### Assignment of layer identities

Coronal brain sections were collected 12 or 28 days after unilateral or bilateral cortical injections of retroviruses encoding Ascl1SA6–Dlx2–dsRed/mScarlet or Ascl1SA6–Bcl2–dsRed/mScarlet. Sections were immunostained for RFP and DAPI following the immunohistochemistry protocol described above. To minimize bias from the accumulation of infected cells at the injection site, only sections located at least 80 µm away from the injection site were analyzed. Images of the entire cortical area were acquired at 10× magnification using a Zeiss LSM 800 confocal microscope (Carl Zeiss Microscopy GmbH, Oberkochen, Germany). The boundary between cortical layers 1 and 2 was delineated on the DAPI channel using ZEN Blue software (Carl Zeiss Microscopy GmbH), with reference lines from the Allen Brain Atlas (atlas.brain-map.org) used to define cortical layer positions in coronal slices prior to switching to the RFP channel. RFP⁺ cells within each cortical layer were quantified and expressed as a percentage of the total labeled cells. Signal at the pial surface was excluded from analysis. A minimum of three animals were used for quantification.

### Statistical Analysis

Statistical analyses for image-based quantifications were performed using GraphPad Prism Version 10.6.1 (799) (GraphPad Software, San Diego, CA, USA). Data are presented as mean ± SD or as box-and-whisker plots depicting the median, 25th and 75th percentiles, and range. Data distributions were assessed for normality using the Shapiro–Wilk test. For comparisons between two groups, independent-sample t tests were applied to normally distributed datasets, whereas the Mann–Whitney U test was used for non-normally distributed data. The specific statistical tests employed for each experiment are indicated in the figure legends. The number of independent experiments (n), cells analyzed, and recorded cells used for electrophysiological analyses are reported in the respective figure legends.

For other datasets, including high-throughput sequencing and electrophysiology analyses, detailed information on the statistical methods, sample sizes, thresholds, and multiple testing corrections is provided in the Supplementary Tables, ensuring full transparency and reproducibility.

### Data availability

All single-cell and single-nucleus RNA-seq datasets generated in this study have been deposited in the Gene Expression Omnibus (GEO) and will be publicly available upon publication. Additional information, materials, or code required to reproduce or reanalyze the data presented in this study can be obtained from the corresponding authors upon reasonable request.

### Code availability

All scripts and pipelines used for bioinformatic analyses in this study have been deposited in GitHub. The repository URL will be made publicly accessible upon publication. Additional details or guidance for reproducing analyses can be obtained from the corresponding authors upon reasonable request.

## AUTHOR CONTRIBUTIONS

Conceptualization: AC, BB, FG.

Methodology and experimental design: AC, AGM, ABA, NM, BB, NM, SL.

Investigation: AC, AGM, ABA, NM, YS.

Bioinformatic analysis: AC, KLG, HF, GHO.

Resources: MS.

Writing: original draft: AC, BB.

Writing: review & editing: FG, SL, GHO, AGM, NM, KLG

Supervision: BB, FG.

Funding acquisition: AC, BB, FG.

All authors reviewed and approved the manuscript.

## ACKNOWLEDGMENTS

We thank the members of the BB and FG laboratories for helpful discussions and critical feedback throughout the course of this study. We are grateful to N. Carvajal Garcia for excellent technical support and genotyping. We thank the support staff from the Centre for Developmental Neurobiology and the Biomedical Service Unit at KCL. We acknowledge P. Rigo and S. Ahmed-de-Prado for their support in experimental design and data analysis. We acknowledge the Vectorcore, Advanced Sequencing, Flow Cytometry, Bioinformatics & Biostatistics and Research Illustration teams at the Francis Crick Institute for technical support. We thank Carol Schuurmans (Sunnybrook Research Institute, Toronto, Canada) for the Ascl1SA6 sequence.

## EXTENDED DATA

**Extended Data Figure 1.**
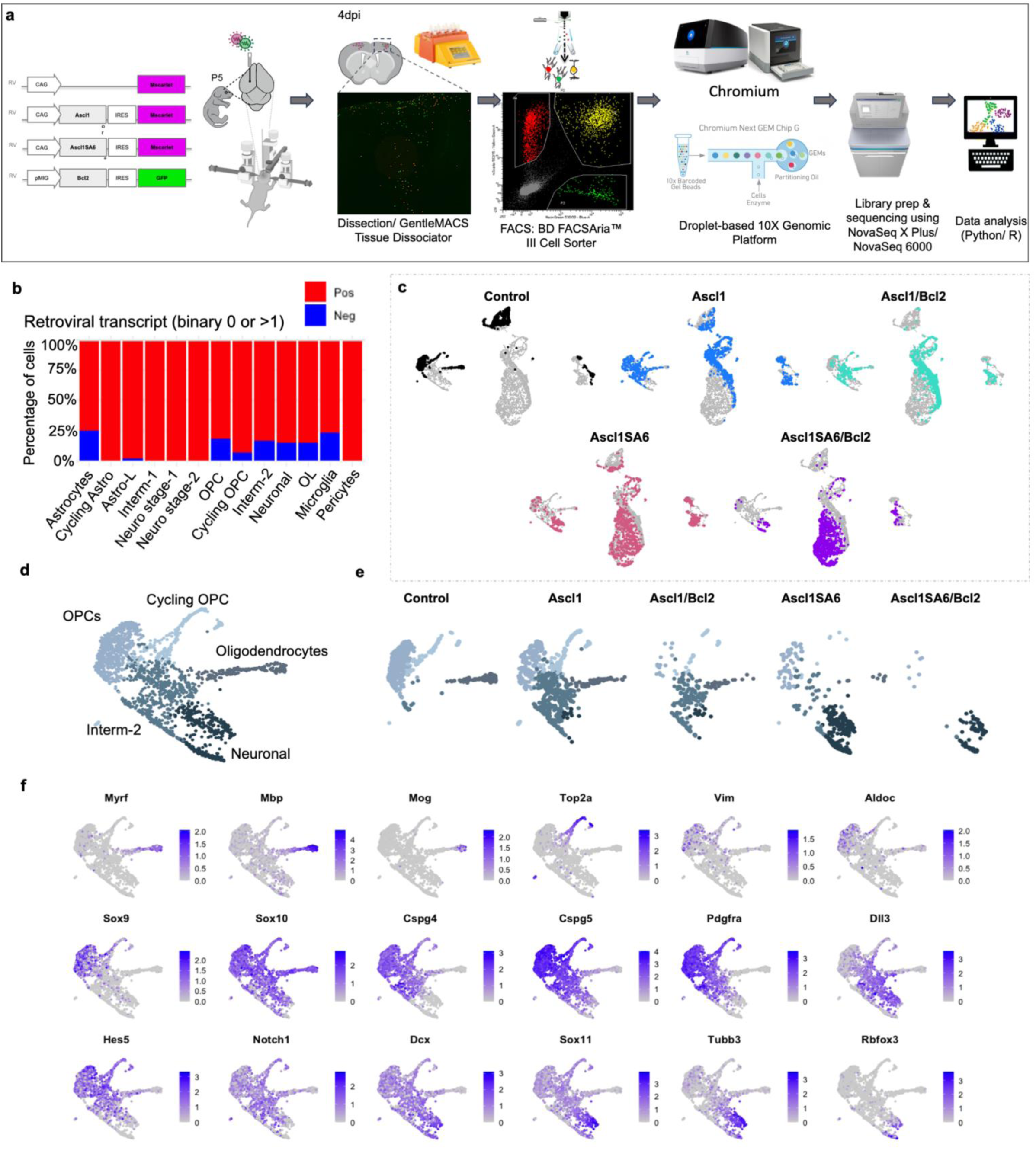
Single-cell transcriptomic analysis of retrovirally transduced glia at 4 dpi. **a,** Schematic of experimental workflow. Retroviruses encoding mScarlet-Control, Ascl1 or Ascl1SA6, with and without Bcl2 were co-injected into the somatosensory cortex at P5. Cortices were dissected and dissociated using GentleMACS Tissue Dissociator. Reporter-positive cells were isolated by FACS. Cells were prepared using 10x platform for scRNA-seq on NovaSeq X Plus or Novaseq 6000. Bioinformatical analysis performed using R or Python. **b,** Stacked bar chart summarizing the percentage of mScarlet-positive vs mScarlet-negative cells (threshold >1 transcript) across all samples at 4dpi (Control, Ascl1WT, Ascl1WT/Bcl2, Ascl1SA6, Ascl1SA6/Bcl2) combined, binary mScarlet mRNA detection (red = positive, blue = negative). **c,** UMAP visualization of retrovirally transduced cells split by condition on merged UMAP embedding (grey cells belonging to other samples). **d,** UMAP colored by cell-types showing merge of OPCs-to-neuron clusters at 4dpi indicating transition from OPCs to either differentiating oligodendrocytes or induced neuron-like cells as Neuronal. **e,** UMAP split by condition highlighting OPC-derived clusters across Control, Ascl1 or Ascl1SA6 with and without Bcl2. **f,** UMAP feature plots by gene expression, highlighting lineage and neurogenic markers across OPCs-to-neuron clusters. Feature plots showing expression of markers of radial glial (Vim, Aldoc, Sox9), OPC maturation (Sox10, Cspg4), neural progenitors (*Hes5, Notch1*) and neurons (*Sox11, Tubb3, Rbfox3*) in OPC clusters.

**Extended Data Figure 2.**
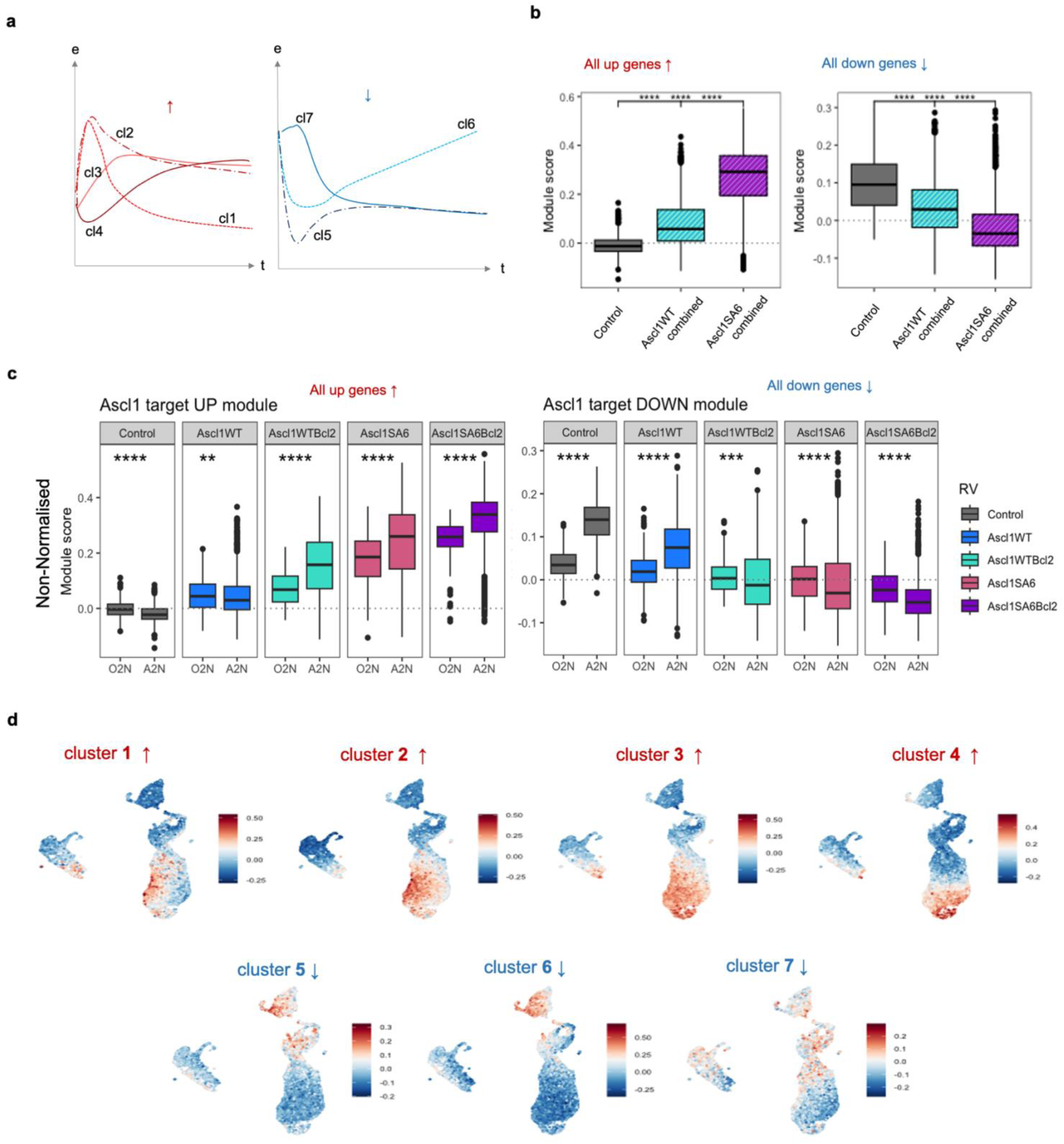
Temporal dynamics and module scores of Ascl1 target gene clusters across glial lineages. **a,** Average temporal expression profiles of gene clusters associated with activation (clusters 1–4, red, top) and repression (clusters 5–7, blue, bottom) of Ascl1 targets, Raposo et al., 2015. **b,** Boxplots show Ascl1 module scores for combined upregulated clusters (left) and downregulated clusters (right), pooling all Ascl1WT and Ascl1SA6 cells, with or without Bcl2, regardless of glial origin (astrocytes and OPCs). Pairwise comparisons of *Ascl1_target_up_genes1* expression between RV groups were performed using Wilcoxon rank-sum tests, with p-values adjusted for multiple comparisons using the Benjamini–Hochberg (BH) method. Significance is indicated by asterisks: **** p ≤ 0.001, *** p ≤ 0.01, ** p ≤ 0.05, * p ≤ 0.1. Adjusted p-values smaller than 2.2×10⁻¹⁶ are reported as <2.2e-16. **c,** Boxplots show Ascl1 module scores for the combined upregulated clusters (left) and downregulated clusters (right) across OPC- and astrocyte-derived lineages per condition (non-normalized). Astrocyte-derived cells consistently scored higher for upregulated targets and lower for downregulated targets compared with OPC-derived cells. Statistical significance was assessed using Wilcoxon tests, with the following p-values: upregulated targets, Control, p = 3.05×10⁻⁹; Ascl1WT, p = 6.50×10⁻⁴; Ascl1WT-Bcl2, p = 7.33×10⁻²⁶; Ascl1SA6, p = 3.15×10⁻²⁶; Ascl1SA6-Bcl2, p = 2.60×10⁻¹⁹; downregulated targets, Control, p = 2.01×10⁻¹⁵⁸; Ascl1WT, p = 1.31×10⁻⁷⁰; Ascl1WT-Bcl2, p = 1.89×10⁻⁴; Ascl1SA6, p = 1.14×10⁻⁶; Ascl1SA6-Bcl2, p = 7.03×10⁻⁵. The horizontal dotted line indicates a module score of 0. **d,** UMAP projection of merged samples (Control, Ascl1, Ascl1/Bcl2, Ascl1SA6, Ascl1SA6/Bcl2) showing module scores derived from all upregulated (clusters 1–4, red) and downregulated (clusters 5–7, blue) gene clusters.

**Extended Data Figure 3.**
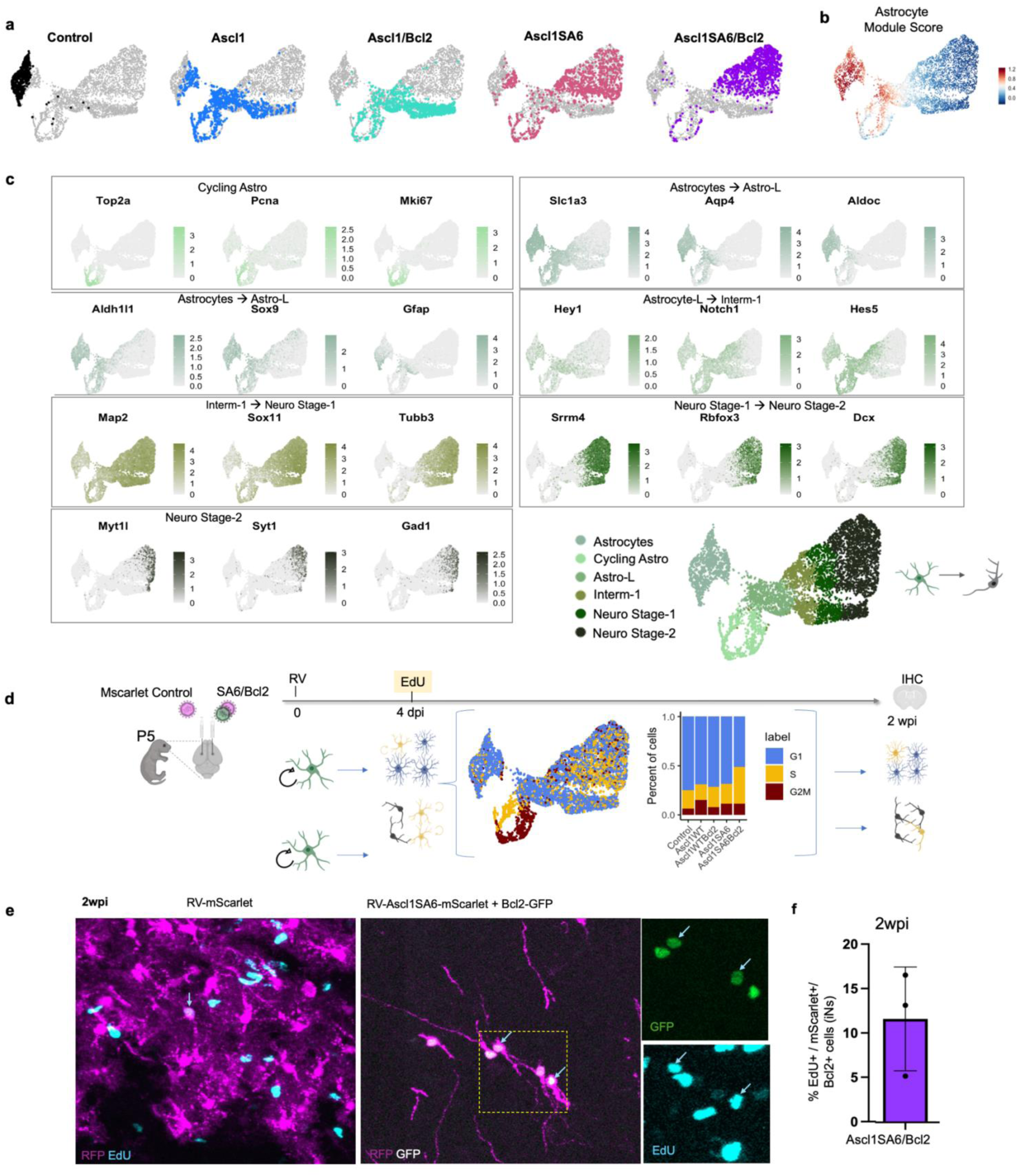
Single-cell transcriptomes and proliferation of RV-transduced glia at 4 dpi. **a,** UMAPs colored by five RV samples, split by condition. Control: 633 cells, Ascl1/SA6 samples: 1000 cells per plot. **b,** Module score generated from genes expressed in cortical astrocytes in the developing mouse cortex ^34^ **c**, UMAP feature plots colored based on annotated cell type, highlighting lineage and neurogenic markers across all cells at 4dpi, bottom right corner shows annotated cells **d**, Experimental timeline: retroviral (RV) injection at day 0, EdU administration at 4 days post-injection (dpi), and immunohistochemical (IHC) analysis at 2 weeks post-injection (wpi). UMAP embedding of single-cell transcriptomes colored by cell cycle phase (G1, S, G2/M) and corresponding stacked bar plots showing cell cycle phase distributions per RV condition. **e,** Representative IHC images at 2wpi showing control (left) or Ascl1SA6 + Bcl2 (right) transduced cells. Magenta: mScarlet; light blue: EdU. Higher-magnification insets (yellow boxes) highlight co-transduced cells expressing Bcl2 (green) and EdU (light blue). **f,** Quantification of EdU+ cells among transduced populations. Ascl1SA6-mScarlet + Bcl2-GFP (11.67 ± 5.85%). Data represent mean ± s.d.

**Extended Data Figure 4.**
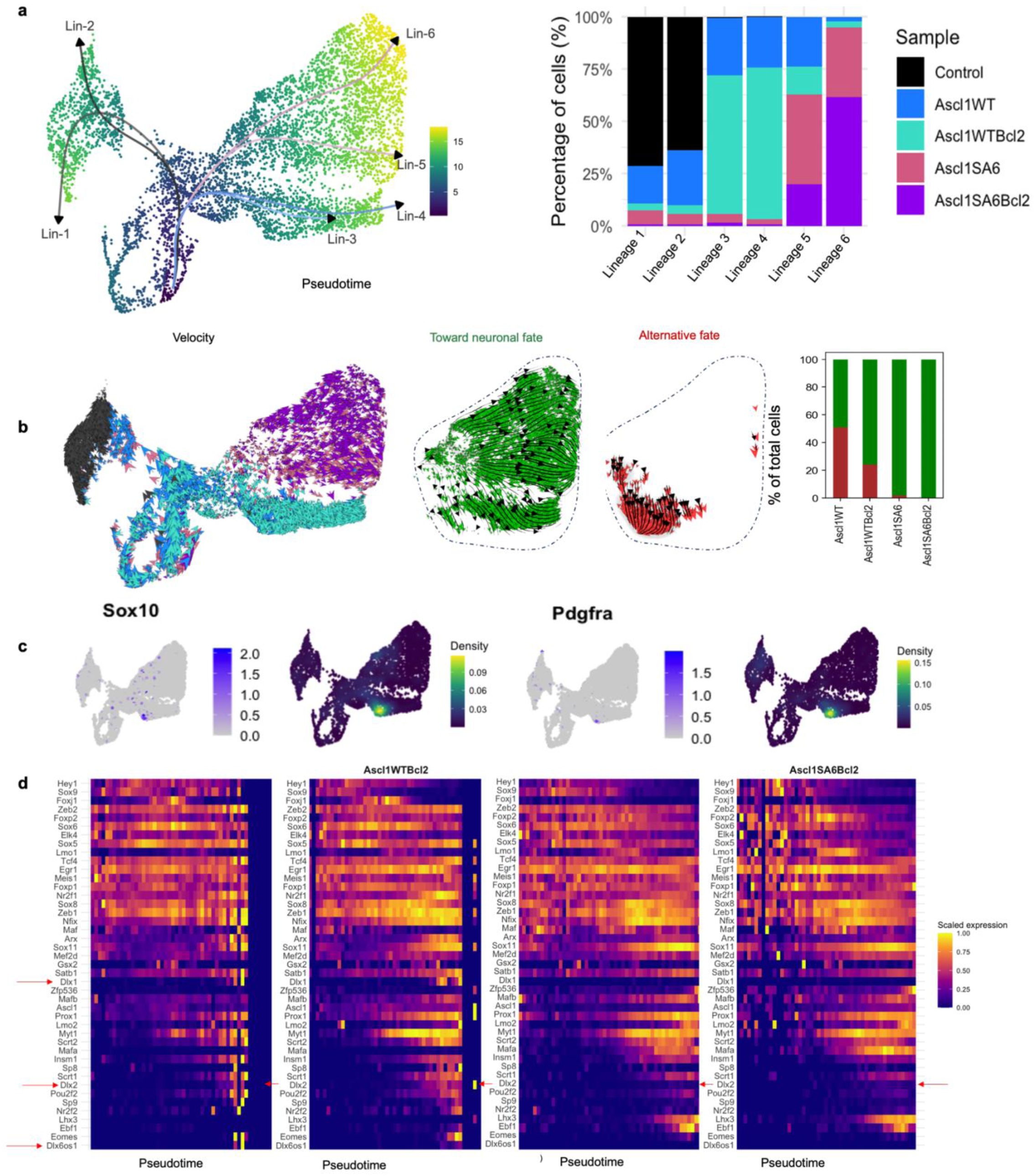
Pseudotime trajectories and RNA velocity reveal astrocyte-to-neuron reprogramming dynamics. **a,** Pseudotime and individual lineages (Lin1-6) inferred using the Slingshot algorithm (Street et al., 2018) from the 4dpi merged dataset. Lineages 1–2 represent transitions from cycling cells to differentiated astrocytes; lineages 3–6 capture astrocyte-to-neuron reprogramming (Lin3 and Lin4 from wtAscl1; Lin5 and Lin6 from Ascl1SA6) (left). Stacked bar plot showing the percentage of cells from each RV sample assigned to each pseudotime lineage (right). **b,** RNA velocity–derived UMAP showing individual cell arrows colored by RV sample (left), RNA velocity stream plots and arrows of individual cells colored by category: toward neuronal fate and towards alternative fate (both only including Intermediate-1, Neuronal stages 1 and 2 within dotted line) (middle). Corresponding proportions in a stacked bar plot (Control sample excluded) (right). **c**, Feature Plot (**left**) and density plot (**right**) showing the expression of oligodendrocyte precursor markers *Sox10* and *Pdgfra* in the “Towards alternative fate” UMAP location. **d,** Heatmap of selected transcription factor expression across the Pseudotime reprogramming axis.

**Extended Data Figure 5.**
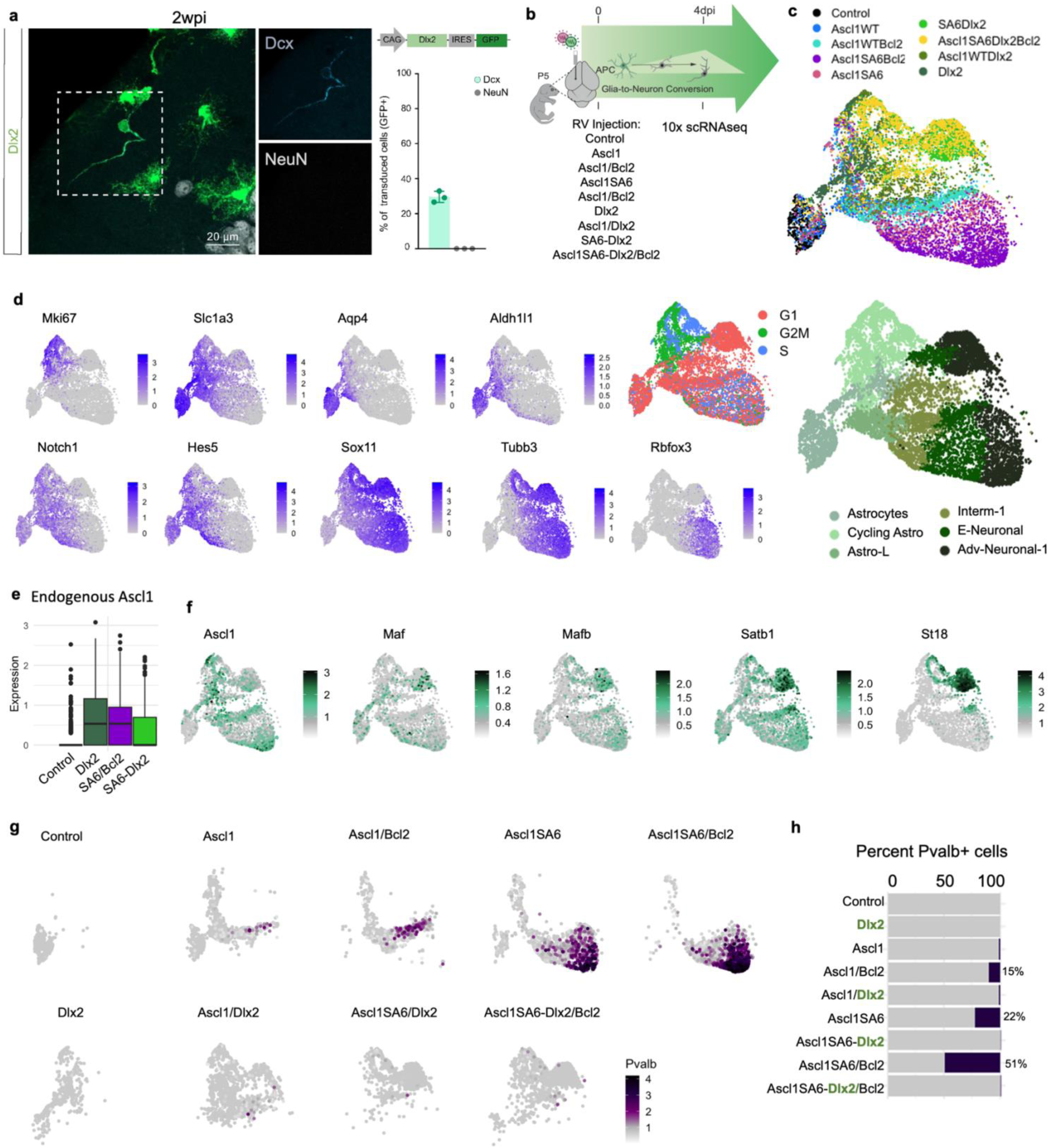
Dlx2-transduced cells activate a more canonical neuronal gene expression program while suppressing Pvalb expression. **a,** Confocal images showing expression of the immature neuronal marker Dcx (cyan) and absence of the mature neuronal marker NeuN (white) in Dlx2-transduced cells at 2wpi. Above is a schematic representation of the retroviral construct encoding Dlx2 (pCAG-Dlx2-GFP) shown. Quantification of the percentage of GFP+ transduced cells expressing Dcx (29.6 ± 3.2%) or NeuN (0.0 ± 0.0%) at 2wpi, presented as mean ± SD (n = 3 mice; 1119 cells). **b,** Experimental timeline: retroviral (RV) injection at day 0 (P5), followed by immunohistochemistry (IHC) and 10x Genomics single-cell RNA sequencing (scRNA-seq) at 4 days post-injection (dpi). **c**, UMAP embedding of astrocyte-derived cells colored by RV samples. **d**, UMAP feature plots, highlighting lineage and neurogenic markers across all cells; UMAP depicting cell cycle phases; UMAP colored by annotation cell type across all 4 dpi samples. **e**, Boxplot showing endogenous Ascl1 expression in selected RV samples. The central horizontal line in each box represents the median expression. The box spans the interquartile range (IQR; 25th to 75th percentile), and whiskers extend to 1.5× the IQR. **f,** UMAP feature plots of selected Dlx2 downstream target genes. **g,** Pvalb expression in selected samples. **h,** Stacked bar chart showing percentage of Pvalb+ cells per sample; Dlx2 samples highlighted in green, indicating suppression of Pvalb expression.

**Extended Data Figure 6.**
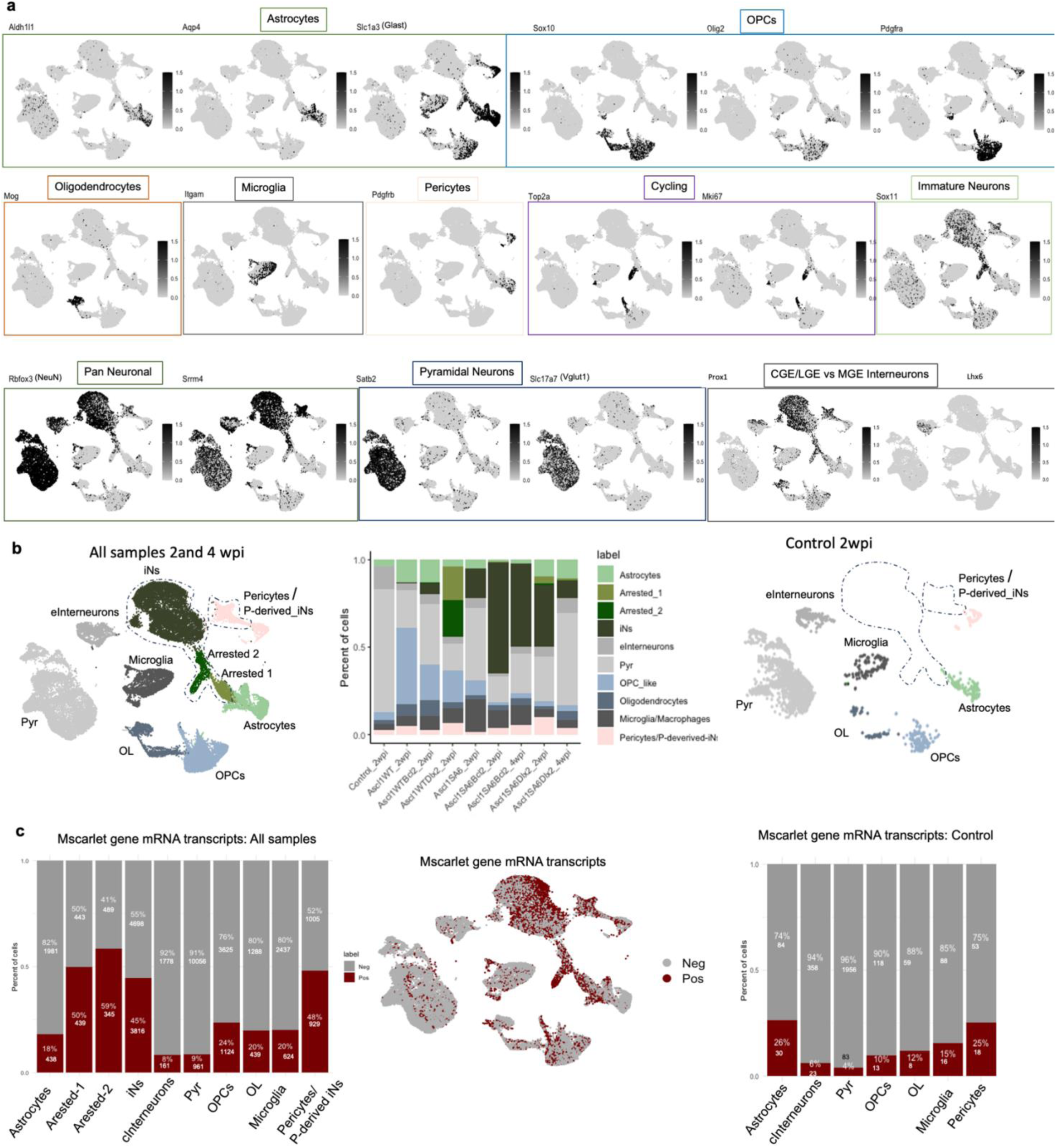
Cell-type composition and mScarlet reporter detection across 2-and 4-wpi single-cell datasets. **a,** UMAP feature plots showing marker genes corresponding to annotated cell types across all cells from 2 and 4 wpi. Sample sizes: Control_2wpi (n=2,908), Ascl1WT_2wpi (n=5,564), Ascl1WTBcl2_2wpi (n=2,806), Ascl1SA6_2wpi (n=1,896), Ascl1SA6Bcl2_2wpi (n=4,859), Ascl1WT/Dlx2_2wpi (n=3,661), Ascl1SA6/Dlx2_2wpi (n=1,970), Ascl1SA6/Bcl2_4wpi (n=6,279), Ascl1SA6/Dlx2_4wpi (n=4,205); total n=34,148 cells. **b,** UMAP colored by annotated cell identity for the merged dataset (2 and 4 wpi combined) (left) with corresponding stacked bar chart showing cell-type proportions per RV sample (middle) and UMAP colored cell type in Control sample (right). **c,** Stacked bar chart summarizing the percentage and cell counts of mScarlet-positive vs mScarlet-negative cells (threshold >1 transcript) across all samples (Control_2wpi, Ascl1WT_2wpi, Ascl1WT/Bcl2_2wpi, Ascl1SA6_2wpi, Ascl1SA6/Bcl2_2wpi, Ascl1WT/Dlx2_2wpi, Ascl1SA6/Dlx2_2wpi, Ascl1SA6/Bcl2_4wpi, Ascl1SA6/Dlx2_4wpi) combined (left); UMAP showing binary mScarlet mRNA detection (red = positive, grey = negative) using a >1 transcript threshold (middle); Stacked bar chart as in panel **c** but restricted to control samples. See technical limitations in METHODS: Technical limitations of snRNAseq.

**Extended Data Figure 7.**
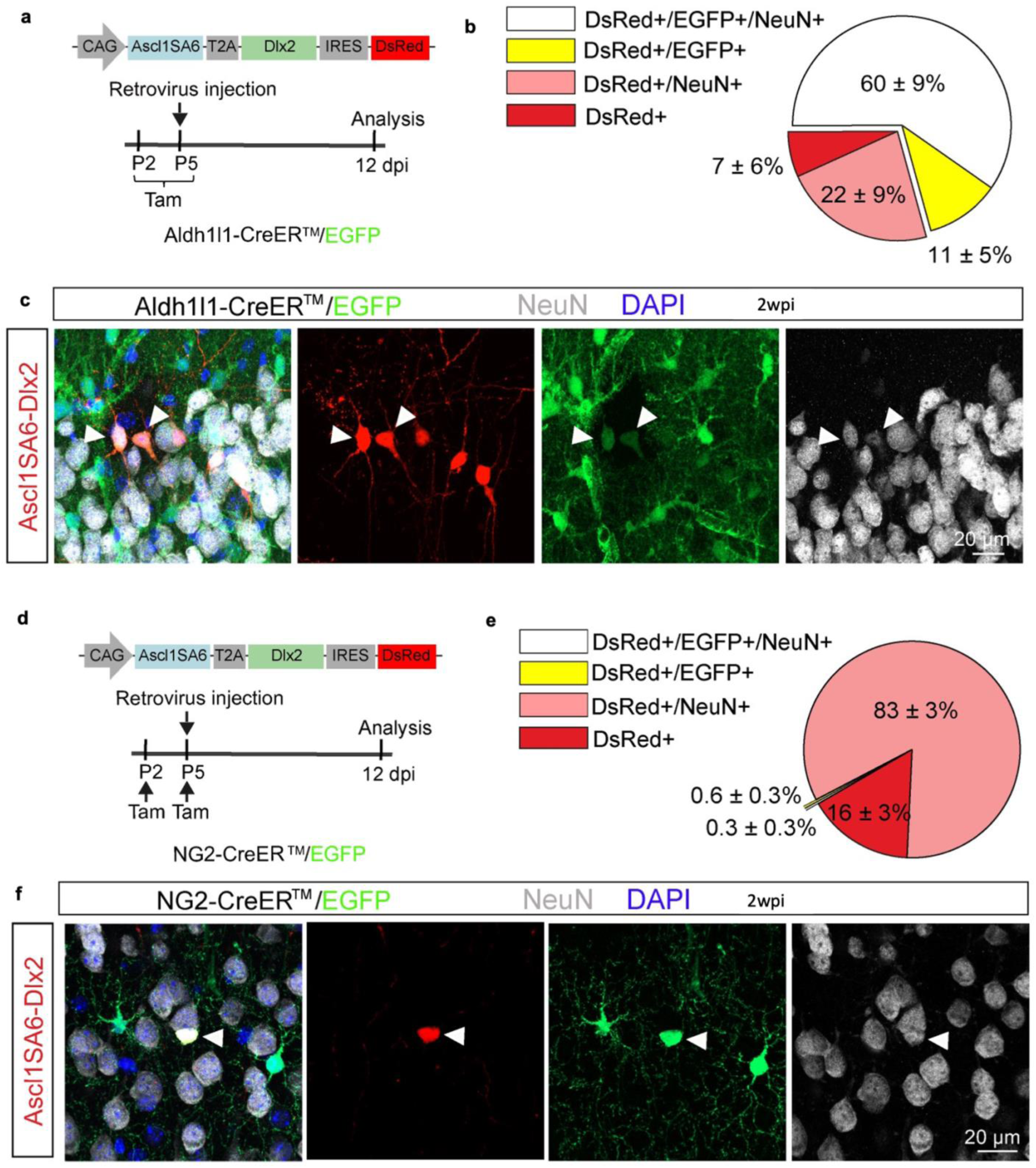
Lineage tracing reveals astrocyte-derived origins of Ascl1SA6-Dlx2-induced neurons at 2 wpi. **a,** Schematic of experimental design. Retroviral constructs encoding Ascl1SA6-Dlx2 were injected into the cortex of Aldh1l1-CreERT2;RCE:loxP mice at P5. Tamoxifen was administered daily from P2–P5 to induce Cre-mediated recombination and label astrocytes with GFP. The glial origin of transduced cells was analyzed at 2 wpi. **b,** pie chart showing the proportion of DsRed+ cells co-expressing GFP and/or NeuN in Aldh1l1-CreERT2;RCE:loxP mice (mean ± SD; n = 3 mice, 577 cells**). c,** confocal images of Ascl1SA6-Dlx2-derived iNs co-expressing NeuN together with DsRed and GFP in Aldh1l1-CreERT2;RCE:loxP mice at 2 wpi. **d,** schematic of experimental design. Retroviral constructs encoding Ascl1SA6-Dlx2 were injected into the cortex of NG2-CreERTM;RCE:loxP mice at P5. Tamoxifen was administered at P2 and P5 to label OPCs with GFP. The glial origin of transduced cells was analyzed at 2 wpi**. e,** pie chart showing the proportion of DsRed+ cells co-expressing GFP and/or NeuN in NG2-CreERTM;RCE:loxP mice (mean ± SD; n = 3 mice, 852 cells). **f,** confocal images of Ascl1SA6-Bcl2-derived iNs co-expressing NeuN and reporter genes in NG2-CreERTM;RCE:loxP mice at 2 wpi.

**Extended Data Figure 8.**
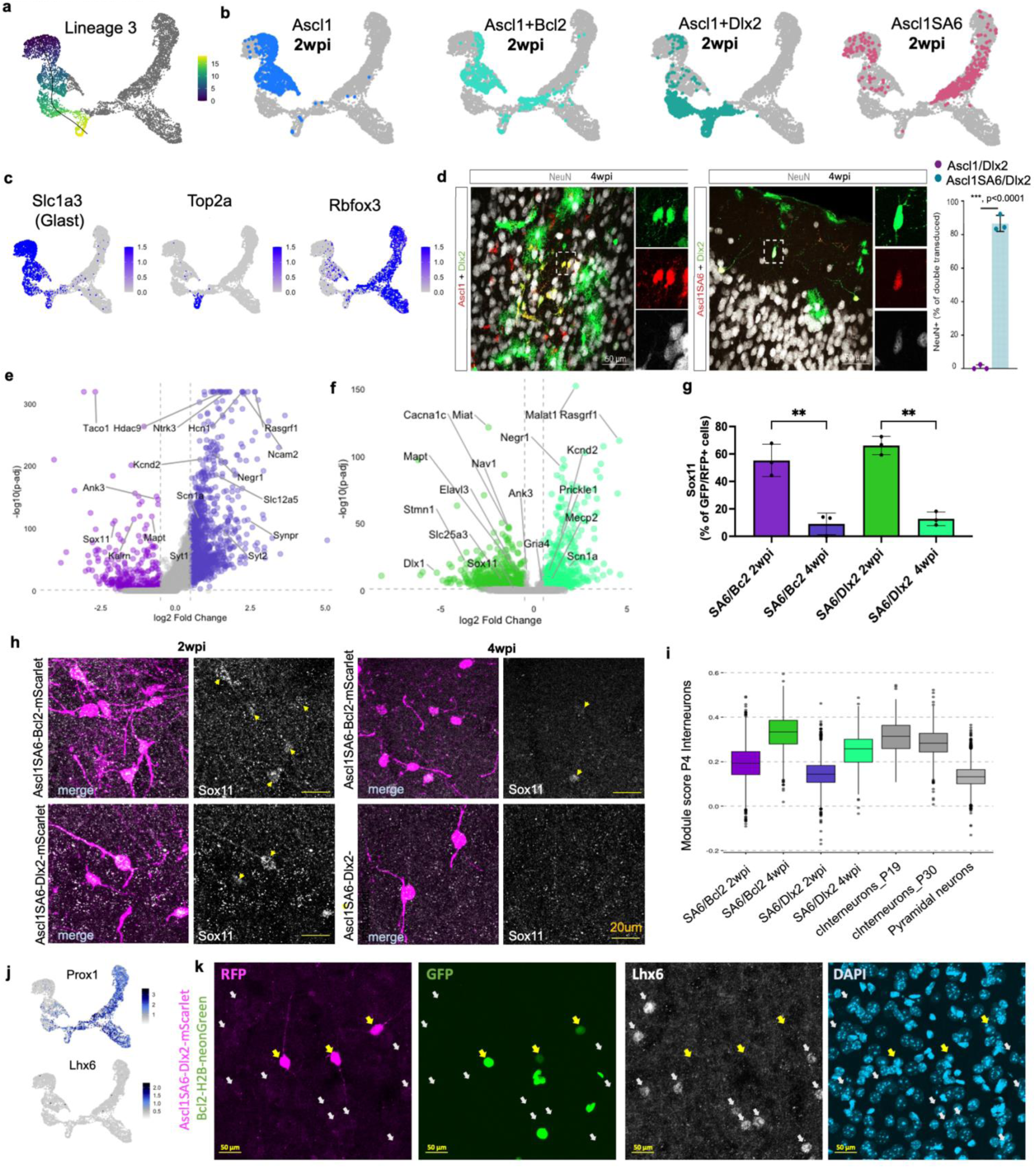
Astrocyte-derived pseudotime trajectories and maturation of Ascl1SA6/Bcl2- and Ascl1SA6/Dlx2-induced neurons. **a,** Pseudotime reconstruction (Slingshot) restricted to astrocyte-derived populations, revealing three lineages: (1) Arrested2 cycling cells, (2) Ascl1SA6/Bcl2 trajectory, and (3) Ascl1SA6/Dlx2 trajectory. Pseudotime lineage 3 in astrocyte-to-neuron clusters at 2- and 4-wpi is shown. **b,** UMAP visualization of retrovirally transduced cells split by condition on merged UMAP embedding (grey cells belonging to other samples). **c,** UMAP feature plots of selected markers in astrocyte-to-neuron clusters at 2- and 4-wpi. **d,** Left: Confocal images depicting NeuN expression (white) in Ascl1 (top) and Ascl1SA6 (bottom) + Dlx2-transduced cells at 4 wpi. Right: Quantification of NeuN+ double-transduced cells at 28 dpi (Ascl1 + Dlx2: 304 cells; Ascl1SA6 + Dlx2: 157 cells; n = 3 mice each). Data shown as mean ± SD; two-tailed unpaired Student’s t-test, ***P < 0.0001. No NeuN expression in Ascl1/Dlx2 transduced cells at 4wpi. **e,f** Differential expression between 2 and 4 wpi. Volcano plots show differentially expressed genes in Ascl1SA6-Bcl2 **(e**) and Ascl1SA6-Dlx2 (**f**) induced neurons, highlighting genes up- and downregulated over time (2 vs 4 wpi). **g,** Quantification of Sox11+ transduced cells. Sox11 expression declined significantly between 2 and 4 wpi in both conditions (SA6/Bcl2: mean diff. = 46.22, **P = 0.0040; SA6/Dlx2: mean diff. = 53.42, **P = 0.0019, Tukey’s multiple comparisons). At 2 wpi, Sox11 levels were higher in Ascl1SA6/Dlx2 compared to Ascl1SA6/Bcl2 (mean diff. = –10.86, n.s.), but by 4 wpi Sox11 expression was similarly low in both conditions (mean diff. = –3.657, n.s.). **h,** Confocal images showing Sox11 (white, initially GFP) expression in transduced cells (mScarlet, magenta). Top panels: Ascl1SA6/Bcl2 and Ascl1SA6/Dlx2 at 2 weeks post-injection (wpi). Bottom panels: Ascl1SA6/Bcl2 and Ascl1SA6/Dlx2 at 4 weeks Left: merged channels; right: Sox11 alone. **i,** Boxplots showing module scores of P4 interneurons, gene lists extracted from Arlotta et al., 2021. Data suggests Ascl1SA6/Dlx2-iNs align more closely with endogenous interneurons compared to Ascl1SA6/Bcl2-iNs. The central horizontal line in each box represents the median expression. The box spans the interquartile range (IQR; 25th to 75th percentile), and whiskers extend to 1.5× the IQR. Pyr (Pyramidal Neurons). **j,** Lhx6 and Prox1 gene expression in 2- and 4-week post injection UMAP embedding. **k,** Lhx6 staining in cortical brain slices. Ascl1SA6-Dlx2-mScarlet + Bcl2-H2B-NeonGreen, DAPI in blue, mScarlet in magenta, Bcl2 in green and Lhx6 in white. Maximum intensity projection is shown in 20x magnification. Yellow arrows: iNs, white arrows: endogenous neurons.

**Extended Data Figure 9.**
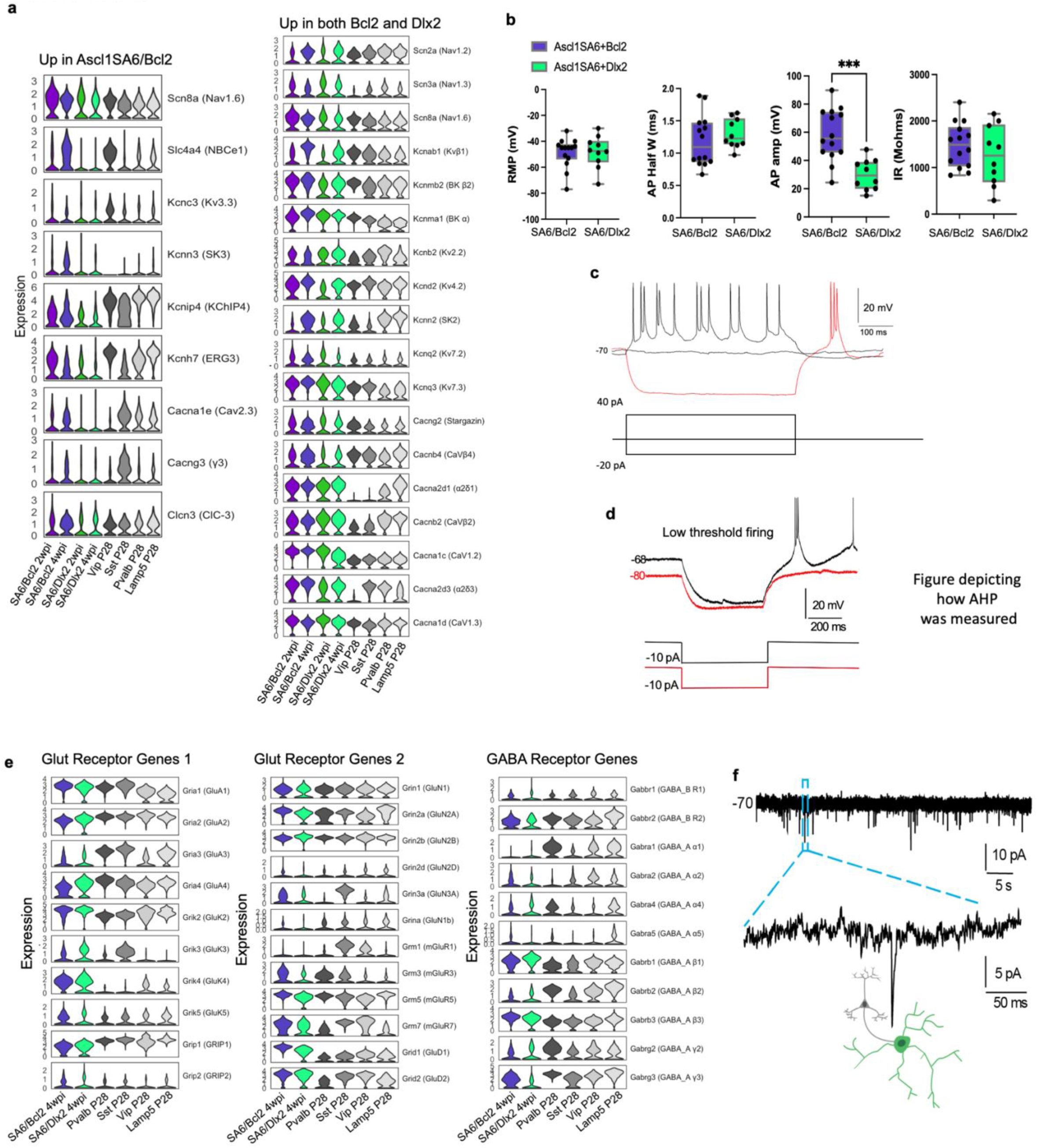
Electrophysiological properties and synaptic integration of Ascl1SA6/Bcl2- and Ascl1SA6/Dlx2-induced neurons. **a,** Stacked violin plots of ion channel genes upregulated in Ascl1SA6/Bcl2 iNs or shared between both conditions, at 2 and 4 weeks post-injection (wpi) compared with P28 endogenous cortical interneurons^41^. **b,** Quantitative analysis of membrane properties. Resting membrane potential (RMP), action potential half-width (AP half-width), AP amplitude and input resistance (IR) were measured in Ascl1SA6/Bcl2 iNs (n = 14 cells) and Ascl1SA6/Dlx2 iNs (n = 10 cells). No significant differences were detected for RMP (–48.7 ± SEM mV vs. –48.2 ± SEM; p = 0.92, Welch’s t-test) or AP half-width (1.19 ± SEM ms vs. 1.30 ± SEM; p = 0.39). In contrast, AP amplitude was significantly higher in Ascl1SA6/Bcl2 iNs (58.4 ± SEM mV vs. 30.0 ± SEM; p = 0.0001). Values for Ascl1SA6/Bcl2 from Marichal et al., 2024. IR did not differ significantly (1259 ± SEM MΩ vs. 1477 ± SEM MΩ; p = 0.38). **c,** Sustained and consistent firing (black) and rebound firing after hyperpolarization (red) in Ascl1/Dlx2-iN. **d,** Voltage-dependent rebound spiking (red) generated in Ascl1/Dlx2-iN following relief from hyperpolarization. **e,** Stacked violin plots of Glutamatergic and GABAergic receptor subunit genes in Ascl1SA6/Bcl2-iNs and Ascl1SA6/Dlx2-iNs at 4 wpi, compared with P28 endogenous cortical interneurons ^41^. **f,** Synaptic integration of Ascl1SA6/Dlx2-iNs. Spontaneous postsynaptic currents were detected in 4/5 recorded cells. Blue inset, magnified trace.

**Extended Data Figure 10.**
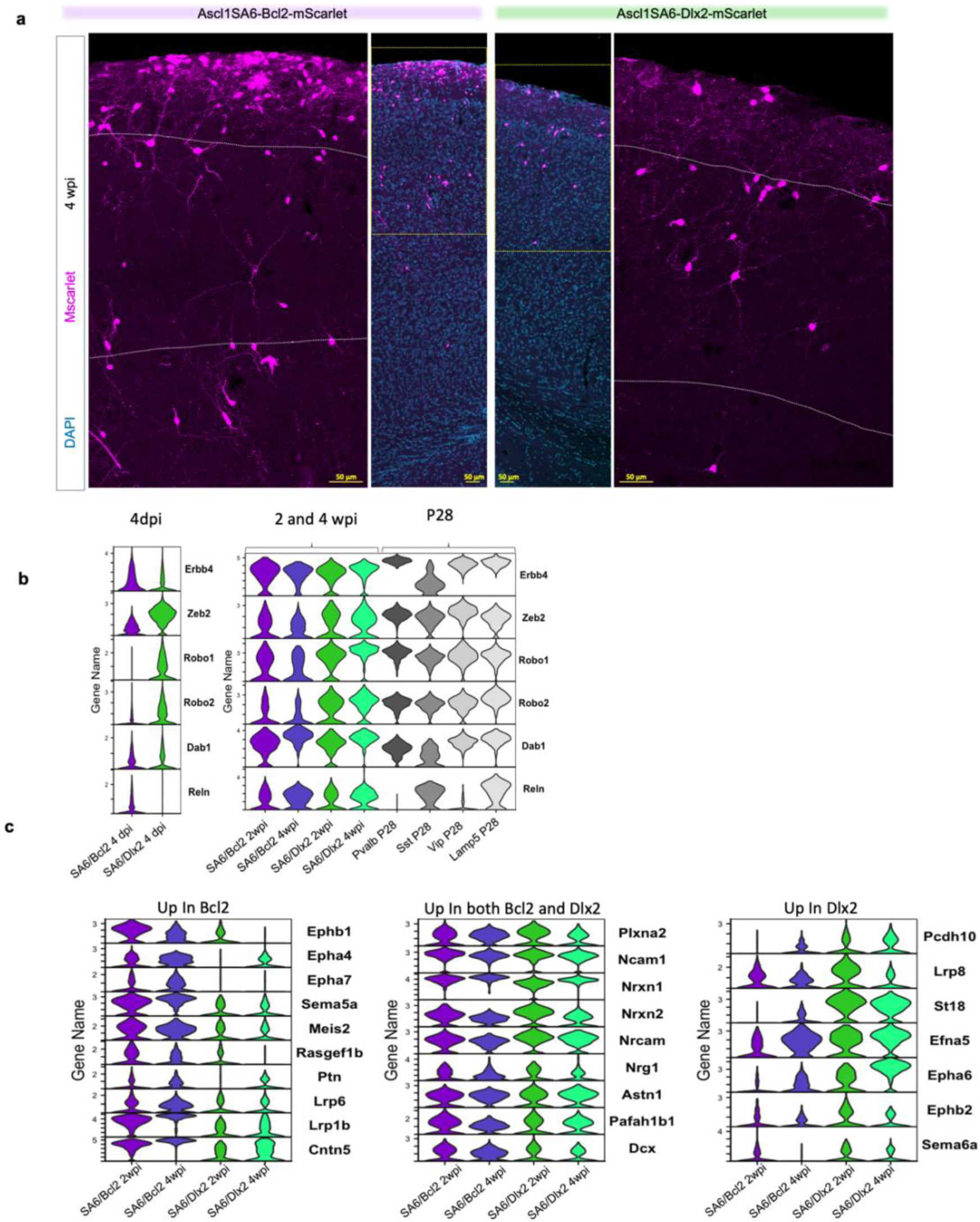
Gene expression and laminar positioning of Ascl1SA6-Bcl2-and Ascl1SA6-Dlx2-induced neurons. **a,** Ascl1SA6-Bcl2-iNs (left) and Ascl1SA6-Dlx2-iNs (right) in the cerebral cortex at 2 wpi stained with DAPI (blue). Cortical layers are delineated by dotted lines (Layer I, 2/3, 4–6**). b,** Stacked violin plots of selected genes associated with interneuronal migration at 4dpi and 2 and 4 wpi compared with P28 endogenous cortical interneurons ^41^. **c,** Differentially expressed genes associated with laminar positioning between Ascl1SA6-Bcl2 and Ascl1SA6-Dlx2 iNs at 4 wpi.

