## SUPPLEMENTARY Figures and Tables for "Dlx2 reprograms the transcriptome and laminar position of glia-derived Ascl1-induced interneurons"

### SUPPLEMENTARY DATA

**Supplementary Figure 1**

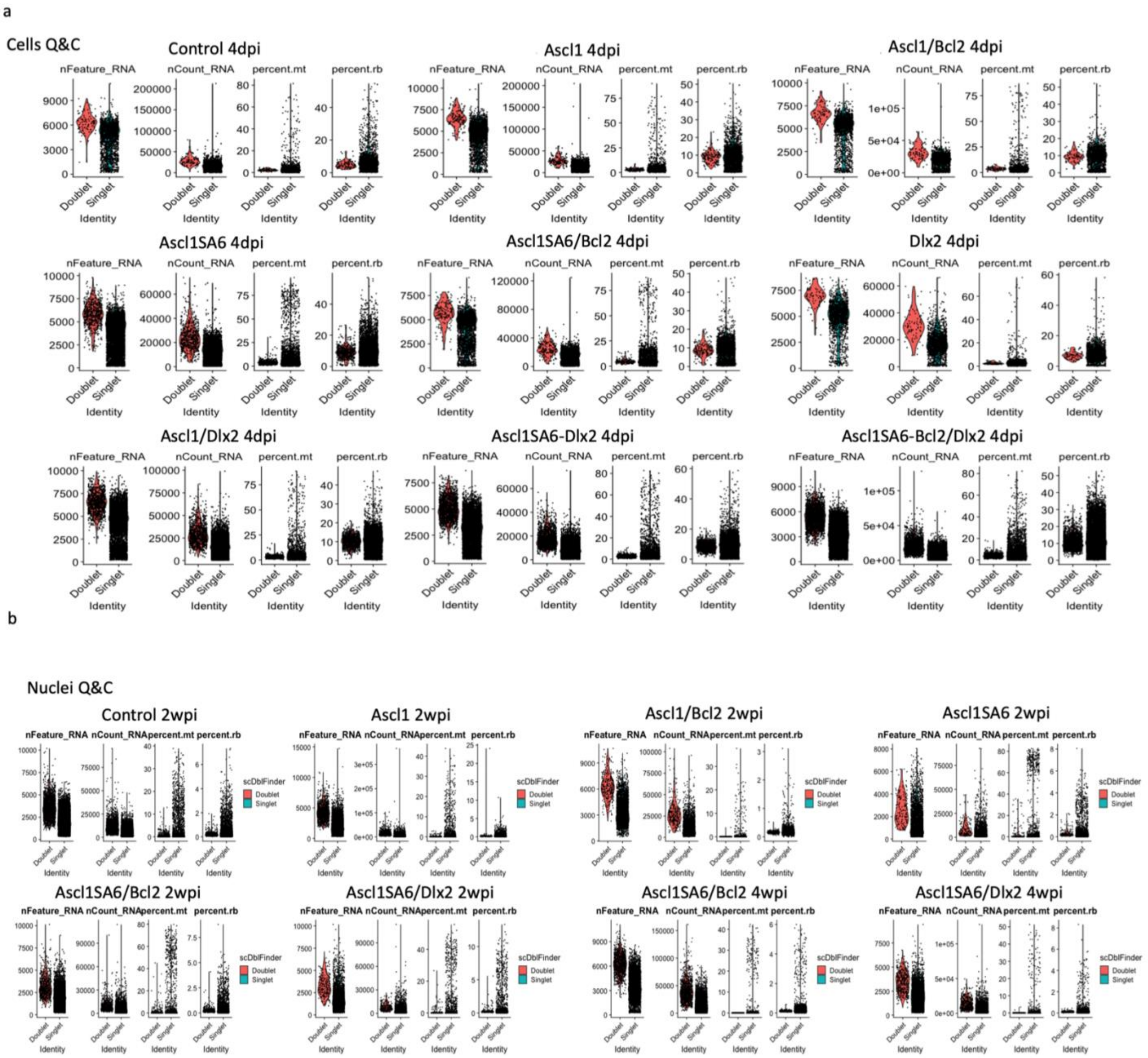

**Supplementary Figure 1. Quality control metrics for single-cell and single-nucleus RNA-seq datasets**

**a,** Cell quality control of all 4dpi samples, numbers of nFeature (genes), nCount (transcripts), percent.mt (mitochondrial) and percent.rb (ribosomal) plotted in vlnplots of doublets and singlets. Doublets were excluded. **b,** Nuclei quality control of all 2- and 4 wpi samples, numbers of nFeature (genes), nCount (transcripts), percent.mt (mitochondrial) and percent.rb (ribosomal) plotted in vlnplots of doublets and singlets. Doublets were excluded from analysis.

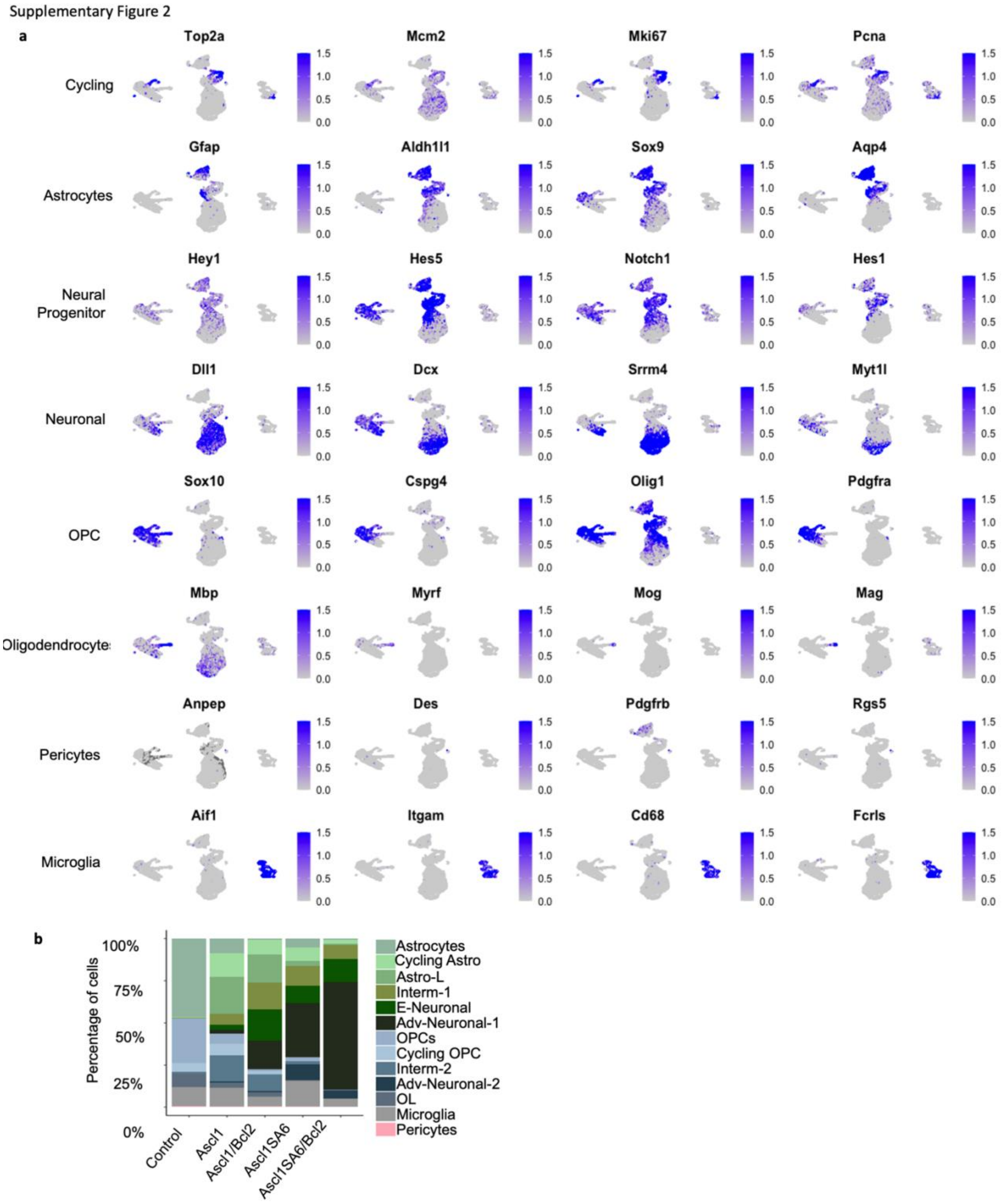

**Supplementary Figure 2. Gene expression–based lineage annotation and cell-type distribution across RV samples at 4 dpi.**

**a,** UMAP feature plots by gene expression, highlighting lineage and neurogenic markers across all cells at 4dpi. **b,** Stacked bar chart showing cell-type proportions per RV-sample. Values are found in SI Table 1.

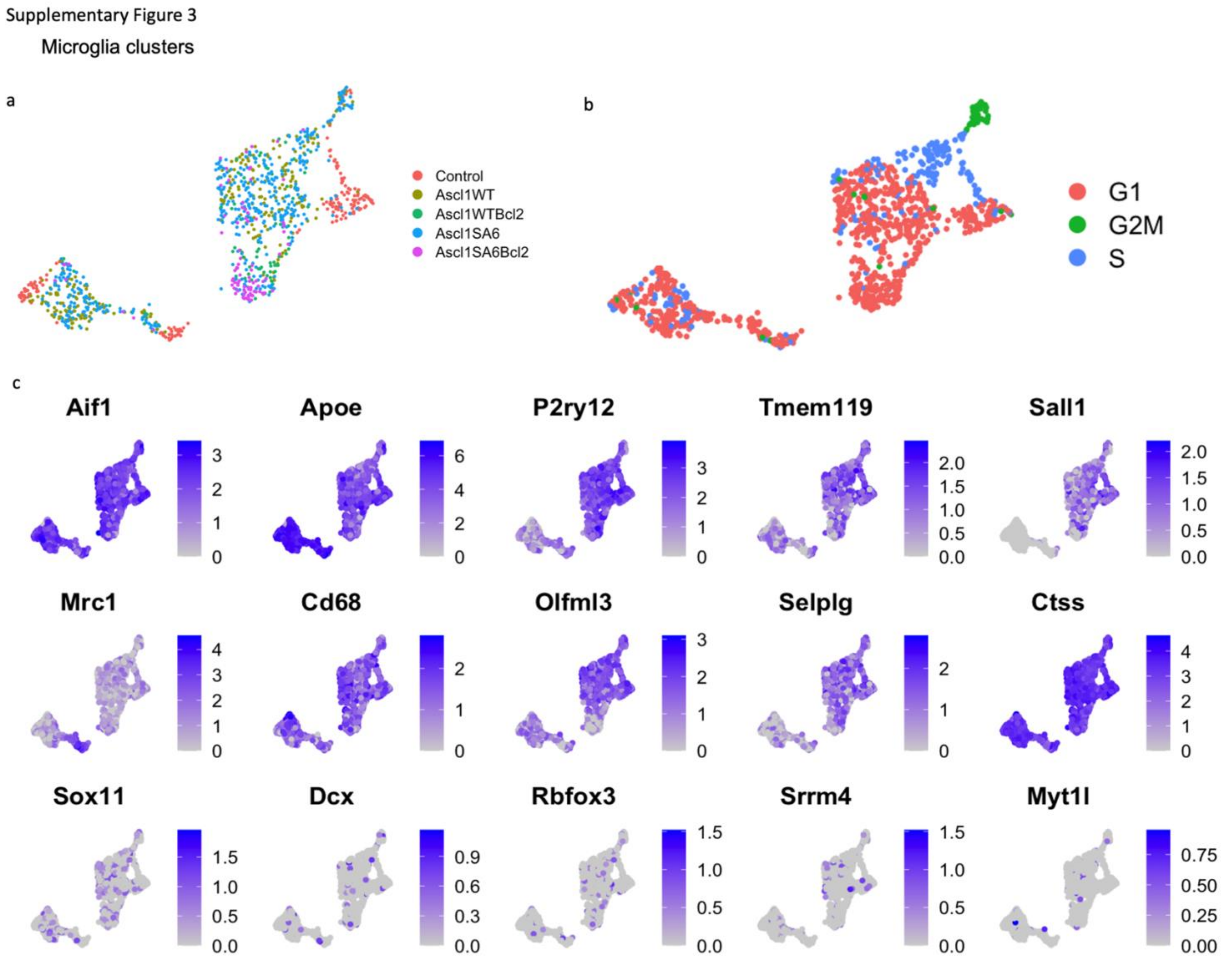

**Supplementary Figure 3. UMAP of microglial cells illustrating transcriptional profiles across Control, Ascl1, or Ascl1SA6 conditions, with or without Bcl2.**

**a,** UMAP of microglial cells showing transcriptional changes across Control, Ascl1 or Ascl1SA6 with and without Bcl2 conditions. **b,** UMAP coloured by cell cycle Phases, S, G2M and G1. **c,** Feature plots of microglial markers confirming stable expression across samples, showing minimal variation by treatment. Sub-clustering of microglia confirms transcriptional stability regardless of retroviral condition.

Supplementary Figure 4

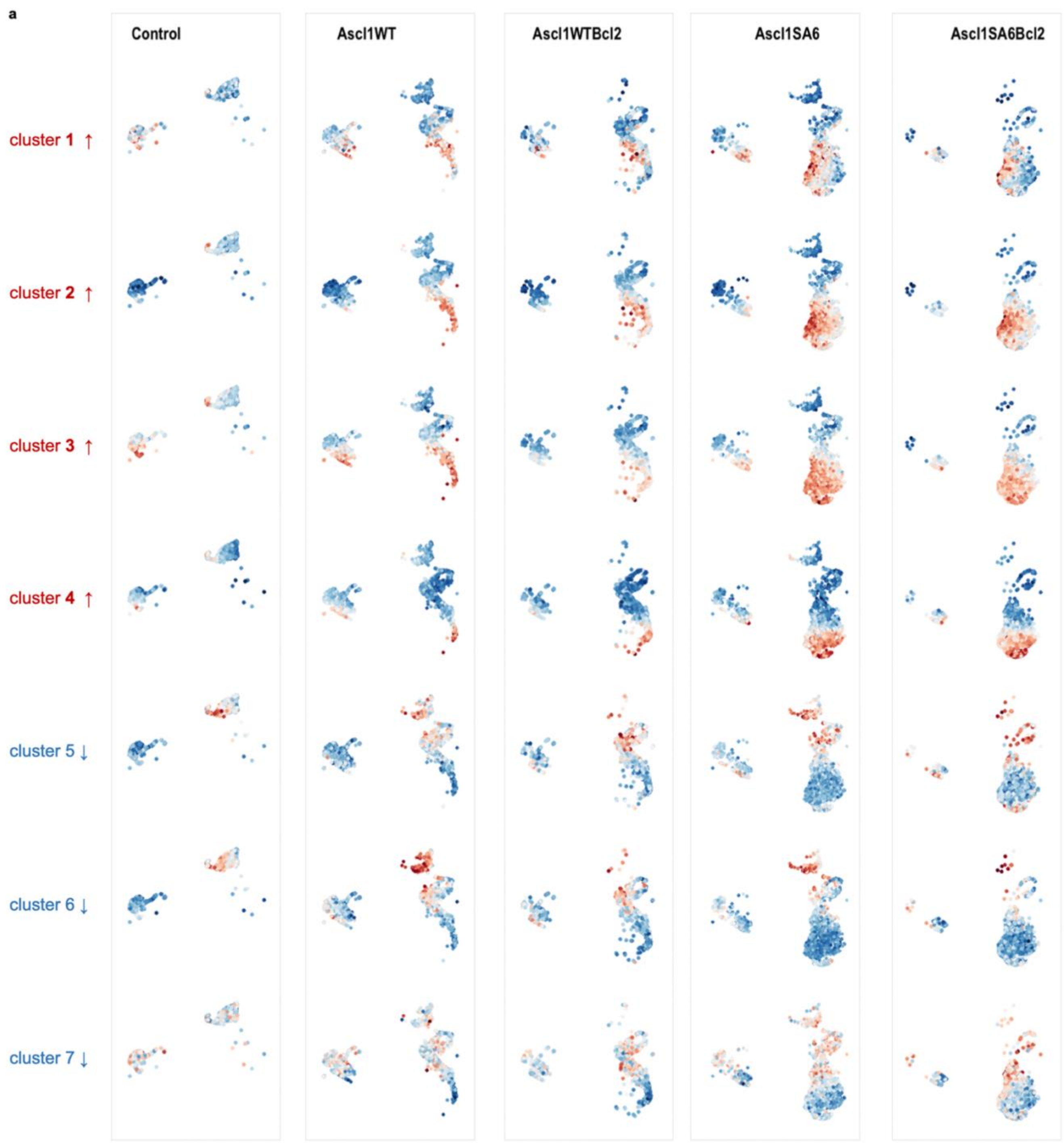

**Supplementary Figure 4. Differential gene module activity across Ascl1- and Ascl1SA6-based reprogramming conditions.**

**a,** UMAP projection split by experimental condition of five samples (Control, Ascl1, Ascl1/Bcl2, Ascl1SA6, Ascl1SA6/Bcl2) showing module scores derived from all upregulated (clusters 1–4, red) and downregulated (clusters 5–7, blue) gene clusters.

Supplementary Figure 5

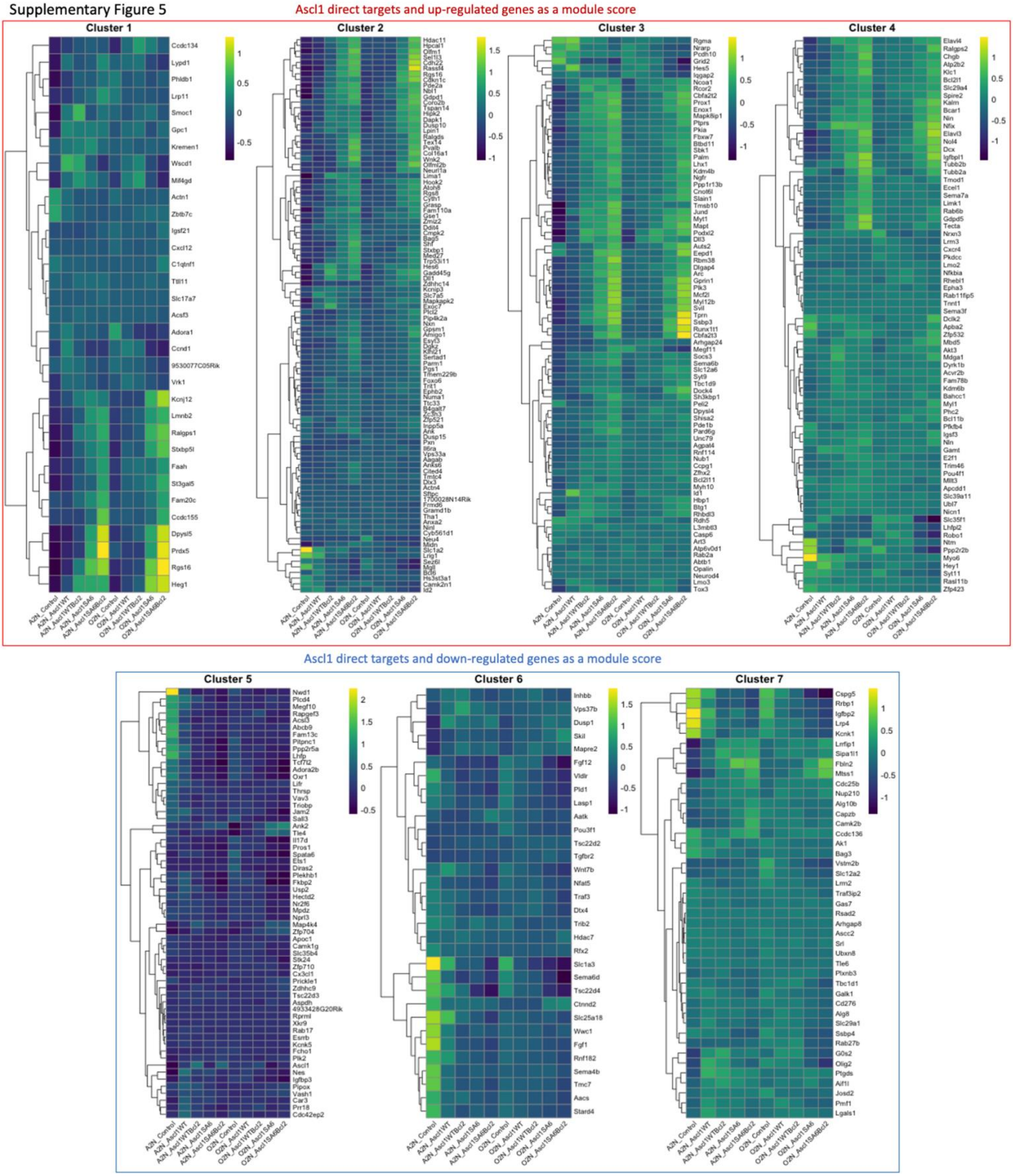

**Supplementary Figure 5. Ascl1-induced gene clusters from NSCs are differentially regulated across A2N and OPC lineages.**

**a,** Heatmap showing expression of all Ascl1-regulated gene clusters (cl1–7), previously identified in NSCs (Raposo *et al.*, 2015), across individual experimental conditions and separated by glial origin into astrocyte-to-neuron (A2N) and OPC-to-neuron (OPC) lineages. Red and blue boxes indicate upregulated and downregulated target gene clusters, respectively.

Supplementary Figure 6

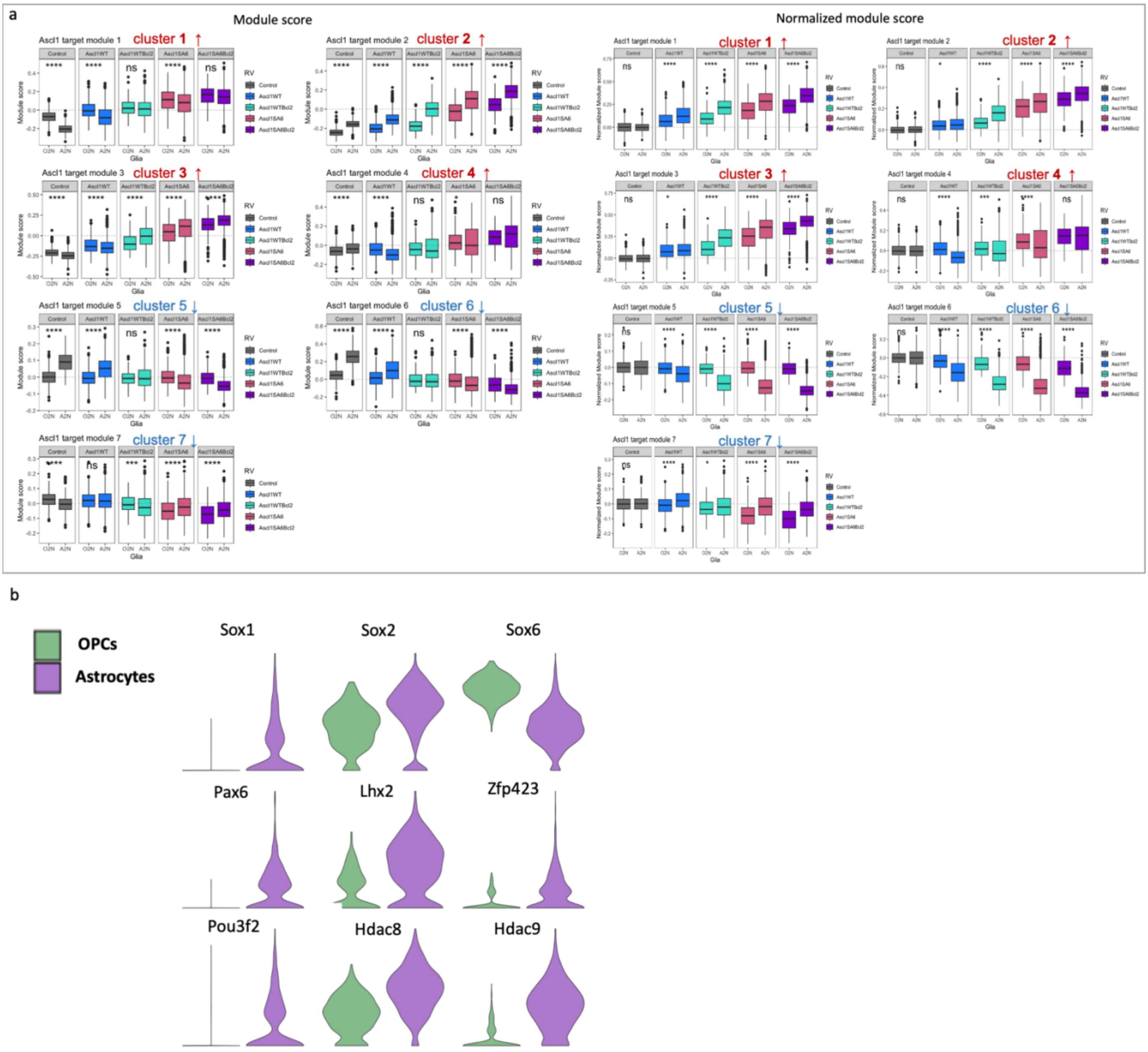

**Supplementary Figure 6. Differential Ascl1 module activation and lineage-specific transcriptional regulators in OPC- and astrocyte-derived cells.**

**a,** Boxplots showing Ascl1 module scores for combined upregulated (top) and downregulated (bottom) clusters across OPC- and astrocyte-derived lineages per condition and temporal cluster. Left panels show unnormalized scores; right panels show scores normalized to Control astrocytes and OPCs. Statistical significance was assessed by Wilcoxon test and is indicated by asterisks: **** $p \leq 0.001$, *** $p \leq 0.01$, ** $p \leq 0.05$, * $p \leq 0.1$. **b,** Violin plots of transcriptional regulators differentially expressed between OPC- and astrocyte-derived lineages.

Supplementary Figure 7

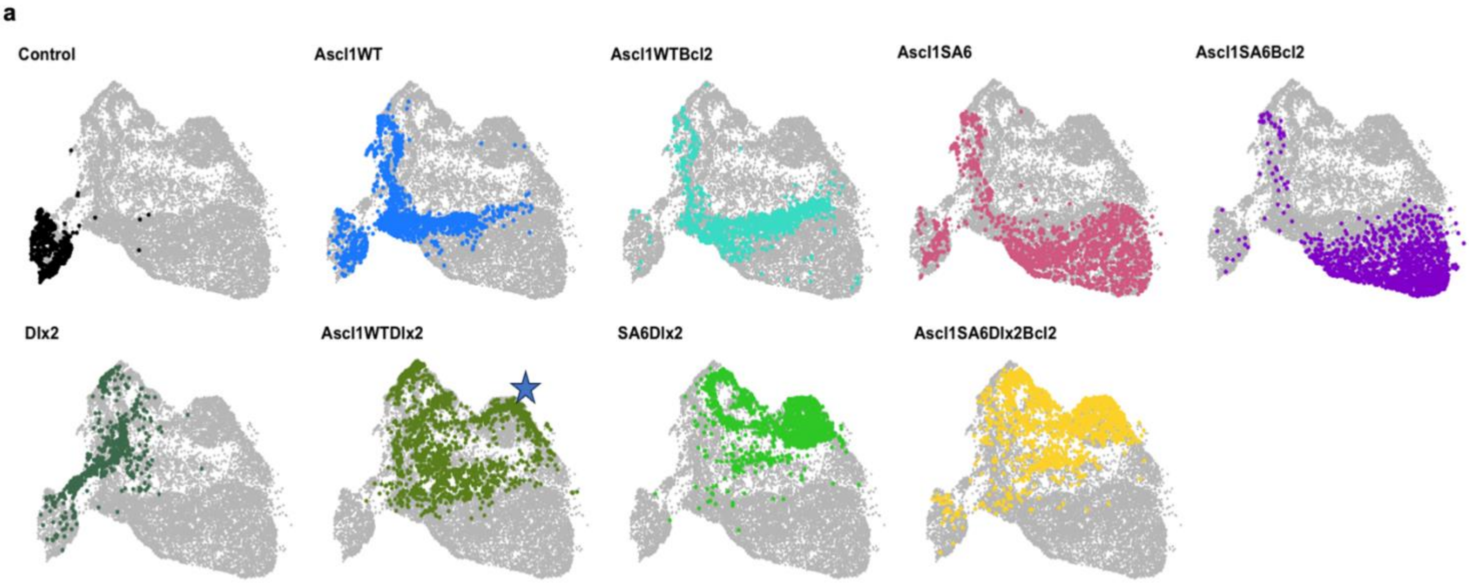

**Supplementary Figure 7. UMAP visualization of RV-infected cells at 4 dpi reveals a new neuronal cluster in Ascl1/Dlx2 and Ascl1SA6-Dlx2 samples.**

**a,** UMAP embedding of all cells sequenced at 4 dpi, split and colored by retroviral condition. The star indicates a newly appearing neuronal cluster in Ascl1/Dlx2, Ascl1SA6-Dlx2, and Ascl1SA6-Dlx2/Bcl2 samples.

Supplementary Figure 8

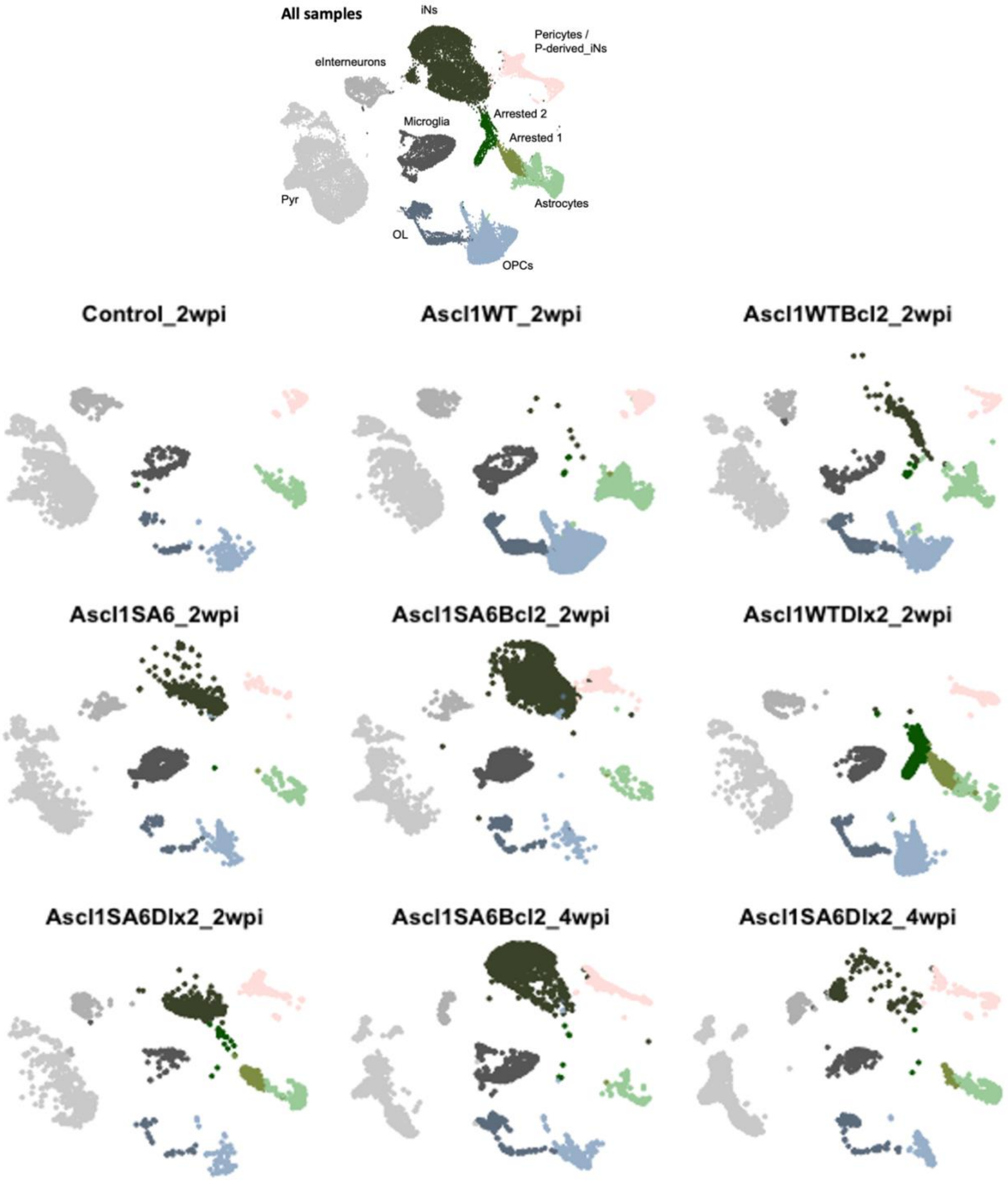

**Supplementary Figure 8. Distribution of cell types across merged 2- and 4-wpi samples, split by retroviral condition.**

**a,** UMAP embedding of all cells in the merged 2- and 4-wpi dataset, colored by cell type. Top panel shows all samples combined (Control_2wpi, Ascl1WT_2wpi, Ascl1WTBcl2_2wpi, Ascl1SA6_2wpi, Ascl1SA6Bcl2_2wpi, Ascl1WTDlx2_2wpi, Ascl1SA6Dlx2_2wpi, Ascl1SA6Bcl2_4wpi, Ascl1SA6Dlx2_4wpi); bottom panel shows each RV sample individually.

Supplementary Figure 9

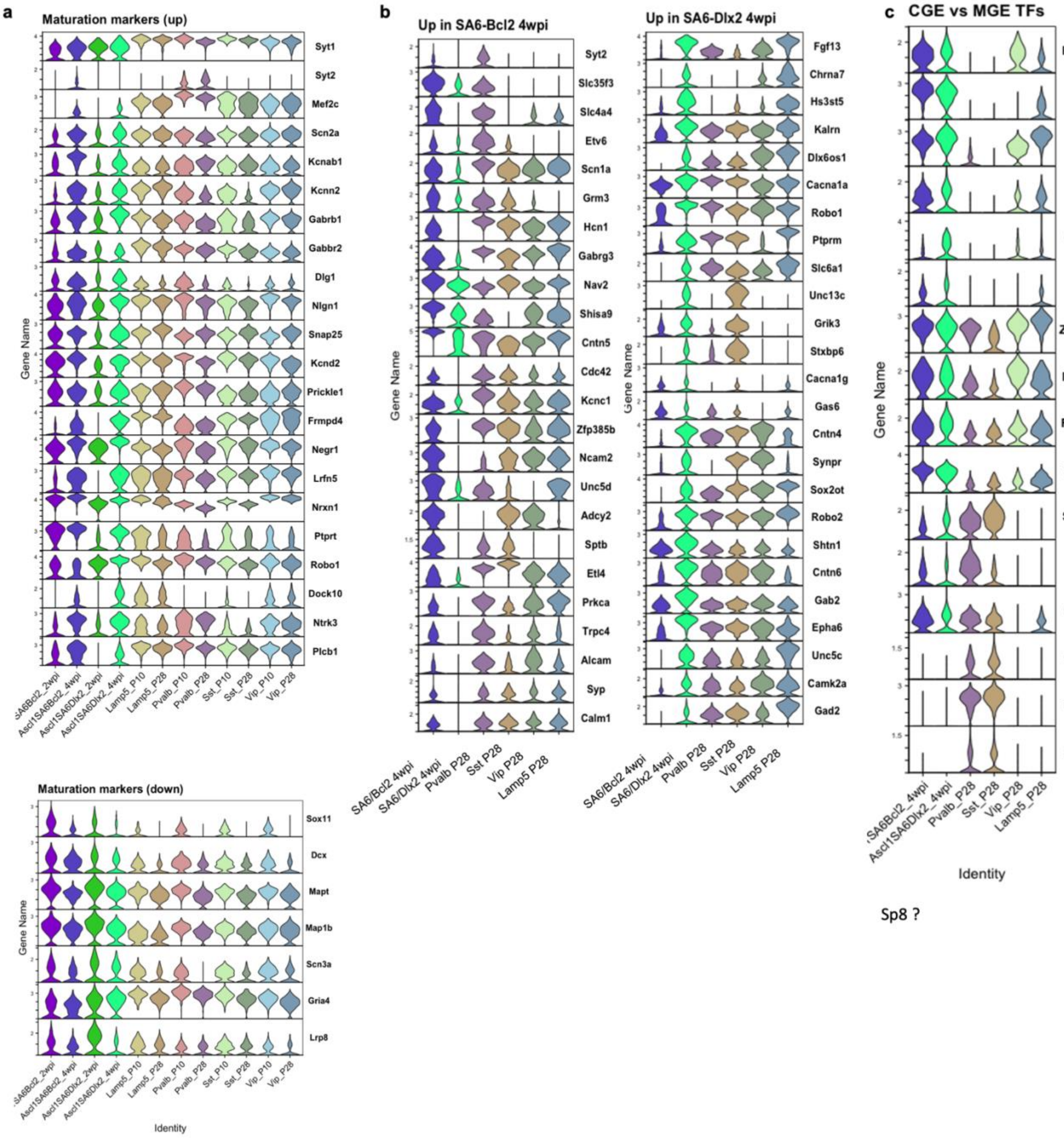

**Supplementary Figure 9. Induced neurons acquire maturation markers and diverge into interneuron subtypes compared to endogenous cortical interneurons.**

**a,** Stacked violin plots showing downregulation of immature neuronal genes and upregulation of maturation- and synapse-associated genes in induced neurons (iNs) compared to endogenous cortical interneurons at P10 and P28 (Allaway *et al.*, 2021). **b,** Stacked violin plots of interneuron subtype markers, demonstrating divergence of iN identities relative to P28 endogenous cortical interneurons. **c,** CGE- and MGE-specific transcription factors differentially expressed in iNs compared to P28 endogenous cortical interneurons (Allaway *et al.*, 2021).

Supplementary Figure 10

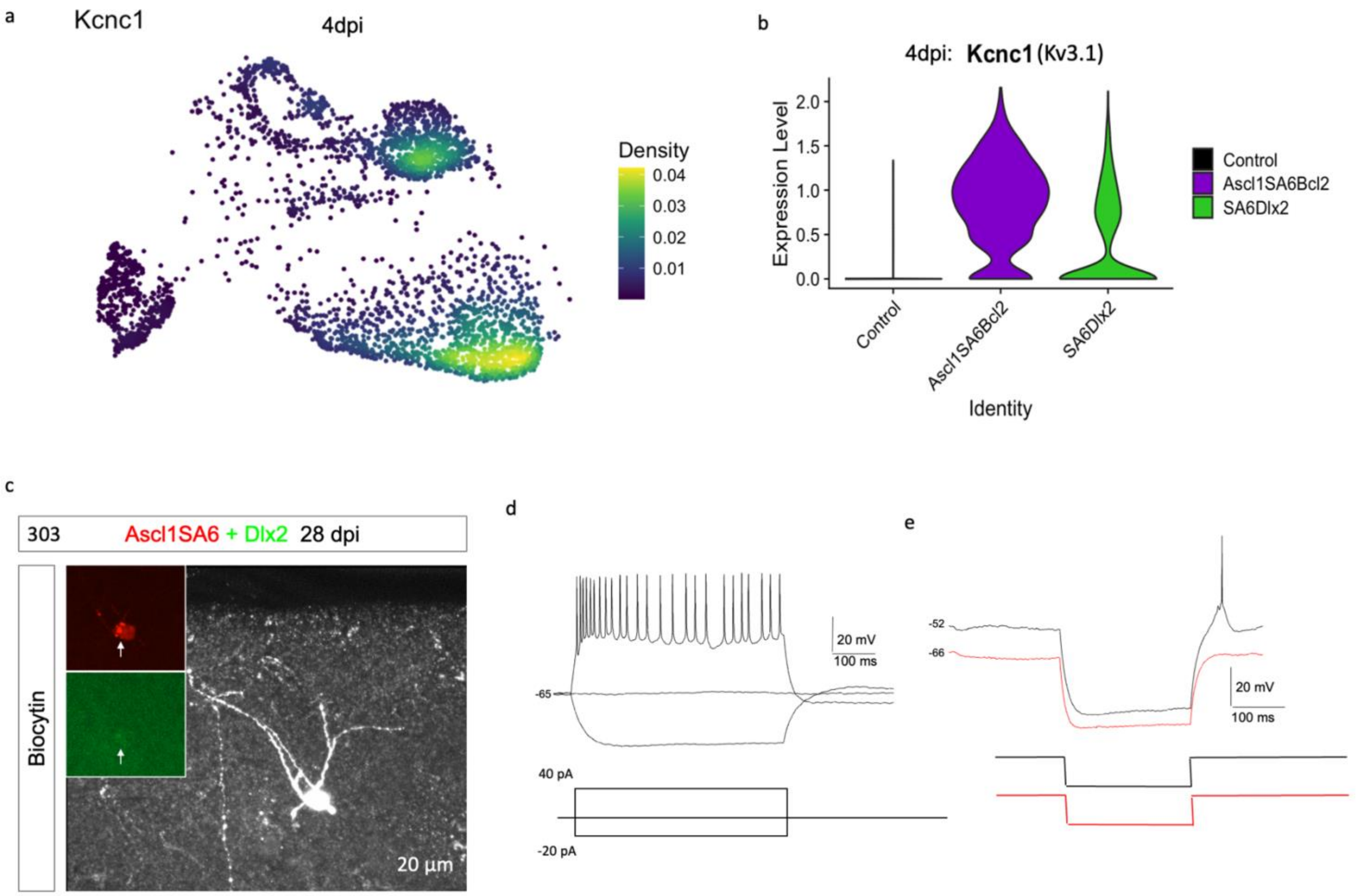

**Supplementary Figure 10. Molecular and electrophysiological characterization of Ascl1SA6/Dlx2-induced neurons.**

**a,b,** Density plot (**a**) and violin plot (**b**) of channel gene expression Kcnc1 (corresponding Protein Kv3.1) at 4dpi in merged object (Control, SA6/Bcl2 and SA6/Dlx2). **c,** Further Example of Biocytin-filled Ascl1SA6/Dlx2-iN in acute slice. Insets: GFP (Dlx2) and mScarlet (Ascl1SA6) colocalization in acute slice (left). Scale bar, 20 µm. **d,e,** Representative patch-clamp recording from Ascl1SA6/Dlx2-iN showing repetitive action potentials **(d)** and rebound spiking after hyperpolarization **(e).**

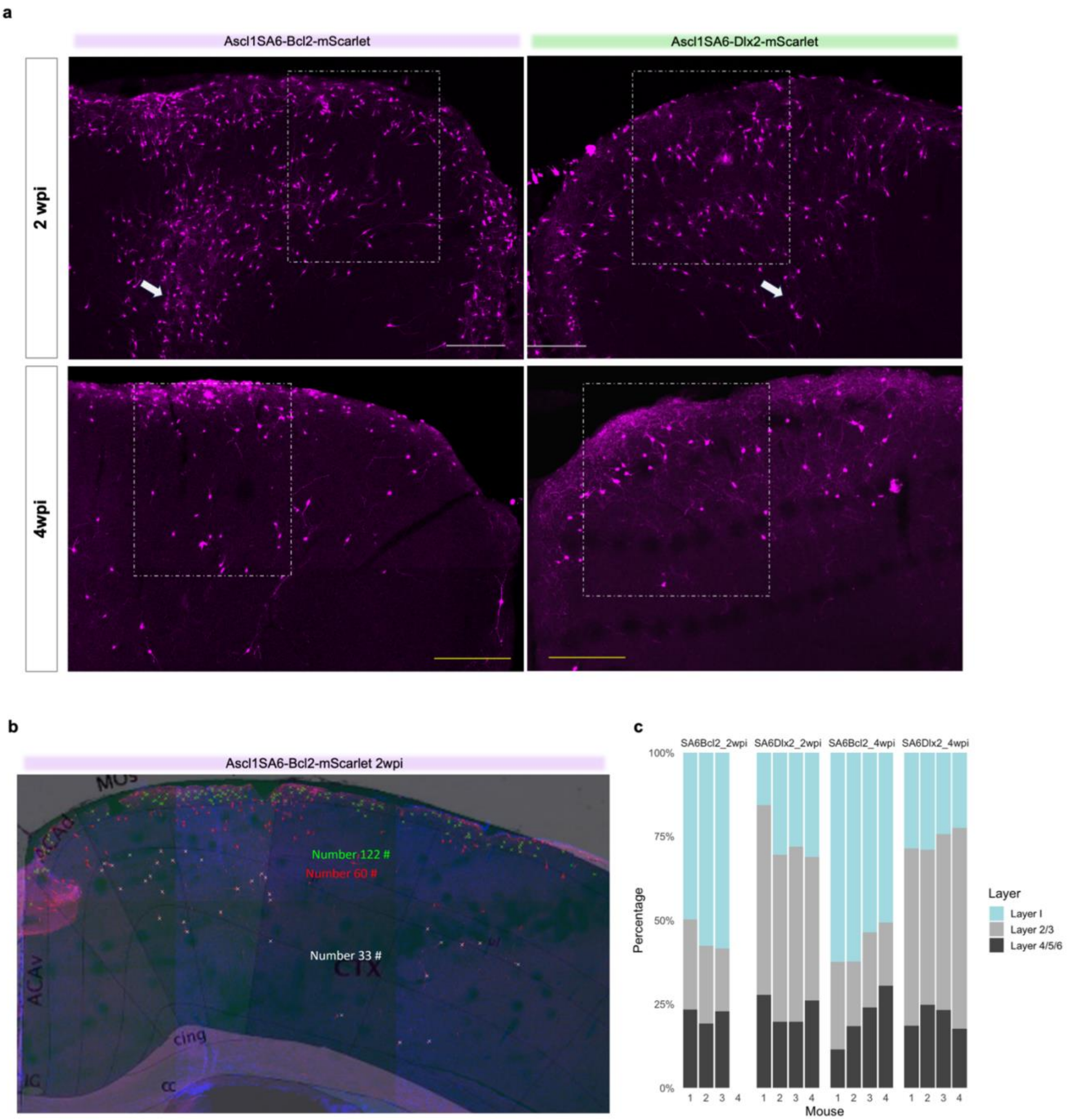

**Supplementary Figure 11. Cortical layer localization of iNs at 2 and 4 wpi following Ascl1SA6-Bcl2 or Ascl1SA6-Dlx2 reprogramming**

**a,** Representative images of Ascl1SA6-Bcl2 and Ascl1SA6-Dlx2-induced neurons (iNs) at 2 wpi (left) and 4 wpi (right). Dotted boxes indicate regions shown in Figure 7c,d and Extended Figure 10b. White arrows mark the approximate retroviral (RV) injection site. **b,** Example distribution of iNs across cortical layers in individual mice for Ascl1SA6-Bcl2 injections. Layer boundaries were defined using atlas.brain-map.org. **c,** Stacked bar plots showing laminar distribution of iNs per mouse at 2 and 4 wpi. Ascl1SA6-Bcl2 iNs are enriched in Layer 1, whereas Ascl1SA6-Dlx2 iNs predominantly localize to Layers 2/3.

**Supplementary Figure 12**

**Key:**
**Backbone:**
**CMV promoter**
**Retroviral Sequences:**
**5'LTR 3'LTR**
***MMLV PSI (CAPS)***
**GAG**

**CAGG promoter**
**[EcoRI site] not unique**
**IRES**
**H2B**
**mScarlet**
**WPRE**

RET29. pCAG ires H2B-mScarlet **WPRE**

gacattgattattgactagttattaatagtaatcaattacggggtcattagttcatagcccatatatggagttccgcgttacat
aacttacggtaaatggcccgcctggctgaccgcccaacgacccccgcccattgacgtcaataatgacgtatgttcccat
agtaacgccaatagggactttccattgacgtcaatgggtggagtatttacggtaaactgcccacttggcagtacatcaagt
gtatcatatgccaagtacgccccctattgacgtcaatgacggtaaatggcccgcctggcattatgcccagtacatgacct
tatgggactttcctacttggcagtacatctacgtattagtcatcgctattaccatggtgatgcggttttggcagtacatcaatg
ggcgtggatagcggtttgactcacggggatttccaagtctccaccccattgacgtcaatgggagtttgttttggcaccaaa
atcaacgggactttccaaaatgtcgtaacaactccgccccattgacgcaaatgggcggtaggcatgtacggtgggaggt
ctatataagcagagctcaataaaagagcccacaacccctcactcggggcgccagtcctccgattgactgagtcgcccg
ggtacccgtattcccaataaagcctcttgctgtttgcatccgaatcgtggtctcgctgttccttgggagggtctcctctgagt
gattgactacccacgacgggggtctttcatttgggggctcgtccgggatttggagacccctgcccagggaccaccgacc
caccaccgggaggt*AAGCTGGCCAGCAACTTATCTGTGTCTGTCCGATTGTCTAGTGTCTATGTTT*
*GATGTTATGCGCCTGCGTCTGTACTAGTTAGCTAACTAGCTCTGTATCTGGCGGACCCGTGGT*
*GGAACTGACGAGTTCTGAACACCCGGCCGCAACCCTGGGAGACGTCCCAGGGACTTTGG*
*GGGCCGTTTTTGTGGCCCGACCTGAG*gaagggagtcgatgtggaatccgaccccgtcaggatatgtggttct
ggtaggagacgagaacctaaaacagttcccgcctccgtctgaatttttgctttcggtttggaaccgaagccgcgcgtcttg
tctgctgcagcgctgcagcatcgttctgtgttgtctctgtctgactgtgtttctgtatttgtctgaaaattagggccagactgtta
ccactcccttaagtttgaccttaggtcactggaaagatgtcgagcggatcgctcacaaccagtcggtagatgtcaagaag
agacgttgggttaccttctgctctgcagaatggccaacctttaacgtcggatggccgcgagacggcacctttaaccgaga
cctcatcacccaggttaagatcaaggtcttttcacctggcccgcatggacacccagaccaggtcccctacatcgtgac
ctgggaagccttggcttttgacccccctccctgggtcaagccctttgtacaccctaagcctccgcctcctcttcctccatc
cgccccgtctctcccccttgaacctcctcgttcgaccccgcctcgatcctccctttatccagccctcactccttctctagg
cgccggaattcgatgtcgacattgattattgactagttattaatagtaatcaattacggggtcattagttcatagcccatatat
ggagttccgcgttacataacttacggtaaatggcccgcctggctgaccgcccaacgacccccgcccattgacgtcaat
aatgacgtatgttcccatagtaacgccaatagggactttccattgacgtcaatgggtggagtatttacggtaaactgccca
cttggcagtacatcaagtgtatcatatgccaagtacgccccctattgacgtcaatgacggtaaatggcccgcctggcatt

atgcccagtacatgaccttatgggactttcctacttggcagtacatctacgtattagtcatcgctattaccatggtcgaggtg
agccccacgttctgcttcactctccccatctcccccccctccccacccccaattttgtatttatttattttttaattattttgtg
cagcgatgggggcggggggggggggggggcccccccccaggcggggcggggcggggcgaggggcggggcggggcga
ggcggaaaggtgcggcggcagccaatcagagcggcgcgctccgaaagtttccttttatggcgaggcggcggcggcggc
ggccctataaaaagcgaagcgcgcggcgggcgggagtcgttgcgcgctgccttcgccccgtgccccgctccgccgcc
gcctcgcgccgcccgccccggctctgactgaccgcgttactcccacaggtgagcgggcgggacggcccttctcctccg
ggctgtaattagcgcttggtttaatgacggcttgtttcttttctgtggctgcgtgaaagccttgaggggctccgggagggccct
ttgtgcggggggagcggctcggggggtgcgtgcgtgtgtgtgtgcgtggggagcgccgcgtgcggctccgcgctgcccgg
cggctgtgagcgctgcgggcgcggcgcggggctttgtgcgctccgcagtgtgcgcgaggggagcgcggccgggggcggt
gccccgcggtgcggggggggctgcgaggggaacaaaggctgcgtgcggggtgtgtgcgtgggggggtgagcagggggt
gtgggcgcgtcggtcgggctgcaaccccccctgcacccccctccccgagttgctgagcacggcccggcttcgggtgcg
gggctccgtacggggcgtggcgcggggctcgccgtgccgggcggggggtggcggcaggtgggggtgccgggcggggcg
gggccgcctcgggccggggagggctcgggggaggggcgcggcggcccccggagcgccggcggctgtcgaggcgcg
gcgagccgcagccattgccttttatggtaatcgtgcgagagggcgcagggacttcctttgtcccaaatctgtgcggagccg
aaatctgggaggcgccgccgcaccccctctagcgggcgcggggcgaagcggtgcggcgccggcaggaaggaaatgg
gcggggagggccttcgtgcgtcgccgcgccgccgtccccttctccctctccagcctcggggctgtccgcggggggacgg
ctgccttcgggggggacggggcagggcggggttcggcttctggcgtgtgaccggcggctctagagcctctgctaaccatg
ttcatgccttcttctttttcctacagctcctgggcaacgtgctggttattgtgctgtctcatcattttggcaaa
**[gaattc] EcoRI**
gctagcggatcggccgatcacaagtttgtacaaaaaagcaggctttaaaggaaccaattcagtcgagctcaagcttcgt
cgagatatctagacccagctttcttgtacaaagtggtgataaacgatccgcccctctccctcccccccccctaacgtta
ctggccgaagccgcttggaataaggccggtgtgcgtttgtctatatgttattttccaccatattgccgtcttttggcaatgtga
gggcccggaaacctggccctgtcttcttgacgagcattcctaggggtctttcccctctcgccaaaggaatgcaaggtctgt
tgaatgtcgtgaaggaagcagttcctctggaagcttcttgaagacaaacaacgtctgtagcgaccctttgcaggcagcgg
aaccccccacctggcgacaggtgcctctgcggccaaaagccacgtgtataagatacacctgcaaaggcggcacaac
cccagtgccacgttgtgagttggatagttgtggaaagagtcaaatggctctcctcaagcgtattcaacaaggggctgaag
gatgcccagaaggtaccccattgtatgggatctgatctggggcctcggtacacatgctttacatgtgtttagtcgaggttaa
aaaaacgtctaggccccccgaaccacggggacgtggttttcctttgaaaaacacgatgataatatggccacaaccatg
ccagagccagcgaagtctgctcccgccccgaaaaagggctccaagaaggcggtgactaaggcgcagaagaaaggc
ggcaagaagcgcaagcgcagccgcaaggagagctattccatctatgtgtataaggttctgaagcaggtccaccctgac
actggcatttcgtccaaggctatgggcatcatgaactcgtttgtgaacgacattttcgagcgcatcgcaggtgaggcttcc
cgcctggcgcattacaacaagcgctcgaccatcacctccagggagatccagacggccgtgcgcctgctgctgcctggg
gagttggccaagcacgccgtgtccgagggtactaaggccatcaccaagtacaccagcgctaaggacccacctgtcgc
caccgctagcatggtgagcaagggcgaggcagtgatcaaggagttcatgcggttcaaggtgcacatggagggctccatg
aacggccacgagttcgagatcgagggcgagggcgagggccgcccctacgagggcacccagaccgccaagctgaag
gtgaccaagggtggccccctgcccttctcctgggacatcctgtcccctcagttcatgtacggctccagggccttcaccaa
gcaccccgccgacatccccgactactataagcagtccttccccgagggcttcaagtgggagcgcgtgatgaacttcga
ggacggcggcgccgtgaccgtgacccaggacacctccctggaggacggcaccctgatctacaaggtgaagctccgcg
gcaccaacttccctcctgacggccccgtaatgcagaagaagacaatgggctgggaagcgtccaccgagcggttgtac
cccgaggacggcgtgctgaagggcgacattaagatggccctgcgcctgaaggacggcgggcgctacctggcggacttc
aagaccacctacaaggccaagaagcccgtgcagatgcccggcgcctacaacgtggaccgcaagttggacatcacct
cccacaacgaggactacaccgtggtggaacagtacgaacgctccgagggccgccactccaccggcggcatggacga
gctgtataagtaagaattcgatatcaagcttatcgat

WPRE:
aatcaacctctggattacaaaatttgtgaaagattgactggtattcttaactatgttgctccttttacgctatgtggatacgct
gctttaatgcctttgtatcatgctattgcttcccgtatggctttcattttctcctccttgtataaatcctggttgctgtctctttatg

aggagttgtggcccgttgtcaggcaacgtggcgtggtgtgcactgtgtttgctgacgcaacccccactggttggggcattg
ccaccacctgtcagctcctttccgggactttcgctttccccctccctattgccacggcggaactcatcgccgcctgcctt
gcccgctgctggacaggggctcggctgttgggcactgacaattccgtggtgttgtcggggaaatcatcgtcctttccttggc
tgctcgcctgtgttgccacctggattctgcgcgggacgtccttctgctacgtcccttcggccctcaatccagcggaccttc
cttcccgcggcctgctgccggctctgcggcctcttccgcgtcttcgccttcgccctcagacgagtcggatctccctttggg
ccgcctccccgcgttaacatcgataaaataaaagattttatttagtctccagaaaaaggggggaatgaaagaccccacc
tgtaggtttggcaagctagcttaagtaacgccattttgcaaggcatggaaaaatacataactgagaatagagaagttcag
atcaaggtcaggaacagatggaacagctgaatatgggccaaacaggatatctgtggtaagcagttcctgccccggctca
gggccaagaacagatggaacagctgaatatgggccaaacaggatatctgtggtaagcagttcctgccccggctcaggg
ccaagaacagatggtccccagatgcggtccagccctcagcagtttctagagaaccatcagatgtttccagggtgcccc
aaggacctgaaatgaccctgtgccttatttgaactaaccaatcagttcgcttctcgcttctgttcgcgcgcttctgctcccc
gagctcaataaaagagcccacaacccctcactcggggcgccagtcctccgattgactgagtcgcccgggtacccgtgt
atccaataaaccctcttgcagttgcatccgacttgtggtctcgctgttccttgggagggtctcctctgagtgattgactaccc
gtcagcgggggtctttcacatgcagcatgtatcaaaattaatttggttttttttcttaagtatttacattaaatggccatagttgc
attaatgaatcggccaacgcgcggggagaggcggtttgcgtattggcgctcttccgcttcctcgctcactgactcgctgcg
ctcggtcgttcggctgcggcgagcggtatcagctcactcaaaggcggtaatacggttatccacagaatcaggggataac
gcaggaaagaacatgtgagcaaaaggccagcaaaaggccaggaaccgtaaaaaggccgcgttgctggcgtttttccat
aggctccgcccccctgacgagcatcacaaaaatcgacgctcaagtcagaggtggcgaaacccgacaggactataaa
gataccaggcgtttccccctggaagctccctcgtgcgctctcctgttccgaccctgccgcttaccggatacctgtccgcc
tttctcccttcgggaagcgtggcgctttctcatagctcacgctgtaggtatctcagttcggtgtaggtcgttcgctccaagct
gggctgtgtgcacgaaccccccgttcagcccgaccgctgcgccttatccggtaactatcgtcttgagtccaacccggta
agacacgacttatcgccactggcagcagccactggtaacaggattagcagagcgaggtatgtaggcggtgctacagag
ttcttgaagtggtggcctaactacggctacactagaaggacagtatttggtatctgcgctctgctgaagccagttaccttcg
gaaaaagagttggtagctcttgatccggcaaacaaaccaccgctggtagcggtggtttttttgtttgcaagcagcagatta
cgcgcagaaaaaaaggatctcaagaagatcctttgatcttttctacggggtctgacgctcagtggaacgaaaactcacgt
taagggattttggtcatgagattatcaaaaaggatcttcacctagatccttttgcggccggccgcaaatcaatctaaagtat
atatgagtaaacttggtctgacagttaccaatgcttaatcagtgaggcacctatctcagcgatctgtctatttcgttcatcca
tagttgcctgactccccgtcgtgtagataactacgatacgggagggcttaccatctggccccagtgctgcaatgataccg
cgagacccacgctcaccggctccagatttatcagcaataaaccagccagccggaagggccgagcgcagaagtggtc
ctgcaactttatccgcctccatccagtctattaattgttgccgggaagctagagtaagtagttcgccagttaatagtttgcgc
aacgttgttgccattgctacaggcatcgtggtgtcacgctcgtcgtttggtatggcttcattcagctccggttcccaacgatc
aaggcgagttacatgatcccccatgttgtgcaaaaaagcggttagctccttcggtcctccgatcgttgtcagaagtaagtt
ggccgcagtgttatcactcatggttatggcagcactgcataattctcttactgtcatgccatccgtaagatgcttttctgtga
ctggtgagtactcaaccaagtcattctgagaatagtgtatgcggcgaccgagttgctcttgcccggcgtcaacacgggat
aataccgcgccacatagcagaactttaaaagtgctcatcattggaaaacgttcttcggggcgaaaactctcaaggatctt
accgctgttgagatccagttcgatgtaacccactcgtgcacccaactgatcttcagcatcttttactttcaccagcgtttct
gggtgagcaaaaacaggaaggcaaaatgccgcaaaaaagggaataagggcgacacggaaatgttgaatactcatact
cttcctttttcaatgatccggccattagccatattattcattggttatatagcataaatcaatattggctattggccattgcata
cgttgtatccatatcataatatgtacatttatattggctcatgtccaacattaccgccatgtt

RET28- pCAG mASCl1 ires H2B-mScarlet WPRE
From EcoRI to WPRE

gctagcggatcggccgatcacaagtttgtacaaaaaagcaggctttaaaggaaccaattcagtcgagctcaagcttcga
agccaccatggagagctctggcaagatggagagtggagccggccagcagccgcagcccccgcagcccttcctgcctc

ccgcagcctgcttctttgcgaccgcggcggcggcagcggcggccgcggcagctcagagcgcgcagcagca
acagccgcaggcgccgccgcagcaggcgccgcagctgagcccggtggccgacagccagccctcaggggcggtca
caagtcagcggccaagcaggtcaagcgccagcgctcgtcctctccggaactgatgcgctgcaaacgccggctcaactt
cagcggcttcggctacagcctgccacagcagcagccggccgccgtggcgcgccgcaacgagcgcgagcgcaaccg
ggtcaagttggtcaacctgggctttgccaccctccgggagcatgtccccaacggcgcggccaacaagaagatgagcaa
ggtggagacgctgcgctcggcggtcgagtacatccgcgcgctgcagcagctgctggacgagcacgacgcggtgagcgc
tgcctttcaggcgggcgtcctgtcgcccaccatctcccccaactactccaacgacttgaactctatggcgggttctccgg
tctcgtcctactcctccgacgagggatcctacgaccctcttagcccagaggaacaagagctgctggactttaccaactg
gttctgaggacctgccaggctctcctgggaatggactttggaagcaggaattctgcagtcgacgaattcgcggccgcact
cgagatatctagacccagctttcttgtacaaagtggtgataaacgatccgcccctctccctcccccccccctaacgtta
ctggccgaagccgcttggaataaggccggtgtgcgtttgtctatatgttattttccaccatattgccgtcttttggcaatgtga
gggcccggaaacctggccctgtcttcttgacgagcattcctaggggtctttcccctctcgccaaaggaatgcaaggtctgt
tgaatgtcgtgaaggaagcagttcctctggaagcttcttgaagacaaacaacgtctgtagcgaccctttgcaggcagcgg
aaccccccacctggcgacaggtgcctctgcggccaaaagccacgtgtataagatacacctgcaaaggcggcacaac
cccagtgccacgttgtgagttggatagttgtggaaagagtcaaatggctctcctcaagcgtattcaacaaggggctgaag
gatgcccagaaggtaccccattgtatgggatctgatctggggcctcggtacacatgctttacatgtgtttagtcgaggttaa
aaaaacgtctaggccccccgaaccacggggacgtggttttcctttgaaaaacacgatgataatatggccacaaccatg
ccagagccagcgaagtctgctcccgccccgaaaaagggctccaagaaggcggtgactaaggcgcagaagaaaggc
ggcaagaagcgcaagcgcagccgcaaggagagctattccatctatgtgtataaggttctgaagcaggtccaccctgac
actggcatttcgtccaaggctatgggcatcatgaactcgtttgtgaacgacattttcgagcgcatcgcaggtgaggcttcc
cgcctggcgcattacaacaagcgctcgaccatcacctccagggagatccagacggccgtgcgcctgctgctgcctggg
gagttggccaagcacgccgtgtccgagggtactaaggccatcaccaagtacaccagcgctaaggacccacctgtcgc
caccgctagcatggtgagcaagggcgaggcagtgatcaaggagttcatgcggttcaaggtgcacatggagggctccatg
aacggccacgagttcgagatcgagggcgagggcgagggccgcccctacgagggcacccagaccgccaagctgaag
gtgaccaagggtggccccctgcccttctcctgggacatcctgtcccctcagttcatgtacggctccagggccttcaccaa
gcaccccgccgacatccccgactactataagcagtccttccccgagggcttcaagtgggagcgcgtgatgaacttcga
ggacggcggcgccgtgaccgtgacccaggacacctccctggaggacggcaccctgatctacaaggtgaagctccgcg
gcaccaacttccctcctgacggccccgtaatgcagaagaagacaatgggctgggaagcgtccaccgagcggttgtac
cccgaggacggcgtgctgaagggcgacattaagatggccctgcgcctgaaggacggcgggcgctacctggcggacttc
aagaccacctacaaggccaagaagcccgtgcagatgcccggcgcctacaacgtggaccgcaagttggacatcacct
cccacaacgaggactacaccgtggtggaacagtacgaacgctccgagggccgccactccaccggcggcatggacga
gctgtataagtaagaattcgatatcaagcttatcgat

RET14-pCAG-mASC1-SA6 ires H2B-mScarlet

gctagcggatcggccgatcacaagtttgtacaaaaaagcaggctttaaaggaaccaattcagtcgagctcaagcttcga
agccaccatggagagctctggcaagatggagagtggagccggccagcagccgcagcccccgcagcccttcctgcctc
ccgcagcctgcttctttgcgaccgcggcggcggcagcggcggccgcggcagctcagagcgcgcagcagca
acagccgcaggcgccgccgcagcaggcgccgcagctggccccggtggccgacagccagccctcaggggcggtca
caagtcagcggccaagcaggtcaagcgccagcgctcgtccgctccggaactgatgcgctgcaaacgccggctcaact
tcagcggcttcggctacagcctgccacagcagcagccggccgccgtggcgcgccgcaacgagcgcgagcgcaaccg
ggtcaagttggtcaacctgggctttgccaccctccgggagcatgtccccaacggcgcggccaacaagaagatgagcaa
ggtggagacgctgcgctcggcggtcgagtacatccgcgcgctgcagcagctgctggacgagcacgacgcggtgagcgc
tgcctttcaggcgggcgtcctggcgcccaccatcgcccccaactactccaacgacttgaactctatggcgggtgctccg

gtctcgtcctactcctccgacgagggatcctacgaccctcttgccccagaggaacaagagctgctggactttaccaact
ggttctgagaattctgcagtcgacgaattcgcggccgcactcgagatatctagacccagctttcttgtacaaagtggtgat
aaacgatccgcccctctccctcccccccccctaacgttactggccgaagccgcttggaataaggccggtgtgcgtttgt
ctatatgttattttccaccatattgccgtcttttggcaatgtgagggcccggaaacctggccctgtcttcttgacgagcattc
ctaggggtctttcccctctcgccaaaggaatgcaaggtctgttgaatgtcgtgaaggaagcagttcctctggaagcttcttg
aagacaaacaacgtctgtagcgaccctttgcaggcagcggaaccccccacctggcgacaggtgcctctgcggccaaa
agccacgtgtataagatacacctgcaaaggcggcacaaccccagtgccacgttgtgagttggatagttgtggaaagagt
caaatggctctcctcaagcgtattcaacaaggggctgaaggatgcccagaaggtaccccattgtatgggatctgatctgg
ggcctcggtacacatgctttacatgtgtttagtcgaggttaaaaaaacgtctaggccccccgaaccacggggacgtggtt
ttcctttgaaaaacacgatgataatatggccacaaccatgccagagccagcgaagtctgctcccgccccgaaaaaggg
ctccaagaaggcggtgactaaggcgcagaagaaaggcggcaagaagcgcaagcgcagccgcaaggagagctattcc
atctatgtgtataaggttctgaagcaggtccaccctgacactggcatttcgtccaaggctatgggcatcatgaactcgtttg
tgaacgacattttcgagcgcatcgcaggtgaggcttcccgcctggcgcattacaacaagcgctcgaccatcacctccag
ggagatccagacggccgtgcgcctgctgctgcctggggagttggccaagcacgccgtgtccgagggtactaaggccat
caccaagtacaccagcgctaaggacccacctgtcgccaccgctagcatggtgagcaagggcgaggcagtgatcaag
gagttcatgcggttcaaggtgcacatggagggctccatgaacggccacgagttcgagatcgagggcgagggcgagggc
cgcccctacgagggcacccagaccgccaagctgaaggtgaccaagggtggccccctgcccttctcctgggacatcct
gtcccctcagttcatgtacggctccagggccttcaccaagcaccccgccgacatccccgactactataagcagtcctt
ccccgagggcttcaagtgggagcgcgtgatgaacttcgaggacggcggcgccgtgaccgtgacccaggacacctccc
tggaggacggcaccctgatctacaaggtgaagctccgcggcaccaacttccctcctgacggccccgtaatgcagaag
aagacaatgggctgggaagcgtccaccgagcggttgtaccccgaggacggcgtgctgaagggcgacattaagatggcc
ctgcgcctgaaggacggcgggcgctacctggcggacttcaagaccacctacaaggccaagaagcccgtgcagatgcc
cggcgcctacaacgtggaccgcaagttggacatcacctcccacaacgaggactacaccgtggtggaacagtacgaa
cgctccgagggccgccactccaccggcggcatggacgagctgtataagtaagaattcgatatcaagcttatcgat

RET23 pCAG Flagx3 DLX2 ires H2B neon green

gctagcggatcggccgatcacaagtttgtacaaaaaagcaggctttaaaggaaccaattcagtcgagctcaagcttcga
agccaccatggactacaaggacgatgatgacaagggggactacaaagacgatgatgataagggagactataaggacg
acgacgataaagggggttccggaatgactggagtctttgacagtctggtggctgatatgcactcgacccagatcaccgcc
tccagcacgtaccaccagcaccagcagcccccgagcggtgcgggcgccggccctggcggcaacagcaacagcagc
agcagcaacagcagcctgcacaagccccaggagtcgccaaccctcccggtgtccacggctacggacagcagctact
acaccaaccagcagcacccggcgggcggcggcggcgggggggcctcgccctacgcgcacatgggctcctaccagta
ccacgccagcggcctcaacaatgtctcctactccgccaaaagcagctacgacctgggctacaccgccgcgtacacct
cctacgcgccctacggcaccagttcgtctccggtcaacaacgagccggacaaggaagaccttgagcctgaaatccga
atagtgaacgggaagccaaagaaagtccggaaaccacgcaccatctactccagtttccagctggcggcccttcaacg
acgcttccagaagacccagtatctggccctgccagagcgagccgagctggcggcgtccctgggcctcacccaaactc
aggtcaaaatctggttccagaaccgccgatccaagttcaagaagatgtggaaaagcggcgagatacccaccgagcag
caccctggagccagcgcttctcctccttgtgcctccccgccggtctcggcgccagcatcctgggacttcggcgcgccgc
agcggatggctggcggcggcccgggcagcggaggcggcggtgcgggcagctctggctccagcccgagcagcgccgc
ctcggcctttctgggaaactacccgtggtaccaccaggcttcgggctccgcttcacacctgcaggccacagcgccactt
ctgcatccttcgcagactccgcaggcgcaccatcaccaccatcaccaccaccacgcaggcgggggcgccccggtga
gcgcggggacgattttctaactcgagatatctagacccagctttcttgtacaaagtggtgataaacgatccgcccctctcc
ctcccccccccctaacgttactggccgaagccgcttggaataaggccggtgtgcgtttgtctatatgttattttccaccata
ttgccgtcttttggcaatgtgagggcccggaaacctggccctgtcttcttgacgagcattcctaggggtctttcccctctcg

ccaaaggaatgcaaggtctgttgaatgtcgtgaaggaagcagttcctctggaagcttcttgaagacaaacaacgtctgta
gcgaccctttgcaggcagcggaaccccccacctggcgacaggtgcctctgcggccaaaagccacgtgtataagatac
acctgcaaaggcggcacaaccccagtgccacgttgtgagttggatagttgtggaaagagtcaaatggctctcctcaagc
gtattcaacaaggggctgaaggatgcccagaaggtaccccattgtatgggatctgatctggggcctcggtacacatgcttt
acatgtgtttagtcgaggttaaaaaaacgtctaggccccccgaaccacggggacgtggttttcctttgaaaaacacgatg
ataatatggccacaaccatgccagagccagcgaagtctgctcccgccccgaaaaagggctccaagaaggcggtgact
aaggcgcagaagaaaggcggcaagaagcgcaagcgcagccgcaaggagagctattccatctatgtgtataaggttctg
aagcaggtccaccctgacactggcatttcgtccaaggctatgggcatcatgaactcgtttgtgaacgacattttcgagcg
catcgcaggtgaggcttcccgcctggcgcattacaacaagcgctcgaccatcacctccagggagatccagacggccgt
gcgcctgctgctgcctggggagttggccaagcacgccgtgtccgagggtactaaggccatcaccaagtacaccagcgc
taaggacccacctgtcgccaccgctagcatggtgagcaagggcgaggaggataacatggcctctctcccagcgacac
atgagttacacatctttggctccatcaacggtgtggactttgacatggtgggtcagggcaccggcaatccaaatgatggtt
atgaggagttaaacctgaagtccaccaagggtgacctccagttctccccctggattctggtccctcatatcgggtatggct
tccatcagtacctgccctaccctgacgggatgtcgcctttccaggccgccatggtagatggctccggataccaagtccat
cgcacaatgcagtttgaagatggtgcctcccttactgttaactaccgctacacctacgagggaagccacatcaaaggag
aggcccaggtgaaggggactggtttccctgctgacggtcctgtgatgaccaactcgctgaccgctgcggactggtgcag
gtcgaagaagacttaccccaacgacaaaaccatcatcagtacctttaagtggagttacaccactggaaatggcaagcg
ctaccggagcactgcgcggaccacctacacctttgccaagccaatggcggctaactatctgaagaaccagccgatgta
cgtgttccgtaagacggagctcaagcactccaagaccgagctcaacttcaaggagtggcaaaaggcctttaccgatgtg
atgggcatggacgagctgtacaagtaacaattgcgat

RET35-pCAG FLAG x 3 mASC1 SA6-T2A Dlx2--ires mScarlet

gctagcggatcggccgatcacaagtttgtacaaaaaagcaggctttaaaggaaccaattcagtcgagctcaagcttcga
agccaccatggactacaaggacgatgatgacaagggggactacaaagacgatgatgataagggagactataaggacg
acgacgataaagggggttccggaatggagagctctggcaagatggagagtggagccggccagcagccgcagccccg
cagcccttcctgcctcccgcagcctgcttctttgcgaccgcggcggcggcggcagcggcggcggccgcggcagctcag
agcgcgcagcagcaacagccgcaggcgccgccgcagcaggcgccgcagctgGCcccggtggccgacagccagc
cctcagggggcggtcacaagtcagcggccaagcaggtcaagcgccagcgctcgtccGctccggaactgatgcgctgc
aaacgccggctcaacttcagcggcttcggctacagcctgccacagcagcagccggccgccgtggcgcgccgcaacg
agcgcgagcgcaaccgggtcaagttggtcaacctgggctttgccaccctccgggagcatgtccccaacggcgcggcc
aacaagaagatgagcaaggtggagacgctgcgctcggcggtcgagtacatccgcgcgctgcagcagctgctggacga
gcacgacgcggtgagcgctgcctttcaggcgggcgtcctggcgcccaccatcgcccccaactactccaacgacttga
actctatggcgggtgctccggtctcgtcctactcctccgacgagggatcctacgaccctcttgccccagaggaacaaga
gctgctggactttaccaactggttcggaagcggagagggcaggggaagtcttctaacatgcggggacgtggaggaaaat
cccggccccatgactggagtctttgacagtctggtggctgatatgcactcgacccagatcaccgcctccagcacgtacc
accagcaccagcagcccccgagcggtgcgggcgccggccctggcggcaacagcaacagcagcagcagcaacagc
agcctgcacaagccccaggagtcgccaaccctcccggtgtccacggctacggacagcagctactacaccaaccagc
agcacccggcgggcggcggcggcgggggggcctcgccctacgcgcacatgggctcctaccagtaccacgccagcgg
cctcaacaatgtctcctactccgccaaaagcagctacgacctgggctacaccgccgcgtacacctcctacgcgccct
acggcaccagttcgtctccggtcaacaacgagccggacaaggaagaccttgagcctgaaatccgaatagtgaacggg
aagccaaagaaagtccggaaaccacgcaccatctactccagtttccagctggcggcccttcaacgacgcttccagaa
gacccagtatctggccctgccagagcgagccgagctggcggcgtccctgggcctcacccaaactcaggtcaaaatctg
gttccagaaccgccgatccaagttcaagaagatgtggaaaagcggcgagatacccaccgagcagcaccctggagcc
agcgcttctcctccttgtgcctccccgccggtctcggcgccagcatcctgggacttcggcgcgccgcagcggatggctg

gcggcggcccgggcagcggaggcggcggtgcgggcagctctggctccagcccgagcagcgccgcctcggcctttctg
ggaaactacccgtggtaccaccaggcttcgggctccgcttcacacctgcaggccacagcgccacttctgcatccttcg
cagactccgcaggcgcaccatcaccaccatcaccaccaccacgcaggcgggggcgccccggtgagcgcggggacg
attttctaactcgagatatctagacccagctttcttgtacaaagtggtgataaacgatccgcccctctccctcccccccc
ctaacgttactggccgaagccgcttggaataaggccggtgtgcgtttgtctatatgttattttccaccatattgccgtcttttg
gcaatgtgagggcccggaaacctggccctgtcttcttgacgagcattcctaggggtctttcccctctcgccaaaggaatg
caaggtctgttgaatgtcgtgaaggaagcagttcctctggaagcttcttgaagacaaacaacgtctgtagcgaccctttgc
aggcagcggaaccccccacctggcgacaggtgcctctgcggccaaaagccacgtgtataagatacacctgcaaagg
cggcacaaccccagtgccacgttgtgagttggatagttgtggaaagagtcaaatggctctcctcaagcgtattcaacaag
gggctgaaggatgcccagaaggtaccccattgtatgggatctgatctggggcctcggtacacatgctttacatgtgtttagt
cgaggttaaaaaaacgtctaggccccccgaaccacggggacgtggttttcctttgaaaaacacgatgataatatggcca
caacccgtacg
atggtgagcaagggcgaggcagtgatcaaggagttcatgcggttcaaggtgcacatggagggctccatgaacggccac
gagttcgagatcgagggcgagggcgagggccgcccctacgagggcacccagaccgccaagctgaaggtgaccaagg
gtggcccccctgcccttctcctgggacatcctgtcccctcagttcatgtacggctccagggccttcaccaagcaccccgc
cgacatccccgactactataagcagtccttccccgagggcttcaagtgggagcgcgtgatgaacttcgaggacggcgg
cgccgtgaccgtgacccaggacacctccctggaggacggcaccctgatctacaaggtgaagctccgcggcaccaact
tccctcctgacggccccgtaatgcagaagaagacaatgggctgggaagcgtccaccgagcggttgtaccccgaggac
ggcgtgctgaagggcgacattaagatggccctgcgcctgaaggacggcggccgctacctggcggacttcaagaccac
ctacaaggccaagaagcccgtgcagatgcccggcgcctacaacgtcgaccgcaagttggacatcacctcccacaac
gaggactacaccgtggtggaacagtacgaacgctccgagggccgccactccaccggcggcatggacgagctgtataa
gtaagaattcgatatcaagcttatcgat

RET36-pCAG FLAG x 3 mASC1-T2A Dlx2--ires mScarlet

gctagcggatcggccgatcacaagtttgtacaaaaaagcaggctttaaaggaaccaattcagtcgagctcaagcttcga
agccaccatggactacaaggacgatgatgacaagggggactacaaagacgatgatgataagggagactataaggacg
acgacgataaaggggttccggaatggagagctctggcaagatggagagtggagccggccagcagccgcagccccg
cagcccttcctgcctcccgcagcctgcttctttgcgaccgcggcggcggcggcagcggcggcggccgcggcagctcag
agcgcgcagcagcaacagccgcaggcgccgccgcagcaggcgccgcagctgagcccggtggccgacagccagcc
ctcagggggcggtcacaagtcagcggccaagcaggtcaagcgccagcgctcgtcctctccggaactgatgcgctgca
aacgccggctcaacttcagcggcttcggctacagcctgccacagcagcagccggccgccgtggcgcgccgcaacga
gcgcgagcgcaaccgggtcaagttggtcaacctgggctttgccaccctccgggagcatgtccccaacggcgcggcca
acaagaagatgagcaaggtggagacgctgcgctcggcggtcgagtacatccgcgcgctgcagcagctgctggacgag
cacgacgcggtgagcgctgcctttcaggcgggcgtcctgtcgcccaccatctcccccaactactccaacgacttgaac
tctatggcgggttctccggtctcgtcctactcctccgacgagggatcctacgaccctcttagcccagaggaacaagagct
gctggactttaccaactggttcggaagcggagagggcaggggaagtcttctaacatgcggggacgtggaggaaaatccc
ggccccatgactggagtctttgacagtctggtggctgatatgcactcgacccagatcaccgcctccagcacgtaccacc
agcaccagcagcccccgagcggtgcgggcgccggccctggcggcaacagcaacagcagcagcagcaacagcagc
ctgcacaagccccaggagtcgccaaccctcccggtgtccacggctacggacagcagctactacaccaaccagcagc
acccggcgggcggcggcggcgggggggcctcgccctacgcgcacatgggctcctaccagtaccacgccagcggcct
caacaatgtctcctactccgccaaaagcagctacgacctgggctacaccgccgcgtacacctcctacgcgccctacg
gcaccagttcgtctccggtcaacaacgagccggacaaggaagaccttgagcctgaaatccgaatagtgaacgggaag
ccaaagaaagtccggaaaccacgcaccatctactccagtttccagctggcggcccttcaacgacgcttccagaagac
ccagtatctggccctgccagagcgagccgagctggcggcgtccctgggcctcacccaaactcaggtcaaaatctggtt
ccagaaccgccgatccaagttcaagaagatgtggaaaagcggcgagatacccaccgagcagcaccctggagccagc

gcttctcctccttgtgcctccccgccggtctcggcgccagcatcctgggacttcggcgcgccgcagcggatggctggcgg
cggcccgggcagcggaggcggcggtgcgggcagctctggctccagcccgagcagcgccgcctcggcctttctgggaa
actacccgtggtaccaccaggcttcgggctccgcttcacacctgcaggccacagcgccacttctgcatccttcgcaga
ctccgcaggcgcaccatcaccaccatcaccaccaccacgcaggcgggggcgccccggtgagcgcggggacgatttt
ctaactcgagatatctagacccagctttcttgtacaaagtggtgataaacgatccgcccctctccctcccccccccta
acgttactggccgaagccgcttggaataaggccggtgtgcgtttgtctatatgttattttccaccatattgccgtcttttggca
atgtgagggcccggaaacctggccctgtcttcttgacgagcattcctaggggtctttcccctctcgccaaaggaatgcaa
ggtctgttgaatgtcgtgaaggaagcagttcctctggaagcttcttgaagacaaacaacgtctgtagcgaccctttgcagg
cagcggaaccccccacctggcgacaggtgcctctgcggccaaaagccacgtgtataagatacacctgcaaaggcgg
cacaaccccagtgccacgttgtgagttggatagttgtggaaagagtcaaatggctctcctcaagcgtattcaacaagggg
ctgaaggatgcccagaaggtaccccattgtatgggatctgatctggggcctcggtacacatgctttacatgtgtttagtcga
ggttaaaaaaacgtctaggccccccgaaccacggggacgtggttttcctttgaaaaacacgatgataatatggccacaa
cccgtacgatggtgagcaagggcgaggcagtgatcaaggagttcatgcggttcaaggtgcacatggagggctccatgaa
cggccacgagttcgagatcgagggcgagggcgagggccgcccctacgagggcacccagaccgccaagctgaaggtg
accaagggtggcccccctgcccttctcctgggacatcctgtcccctcagttcatgtacggctccagggccttcaccaagc
accccgccgacatccccgactactataagcagtccttccccgagggcttcaagtgggagcgcgtgatgaacttcgagg
acggcggcgccgtgaccgtgacccaggacacctccctggaggacggcaccctgatctacaaggtgaagctccgcggc
accaacttccctcctgacggccccgtaatgcagaagaagacaatgggctgggaagcgtccaccgagcggttgtaccc
cgaggacggcgtgctgaagggcgacattaagatggccctgcgcctgaaggacggcggccgctacctggcggacttcaa
gaccacctacaaggccaagaagcccgtgcagatgcccggcgcctacaacgtcgaccgcaagttggacatcacctcc
cacaacgaggactacaccgtggtggaacagtacgaacgctccgagggccgccactccaccggcggcatggacgagc
tgtataagtaagaattcgatatcaagcttatcgat

RET46 pCAG mScarlet

gctagcggatcggccgatcacaagtttgtacaaaaaagcaggctttaaaggaaccaattcagtcgagctcaagcttccc
acaacccgtacgatggtgagcaagggcgaggcagtgatcaaggagttcatgcggttcaaggtgcacatggagggctcc
atgaacggccacgagttcgagatcgagggcgagggcgagggccgcccctacgagggcacccagaccgccaagctga
aggtgaccaagggtggcccccctgcccttctcctgggacatcctgtcccctcagttcatgtacggctccagggccttcacc
aagcaccccgccgacatccccgactactataagcagtccttccccgagggcttcaagtgggagcgcgtgatgaacttc
gaggacggcggcgccgtgaccgtgacccaggacacctccctggaggacggcaccctgatctacaaggtgaagctccg
cggcaccaacttccctcctgacggccccgtaatgcagaagaagacaatgggctgggaagcgtccaccgagcggttgta
ccccgaggacggcgtgctgaagggcgacattaagatggccctgcgcctgaaggacggcggccgctacctggcggactt
caagaccacctacaaggccaagaagcccgtgcagatgcccggcgcctacaacgtcgaccgcaagttggacatcacc
tcccacaacgaggactacaccgtggtggaacagtacgaacgctccgagggccgccactccaccggcggcatggacg
agctgtataagtaagaattcgatatcaagcttatcgat

RET62-pCAG FLAG x 3 mASC1 SA6-ires mScarlet

gctagcggatcggccgatcacaagtttgtacaaaaaagcaggctttaaaggaaccaattcagtcgagctcaagcttcga
agccaccatggactacaaggacgatgatgacaagggggactacaaagacgatgatgataagggagactataaggacg
acgacgataaagggggttccggaatggagagctctggcaagatggagagtggagccggccagcagccgcagccccg
cagcccttcctgcctcccgcagcctgcttctttgcgaccgcggcggcggcagcggcggccgcggcagctcag
agcgcgcagcagcaacagccgcaggcgccgccgcagcaggcgccgcagctggccccggtggccgacagccagcc

ctcagggggcggtcacaagtcagcggccaagcaggtcaagcgccagcgctcgtccgctccggaactgatgcgctgca
aacgccggctcaacttcagcggcttcggctacagcctgccacagcagcagccggccgccgtggcgcgccgcaacga
gcgcgagcgcaaccgggtcaagttggtcaacctgggctttgccaccctccgggagcatgtccccaacggcgcggcca
acaagaagatgagcaaggtggagacgctgcgctcggcggtcgagtacatccgcgcgctgcagcagctgctggacgag
cacgacgcggtgagcgctgcctttcaggcgggcgtcctggcgcccaccatcgcccccaactactccaacgacttgaa
ctctatggcgggtgctccggtctcgtcctactcctccgacgagggatcctacgaccctcttgccccagaggaacaagag
ctgctggactttaccaactggttctgagaattctgcagtcgacgaattcgcggccgcactcgagatatctagacccagctt
tcttgtacaaagtggtgataaacgatccgcccctctccctcccccccccctaacgttactggccgaagccgcttggaat
aaggccggtgtgcgtttgtctatatgttattttccaccatattgccgtcttttggcaatgtgagggcccggaaacctggccct
gtcttcttgacgagcattcctaggggtctttcccctctcgccaaaggaatgcaaggtctgttgaatgtcgtgaaggaagca
gttcctctggaagcttcttgaagacaaacaacgtctgtagcgaccctttgcaggcagcggaaccccccacctggcgac
aggtgcctctgcggccaaaagccacgtgtataagatacacctgcaaaggcggcacaaccccagtgccacgttgtgagt
tggatagttgtggaaagagtcaaatggctctcctcaagcgtattcaacaaggggctgaaggatgcccagaaggtacccc
attgtatgggatctgatctggggcctcggtacacatgctttacatgtgtttagtcgaggttaaaaaacgtctaggccccc
gaaccacggggacgtggttttcctttgaaaaacacgatgataatatggccacaacccgtacgatggtgagcaagggcga
ggcagtgatcaaggagttcatgcggttcaaggtgcacatggagggctccatgaacggccacgagttcgagatcgagggc
gagggcgagggccgcccctacgagggcacccagaccgccaagctgaaggtgaccaagggtggccccctgcccttct
cctgggacatcctgtcccctcagttcatgtacggctccagggccttcaccaagcaccccgccgacatccccgactact
ataagcagtccttccccgagggcttcaagtgggagcgcgtgatgaacttcgaggacggcggcgccgtgaccgtgaccc
aggacacctccctggaggacggcaccctgatctacaaggtgaagctccgcggcaccaacttccctcctgacggcccc
gtaatgcagaagaagacaatgggctgggaagcgtccaccgagcggttgtaccccgaggacggcgtgctgaagggcga
cattaagatggccctgcgcctgaaggacggcggccgctacctggcggacttcaagaccacctacaaggccaagaagc
ccgtgcagatgcccggcgcctacaacgtcgaccgcaagttggacatcacctcccacaacgaggactacaccgtggtg
gaacagtacgaacgctccgagggccgccactccaccggcggcatggacgagctgtataagtaagaattcgatatcaag
cttatcgat

RET65-pCAG FLAG x 3 mASC1 SA6 T2A Bcl2-ires mScarlet

Gctagcggatcggccgatcacaagtttgtacaaaaaagcaggctttaaggaaccaattcagtcgagctcaagcttcg
aagccaccatggactacaaggacgatgatgacaaggggactacaaagacgatgatgataagggagactataaggac
gacgacgataaagggggttccgga
atggagagctctggcaagatggagagtggagccggccagcagccgcagcccccgcagcccttcctgcctcccgcagc
ctgcttctttgcgaccgcggcggcggcggcagcggcggcggccgcggcagctcagagcgcgcagcagcaacagccg
caggcgccgccgcagcaggcgccgcagctggccccggtggccgacagccagccctcagggggcggtcacaagtca
gcggccaagcaggtcaagcgccagcgctcgtccgctccggaactgatgcgctgcaaacgccggctcaacttcagcgg
cttcggctacagcctgccacagcagcagccggccgccgtggcgcgccgcaacgagcgcgagcgcaaccgggtcaag
ttggtcaacctgggctttgccaccctccgggagcatgtccccaacggcgcggccaacaagaagatgagcaaggtggag
acgctgcgctcggcggtcgagtacatccgcgcgctgcagcagctgctggacgagcacgacgcggtgagcgctgccttt
caggcgggcgtcctggcgcccaccatcgcccccaactactccaacgacttgaactctatggcgggtgctccggtctcgt
cctactcctccgacgagggatcctacgaccctcttgccccagaggaacaagagctgctggactttaccaactggttcgg
aagcggagagggcaggggaagtcttctaacatgcggggacgtggaggaaaatcccggccccatggcgcacgctggga
gaacagggtacgataaccgggagatagtgatgaagtacatccattataagctgtcgcagaggggctacgagtgggatgc
gggagatgtgggcgccgcgcccccgggggccgcccccgcaccgggcatcttctcctcccagcccgggcacacgccc
catccagccgcatcccgggacccggtcgccaggacctcgccgctgcagaccccggctgcccccggcgccgccgcg
gggcctgcgctcagcccggtgccacctgtggtccacctgaccctccgccaggccggcgacgacttctcccgccgctac
cgccgcgacttcgccgagatgtccagccagctgcacctgacgcccttcaccgcgcggggacgctttgccacggtggtg

gaggagctcttcagggacggggtgaactggggaggattgtggccttctttgagttcggtggggtcatgtgtgtggagagcg
tcaaccgggagatgtcgcccctggtggacaacatcgccctgtggatgactgagtacctgaaccggcacctgcacacct
ggatccaggataacggaggctgggatgcctttgtggaactgtacggccccagcatgcggcctctgtttgatttctcctggct
gtctctgaagactctgctcagtttggccctggtgggagcttgcatcaccctgggtgcctatctgggccacaagtgactcga
gatatctagacccagctttcttgtacaaagtggtgataaacgatccgcccctctccctcccccccccctaacgttactgg
ccgaagccgcttggaataaggccggtgtgcgtttgtctatatgttattttccaccatattgccgtcttttggcaatgtgagggc
ccggaaacctggccctgtcttcttgacgagcattcctaggggtctttcccctctcgccaaaggaatgcaaggtctgttgaa
tgtcgtgaaggaagcagttcctctggaagcttcttgaagacaaacaacgtctgtagcgaccctttgcaggcagcggaac
cccccacctggcgacaggtgcctctgcggccaaaagccacgtgtataagatacacctgcaaaggcggcacaacccc
agtgccacgttgtgagttggatagttgtggaaagagtcaaatggctctcctcaagcgtattcaacaaggggctgaaggatg
cccagaaggtaccccattgtatgggatctgatctggggcctcggtacacatgctttacatgtgtttagtcgaggttaaaaaa
acgtctaggccccccgaaccacggggacgtggttttcctttgaaaaacacgatgataatatggccacaacccgtacgat
ggtgagcaagggcgaggcagtgatcaaggagttcatgcggttcaaggtgcacatggagggctccatgaacggccacga
gttcgagatcgagggcgagggcgagggccgcccctacgagggcacccagaccgccaagctgaaggtgaccaagggt
ggcccctgcccttctcctgggacatcctgtcccctcagttcatgtacggctccagggccttcaccaagcaccccgccg
acatccccgactactataagcagtccttccccgagggcttcaagtgggagcgcgtgatgaacttcgaggacggcggcg
ccgtgaccgtgacccaggacacctccctggaggacggcaccctgatctacaaggtgaagctccgcggcaccaacttc
cctcctgacggccccgtaatgcagaagaagacaatgggctgggaagcgtccaccgagcggttgtaccccgaggacgg
cgtgctgaagggcgacattaagatggccctgcgcctgaaggacggcggccgctacctggcggacttcaagaccaccta
caaggccaagaagcccgtgcagatgcccggcgcctacaacgtcgaccgcaagttggacatcacctcccacaacgag
gactacaccgtggtggaacagtacgaacgctccgagggccgccactccaccggcggcatggacgagctgtataagta
agaattcgatatcaagcttatcgat

RET15-pCAG ires H2B mScarlet

gctagcggatcggccgatcacaagtttgtacaaaaaagcaggctttaaaggaaccaattcagtcgagctcaagcttcgt
cgagatatctagacccagctttcttgtacaaagtggtgataaacgatccgcccctctccctcccccccccctaacgtta
ctggccgaagccgcttggaataaggccggtgtgcgtttgtctatatgttattttccaccatattgccgtcttttggcaatgtga
gggcccggaaacctggccctgtcttcttgacgagcattcctaggggtctttcccctctcgccaaaggaatgcaaggtctgt
tgaatgtcgtgaaggaagcagttcctctggaagcttcttgaagacaaacaacgtctgtagcgaccctttgcaggcagcgg
aaccccccacctggcgacaggtgcctctgcggccaaaagccacgtgtataagatacacctgcaaaggcggcacaac
cccagtgccacgttgtgagttggatagttgtggaaagagtcaaatggctctcctcaagcgtattcaacaaggggctgaag
gatgcccagaaggtaccccattgtatgggatctgatctggggcctcggtacacatgctttacatgtgtttagtcgaggttaa
aaaaacgtctaggccccccgaaccacggggacgtggttttcctttgaaaaacacgatgataatatggccacaaccatg
ccagagcatgataatatggccacaaccatgccagagccagcgaagtctgctcccgccccgaaaaagggctccaaga
aggcggtgactaaggcgcagaagaaaggcggcaagaagcgcaagcgcagccgcaaggagagctattccatctatgtg
tataaggttctgaagcaggtccaccctgacactggcatttcgtccaaggctatgggcatcatgaactcgtttgtgaacgac
attttcgagcgcatcgcaggtgaggcttcccgcctggcgcattacaacaagcgctcgaccatcacctccagggagatcc
agacggccgtgcgcctgctgctgcctggggagttggccaagcacgccgtgtccgagggtactaaggccatcaccaagt
acaccagcgctaaggacccacctgtcgccaccgctagcatggtgagcaagggcgaggcagtgatcaaggagttcatg
cggttcaaggtgcacatggagggctccatgaacggccacgagttcgagatcgagggcgagggcgagggccgcccccta
cgagggcacccagaccgccaagctgaaggtgaccaagggtggcccctgcccttctcctgggacatcctgtcccctc
agttcatgtacggctccagggccttcaccaagcaccccgccgacatccccgactactataagcagtccttccccgagg
gcttcaagtgggagcgcgtgatgaacttcgaggacggcggcgccgtgaccgtgacccaggacacctccctggaggacg
gcaccctgatctacaaggtgaagctccgcggcaccaacttccctcctgacggccccgtaatgcagaagaagacaatg
ggctgggaagcgtccaccgagcggttgtaccccgaggacggcgtgctgaagggcgacattaagatggccctgcgcctg

aaggacggcgggcgctacctggcggacttcaagaccacctacaaggccaagaagcccgtgcagatgcccggcgcct
acaacgtggaccgcaagttggacatcacctcccacaacgaggactacaccgtggtggaacagtacgaacgctccga
gggccgccactccaccggcggcatggacgagctgtataagtaagaattcgatatcaagcttatcgat

RET19-pCAG FLAG x 3 mASC1 SA6 -ires H2B mScarlet

gctagcggatcggccgatcacaagtttgtacaaaaaagcaggctttaaaggaaccaattcagtcgagctcaagcttcga
agccaccatggactacaaggacgatgatgacaagggggactacaaagacgatgatgataagggagactataaggacg
acgacgataaagggggttccggaatggagagctctggcaagatggagagtggagccggccagcagccgcagcccccg
cagcccttcctgcctcccgcagcctgcttctttgcgaccgcggcggcggcggcagcggcggcggccgcggcagctcag
agcgcgcagcagcaacagccgcaggcgccgccgcagcaggcgccgcagctggccccggtggccgacagccagcc
ctcagggggcggtcacaagtcagcggccaagcaggtcaagcgccagcgctcgtccgctccggaactgatgcgctgca
aacgccggctcaacttcagcggcttcggctacagcctgccacagcagcagccggccgccgtggcgcgccgcaacga
gcgcgagcgcaaccgggtcaagttggtcaacctgggctttgccaccctccgggagcatgtccccaacggcgcggcca
acaagaagatgagcaaggtggagacgctgcgctcggcggtcgagtacatccgcgcgctgcagcagctgctggacgag
cacgacgcggtgagcgctgcctttcaggcgggcgtcctggcgcccaccatcgcccccaactactccaacgacttgaa
ctctatggcgggtgctccggtctcgtcctactcctccgacgagggatcctacgaccctcttgccccagaggaacaagag
ctgctggactttaccaactggttctgagaattctgcagtcgacgaattcgcggccgcactcgagatatctagacccagctt
tcttgtacaaagtggtgataaacgatccgcccctctccctcccccccccctaacgttactggccgaagccgcttggaat
aaggccggtgtgcgtttgtctatatgttattttccaccatattgccgtcttttggcaatgtgagggcccggaaacctggccct
gtcttcttgacgagcattcctaggggtctttcccctctcgccaaaggaatgcaaggtctgttgaatgtcgtgaaggaagca
gttcctctggaagcttcttgaagacaaacaacgtctgtagcgaccctttgcaggcagcggaaccccccacctggcgac
aggtgcctctgcggccaaaagccacgtgtataagatacacctgcaaaggcggcacaaccccagtgccacgttgtgagt
tggatagttgtggaaagagtcaaatggctctcctcaagcgtattcaacaaggggctgaaggatgcccagaaggtacccc
attgtatgggatctgatctggggcctcggtacacatgctttacatgtgtttagtcgaggttaaaaaacgtctaggccccc
gaaccacggggacgtggttttcctttgaaaaacacgatgataatatggccacaaccatgccagagccagcgaagtctg
ctcccgccccgaaaaagggctccaagaaggcggtgactaaggcgcagaagaaaggcggcaagaagcgcaagcgca
gccgcaaggagagctattccatctatgtgtataaggttctgaagcaggtccaccctgacactggcatttcgtccaaggct
atgggcatcatgaactcgtttgtgaacgacattttcgagcgcatcgcaggtgaggcttcccgcctggcgcattacaacaa
gcgctcgaccatcacctccagggagatccagacggccgtgcgcctgctgctgcctggggagttggccaagcacgccgt
gtccgagggtactaaggccatcaccaagtacaccagcgctaaggacccacctgtcgccaccgctagcatggtgagca
agggcgaggcagtgatcaaggagttcatgcggttcaaggtgcacatggagggctccatgaacggccacgagttcgagat
cgagggcgagggcgagggccgcccctacgagggcacccagaccgccaagctgaaggtgaccaagggtggcccct
gcccttctcctgggacatcctgtcccctcagttcatgtacggctccagggccttcaccaagcaccccgccgacatcccc
gactactataagcagtccttccccgagggcttcaagtgggagcgcgtgatgaacttcgaggacggcggcgccgtgaccg
tgacccaggacacctccctggaggacggcaccctgatctacaaggtgaagctccgcggcaccaacttccctcctgac
ggccccgtaatgcagaagaagacaatgggctgggaagcgtccaccgagcggttgtaccccgaggacggcgtgctgaa
gggcgacattaagatggccctgcgcctgaaggacggcgggcgctacctggcggacttcaagaccacctacaaggcca
agaagcccgtgcagatgcccggcgcctacaacgtggaccgcaagttggacatcacctcccacaacgaggactacac
cgtggtggaacagtacgaacgctccgagggccgccactccaccggcggcatggacgagctgtataagtaagaattcga
tatcaagcttatcgat

RET13- pCAG mASC1 ires H2B mScarlet
gctagcggatcggccgatcacaagtttgtacaaaaaagcaggctttaaaggaaccaattcagtcgagctcaagcttcga
agccaccatggagagctctggcaagatggagagtggagccggccagcagccgcagcccccgcagcccttcctgcctc
ccgcagcctgcttctttgcgaccgcggcggcggcggcagcggcggcggccgcggcagctcagagcgcgcagcagca

acagccgcaggcgccgccgcagcaggcgccgcagctgagcccggtggccgacagccagccctcagggggcggtca
caagtcagcggccaagcaggtcaagcgccagcgctcgtcctctccggaactgatgcgctgcaaacgccggctcaactt
cagcggcttcggctacagcctgccacagcagcagccggccgccgtggcgcgccgcaacgagcgcgagcgcaaccg
ggtcaagttggtcaacctgggctttgccaccctccgggagcatgtccccaacggcgcggccaacaagaagatgagcaa
ggtggagacgctgcgctcggcggtcgagtacatccgcgcgctgcagcagctgctggacgagcacgacgcggtgagcgc
tgcctttcaggcgggcgtcctgtcgcccaccatctcccccaactactccaacgacttgaactctatggcgggttctccgg
tctcgtcctactcctccgacgagggatcctacgaccctcttagcccagaggaacaagagctgctggactttaccaactg
gttctgaggacctgccaggctctcctgggaatggactttggaagcaggaattctgcagtcgacgaattcgcggccgcact
cgagatatctagacccagctttcttgtacaaagtggtgataaacgatccgcccctctccctcccccccccctaacgtta
ctggccgaagccgcttggaataaggccggtgtgcgtttgtctatatgttattttccaccatattgccgtcttttggcaatgtga
gggcccggaaacctggccctgtcttcttgacgagcattcctaggggtctttcccctctcgccaaaggaatgcaaggtctgt
tgaatgtcgtgaaggaagcagttcctctggaagcttcttgaagacaaacaacgtctgtagcgaccctttgcaggcagcgg
aaccccccacctggcgacaggtgcctctgcggccaaaagccacgtgtataagatacacctgcaaaggcggcacaac
cccagtgccacgttgtgagttggatagttgtggaaagagtcaaatggctctcctcaagcgtattcaacaaggggctgaag
gatgcccagaaggtaccccattgtatgggatctgatctggggcctcggtacacatgctttacatgtgtttagtcgaggttaa
aaaaacgtctaggccccccgaaccacggggacgtggttttcctttgaaaaacacgatgataatatggccacaaccatg
ccagagccagcgaagtctgctcccgccccgaaaaagggctccaagaaggcggtgactaaggcgcagaagaaaggc
ggcaagaagcgcaagcgcagccgcaaggagagctattccatctatgtgtataaggttctgaagcaggtccaccctgac
actggcatttcgtccaaggctatgggcatcatgaactcgtttgtgaacgacattttcgagcgcatcgcaggtgaggcttcc
cgcctggcgcattacaacaagcgctcgaccatcacctccagggagatccagacggccgtgcgcctgctgctgcctggg
gagttggccaagcacgccgtgtccgagggtactaaggccatcaccaagtacaccagcgctaaggacccacctgtcgc
caccgctagcatggtgagcaagggcgaggcagtgatcaaggagttcatgcggttcaaggtgcacatggagggctccatg
aacggccacgagttcgagatcgagggcgagggcgagggccgcccctacgagggcacccagaccgccaagctgaag
gtgaccaagggtggccccctgcccttctcctgggacatcctgtcccctcagttcatgtacggctccagggccttcaccaa
gcaccccgccgacatccccgactactataagcagtccttccccgagggcttcaagtgggagcgcgtgatgaacttcga
ggacggcggcgccgtgaccgtgacccaggacacctccctggaggacggcaccctgatctacaaggtgaagctccgcg
gcaccaacttccctcctgacggccccgtaatgcagaagaagacaatgggctgggaagcgtccaccgagcggttgtac
cccgaggacggcgtgctgaagggcgacattaagatggccctgcgcctgaaggacggcggccgctacctggcggacttc
aagaccacctacaaggccaagaagcccgtgcagatgcccggcgcctacaacgtcgaccgcaagttggacatcacct
cccacaacgaggactacaccgtggtggaacagtacgaacgctccgagggccgccactccaccggcggcatggacga
gctgtataagtaagaattcgatatcaagcttatcgat

RET18- pCAG FLAG x 3 mASC1 ires H2B mScarlet

gctagcggatcggccgatcacaagtttgtacaaaaaagcaggctttaaaggaaccaattcagtcgagctcaagcttcga
agccaccatggactacaaggacgatgatgacaagggggactacaaagacgatgatgataagggagactataaggacg
acgacgataaagggggttccggaatggagagctctggcaagatggagagtggagccggccagcagccgcagccccg
cagcccttcctgcctcccgcagcctgcttctttgcgaccgcggcggcggcggcagcggcggcggccgcggcagctcag
agcgcgcagcagcaacagccgcaggcgccgccgcagcaggcgccgcagctgagcccggtggccgacagccagcc
ctcagggggcggtcacaagtcagcggccaagcaggtcaagcgccagcgctcgtcctctccggaactgatgcgctgca
aacgccggctcaacttcagcggcttcggctacagcctgccacagcagcagccggccgccgtggcgcgccgcaacga
gcgcgagcgcaaccgggtcaagttggtcaacctgggctttgccaccctccgggagcatgtccccaacggcgcggcca
acaagaagatgagcaaggtggagacgctgcgctcggcggtcgagtacatccgcgcgctgcagcagctgctggacgag
cacgacgcggtgagcgctgcctttcaggcgggcgtcctgtcgcccaccatctcccccaactactccaacgacttgaac
tctatggcgggttctccggtctcgtcctactcctccgacgagggatcctacgaccctcttagcccagaggaacaagagct
gctggactttaccaactggttctgaggacctgccaggctctcctgggaatggactttggaagcaggaattctgcagtcgac

gaattcgcggccgcactcgagatatctagacccagctttcttgtacaaagtggtgataaacgatccgcccctctccctcc
cccccccctaacgttactggccgaagccgcttggaataaggccggtgtgcgtttgtctatatgttattttccaccatattgc
cgtcttttggcaatgtgagggcccggaaacctggccctgtcttcttgacgagcattcctaggggtctttcccctctcgccaa
aggaatgcaaggtctgttgaatgtcgtgaaggaagcagttcctctggaagcttcttgaagacaaacaacgtctgtagcga
ccctttgcaggcagcggaaccccccacctggcgacaggtgcctctgcggccaaaagccacgtgtataagatacacct
gcaaaggcggcacaaccccagtgccacgttgtgagttggatagttgtggaaagagtcaaatggctctcctcaagcgtatt
caacaaggggctgaaggatgcccagaaggtaccccattgtatgggatctgatctggggcctcggtacacatgctttacat
gtgtttagtcgaggttaaaaaaacgtctaggccccccgaaccacggggacgtggttttcctttgaaaaacacgatgataa
tatggccacaaccatgccagagccagcgaagtctgctcccgccccgaaaaagggctccaagaaggcggtgactaag
gcgcagaagaaaggcggcaagaagcgcaagcgcagccgcaaggagagctattccatctatgtgtataaggttctgaag
caggtccaccctgacactggcatttcgtccaaggctatgggcatcatgaactcgtttgtgaacgacattttcgagcgcatc
gcaggtgaggcttcccgcctggcgcattacaacaagcgctcgaccatcacctccagggagatccagacggccgtgcg
cctgctgctgcctggggagttggccaagcacgccgtgtccgagggtactaaggccatcaccaagtacaccagcgctaa
ggacccacctgtcgccaccgctagcatggtgagcaagggcgaggcagtgatcaaggagttcatgcggttcaaggtgca
catggagggctccatgaacggccacgagttcgagatcgagggcgagggcgagggccgcccctacgagggcacccag
accgccaagctgaaggtgaccaagggtggcccccctgcccttctcctgggacatcctgtcccctcagttcatgtacggct
ccagggccttcaccaagcaccccgccgacatccccgactactataagcagtccttccccgagggcttcaagtgggag
cgcgtgatgaacttcgaggacggcggcgccgtgaccgtgacccaggacacctccctggaggacggcaccctgatcta
caaggtgaagctccgcggcaccaacttccctcctgacggccccgtaatgcagaagaagacaatgggctgggaagcgt
ccaccgagcggttgtaccccgaggacggcgtgctgaagggcgacattaagatggccctgcgcctgaaggacggcggc
cgctacctggcggacttcaagaccacctacaaggccaagaagcccgtgcagatgcccggcgcctacaacgtcgacc
gcaagttggacatcacctcccacaacgaggactacaccgtggtggaacagtacgaacgctccgagggccgccactcc
accggcggcatggacgagctgtataagtaagaattcgatatcaagcttatcgat

plasmid 113-pCAG mASC1 ires dsRED
gctagcggatcggccgatcacaagtttGTACAAAAAAGCAGGCTTTAAAGGAACCAATTCAGtcgagCT
CAAGCTTCGAATTCGCGTCCCCTTCTCGTTCTCCCCCGCGACAGTTTGGCCCGGCatggaga
gctctggcaagatggagagtggagccggccagcagccgcagcccccgcagcccttcctgcctcccgcagcctgcttct
ttgcgaccgcggcggcggcggcagcggcggcggccgcggcagctcagagcgcgcagcagcaacagccgcaggcgc
cgccgcagcaggcgccgcagctgagcccggtggccgacagccagccctcagggggcggtcacaagtcagcggcca
agcaggtcaagcgccagcgctcgtcctctccggaactgatgcgctgcaaacgccggctcaacttcagcggcttcggct
acagcctgccacagcagcagccggccgccgtggcgcgccgcaacgagcgcgagcgcaaccgggtcaagttggtcaa
cctgggctttgccaccctccgggagcatgtccccaacggcgcggccaacaagaagatgagcaaggtggagacgctgc
gctcggcggtcgagtacatccgcgcgctgcagcagctgctggacgagcacgacgcggtgagcgctgcctttcaggcgg
gcgtcctgtcgcccaccatctcccccaactactccaacgacttgaactctatggcgggttctccggtctcgtcctactcc
tccgacgagggatcctacgaccctcttagcccagaggaacaagagctgctggactttaccaactggttctgaGGACC
TGCCAGGCTCTCCTGGGAATGGACTTTGGAAGCAGGAATTCTGCAGTCGACGAATTCGCGG
CCGCACTCGAGATATCTAGACCCAGCTTTcttgtacaaagtggtgataaacgatccgcccctctccctccc
ccccccctaacgttactggccgaagccgcttggaataaggccggtgtgcgtttgtctatatgttattttccaccatattgcc
gtcttttggcaatgtgagggcccggaaacctggccctgtcttcttgacgagcattcctaggggtctttcccctctcgccaaa
ggaatgcaaggtctgttgaatgtcgtgaaggaagcagttcctctggaagcttcttgaagacaaacaacgtctgtagcgac
cctttgcaggcagcggaaccccccacctggcgacaggtgcctctgcggccaaaagccacgtgtataagatacacctg
caaaggcggcacaaccccagtgccacgttgtgagttggatagttgtggaaagagtcaaatggctctcctcaagcgtattc
aacaaggggctgaaggatgcccagaaggtaccccattgtatgggatctgatctggggcctcggtacacatgctttacatg
tgtttagtcgaggttaaaaaaacgtctaggccccccgaaccacggggacgtggttttcctttgaaaaacacgatgataat

atggccacaaccATGGCCTCCTCCGAGGACGTCATCAAGGAGTTCATGCGCTTCAAGGTGCGC
ATGGAGGGCTCCGTGAACGGCCACGAGTTCGAGATCGAGGGCGAGGGCGAGGGCCGCC
CCTACGAGGGCACCCAGACCGCCAAGCTGAAGGTGACCAAGGGCGGCCCCCTGCCCTT
CGCCTGGGACATCCTGTCCCCCCAGTTCCAGTACGGCTCCAAGGTGTACGTGAAGCACCC
CGCCGACATCCCCGACTACAAGAAGCTGTCCTTCCCCGAGGGCTTCAAGTGGGAGCGCG
TGATGAACTTCGAGGACGGCGGCGTGGTGACCGTGACCCAGGACTCCTCCCTGCAGGAC
GGCTGCTTCATCTACAAGGTGAAGTTCATCGGCGTGAACTTCCCCTCCGACGGCCCCGTAA
TGCAGAAGAAGACTATGGGCTGGGAGCCCTCCACCGAGCGCCTGTACCCCCGCGACGG
CGTGCTGAAGGGCGAGATCCACAAGGCCCTGAAGCTGAAGGACGGCGGCCACTACCTGG
TGGAGTTCAAGTCCATCTACATGGCCAAGAAGCCCGTGCAGCTGCCCGGCTACTACTACGT
GGACTCCAAGCTGGACATCACCTCCCACAACGAGGACTACACCATCGTGGAGCAGTACGA
GCGCGCCGAGGGCCGCCACCACCTGTTCCTGTAGcggccgcgtcga

plasmid 114-pCAG mASC1 SA6 ires dsRED
gctagcggatcggccgatcacaagtttGTACAAAAAAGCAGGCTTTAAAGGAACCAATTCAGtcgagatg
gagagctctggcaagatggagagtggagccggccagcagccgcagcccccgcagcccttcctgcctcccgcagcctg
cttctttgcgaccgcggcggcggcagcggcggcggccgcggcagctcagagcgcgcagcagcaacagccgcag
gcgccgccgcagcaggcgccgcagctggccccggtggccgacagccagccctcagggggcggtcacaagtcagcg
gccaagcaggtcaagcgccagcgctcgtccgctccggaactgatgcgctgcaaacgccggctcaacttcagcggcttc
ggctacagcctgccacagcagcagccggccgccgtggcgcgccgcaacgagcgcgagcgcaaccgggtcaagttgg
tcaacctgggctttgccaccctccgggagcatgtccccaacggcgcggccaacaagaagatgagcaaggtggagacg
ctgcgctcggcggtcgagtacatccgcgcgctgcagcagctgctggacgagcacgacgcggtgagcgctgcctttcagg
cgggcgtcctggcgcccaccatcgcccccaactactccaacgacttgaactctatggcgggtgctccggtctcgtccta
ctcctccgacgagggatcctacgaccctcttgccccagaggaacaagagctgctggactttaccaactggttctgaGA
ATTCTGCAGTCGACGAATTCGCGGCCGCACTCGAGATATCTAGACCCAGCTTTcttgtacaaagt
ggtgataaacgatccgcccctctccctcccccccccctaacgttactggccgaagccgcttggaataaggccggtgtg
cgtttgtctatatgttattttccaccatattgccgtcttttggcaatgtgagggcccggaaacctggccctgtcttcttgacga
gcattcctaggggtctttcccctctcgccaaaggaatgcaaggtctgttgaatgtcgtgaaggaagcagttcctctggaag
cttcttgaagacaaacaacgtctgtagcgaccctttgcaggcagcggaaccccccacctggcgacaggtgcctctgcg
gccaaaagccacgtgtataagatacacctgcaaaggcggcacaaccccagtgccacgttgtgagttggatagttgtgga
aagagtcaaatggctctcctcaagcgtattcaacaaggggctgaaggatgcccagaaggtaccccattgtatgggatct
gatctggggcctcggtacacatgctttacatgtgtttagtcgaggttaaaaaacgtctaggccccccgaaccacgggga
cgtggttttcctttgaaaacacgatgataatatggccacaaccATGGCCTCCTCCGAGGACGTCATCAAG
GAGTTCATGCGCTTCAAGGTGCGCATGGAGGGCTCCGTGAACGGCCACGAGTTCGAGATC
GAGGGCGAGGGCGAGGGCCGCCCCTACGAGGGCACCCAGACCGCCAAGCTGAAGGTG
ACCAAGGGCGGCCCCCTGCCCTTCGCCTGGGACATCCTGTCCCCCCAGTTCCAGTACGG
CTCCAAGGTGTACGTGAAGCACCCCGCCGACATCCCCGACTACAAGAAGCTGTCCTTCCC
CGAGGGCTTCAAGTGGGAGCGCGTGATGAACTTCGAGGACGGCGGCGTGGTGACCGTGA
CCCAGGACTCCTCCCTGCAGGACGGCTGCTTCATCTACAAGGTGAAGTTCATCGGCGTGAA
CTTCCCCTCCGACGGCCCCGTAATGCAGAAGAAGACTATGGGCTGGGAGCCCTCCACCG
AGCGCCTGTACCCCCGCGACGGCGTGCTGAAGGGCGAGATCCACAAGGCCCTGAAGCT
GAAGGACGGCGGCCACTACCTGGTGGAGTTCAAGTCCATCTACATGGCCAAGAAGCCCGT
GCAGCTGCCCGGCTACTACTACGTGGACTCCAAGCTGGACATCACCTCCCACAACGAGGA
CTACACCATCGTGGAGCAGTACGAGCGCGCCGAGGGCCGCCACCACCTGTTCCTGTAGcg
gccgcgtcgac

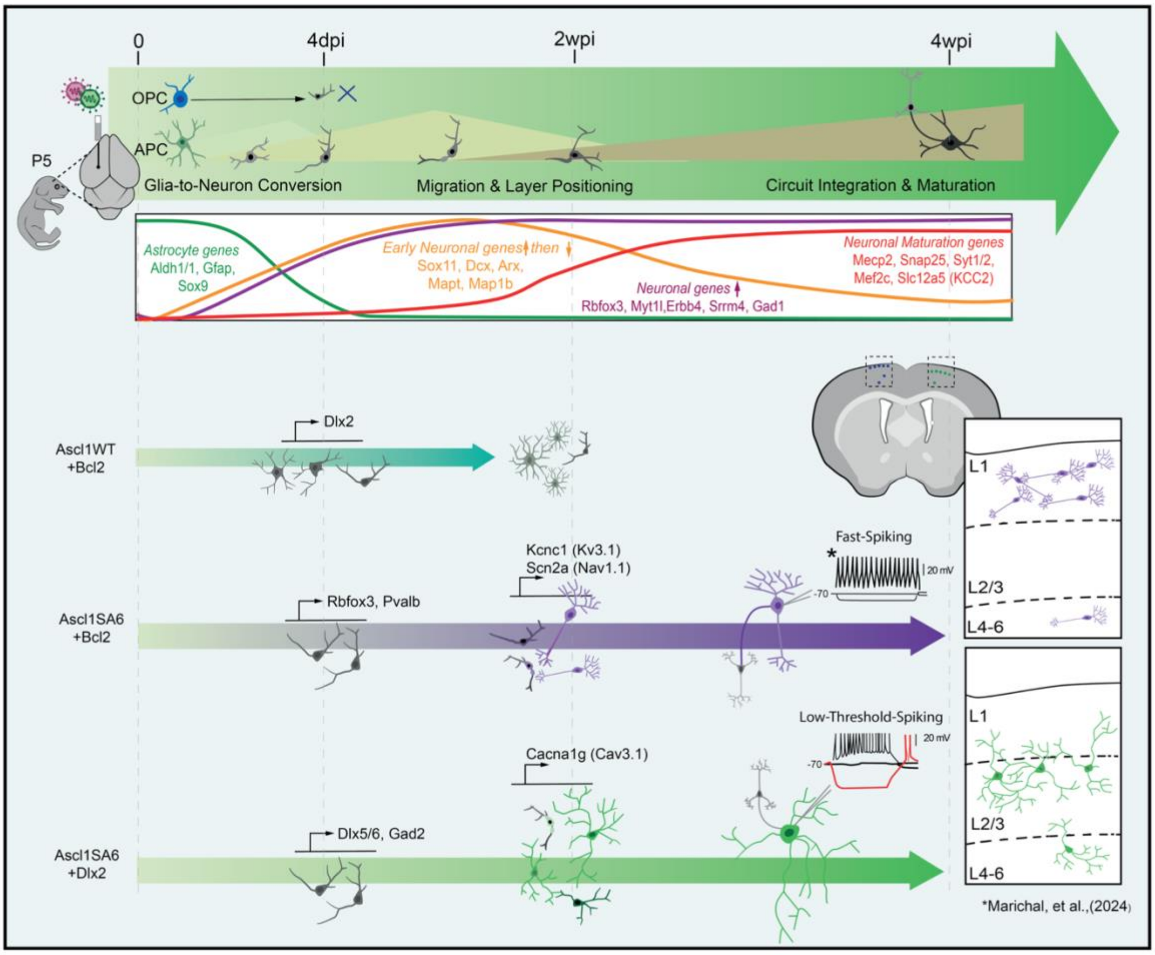

**Supplementary Figure 13: Overview of experimental design, key findings, and lineage reprogramming outcomes.** Schematic summarizing the study, including experimental design, retroviral conditions, glial-to-neuron reprogramming strategies, and key molecular and functional outcomes observed across different time points and lineages.

**Table 1.**

**Cell type proportions in each RV sample at 4 dpi derived for astrocytes versus OPCs.**

1a

| cell.type | Control | Ascl1WT | Ascl1WT/Bcl2 | Ascl1SA6 | Ascl1SA6/Bcl2 |
|---|---|---|---|---|---|
| Astrocytes | 46 50% | 8 60% | 0 60% | 5 40% | 0 60% |
| Cycling Astro | 0 30% | 14 00% | 8 80% | 7 90% | 2 40% |
| Astro-L | 0 20% | 22 00% | 16 90% | 3 00% | 0 80% |
| Interm-1 (Astro) | 0 10% | 6 40% | 15 60% | 11 70% | 8 30% |
| E-Neuronal | NA | 3 20% | 18 50% | 10 30% | 13 80% |
| Adv-Neuronal-1 | NA | 2 00% | 17 20% | 31 90% | 63 60% |
| Combined E-and Adv | | 5 20% | 35 70% | 42 20% | 77 40% |
| OPCs | 26 70% | 6 00% | 1 00% | 1 90% | 0 40% |
| Cycling OPCs | 5 10% | 7 00% | 1 90% | 0 30% | NA |
| Interm-2 (OPCs) | 1 20% | 15 60% | 9 80% | 1 90% | 0 10% |
| Adv-Neuronal-2 | NA | 0 60% | 1 00% | 9 50% | 4 80% |
| OL | 7 80% | 2 90% | 2 50% | 0 40% | NA |
| Microglia | 11 50% | 11 50% | 5 90% | 15 60% | 5 10% |
| Pericytes | 0 50% | 0 20% | 0 30% | 0 20% | 0 10% |

**combined for astrocytes verus OPC-devived Neuronal clusters**

1b

| Cell Type | Ascl1WT | Ascl1WT/Bcl2 | Ascl1SA6 | Ascl1SA6/Bcl2 |
|---|---|---|---|---|
| Astro-Neuronal combined | 5 20% | 35 70% | 42 20% | 77 40% |
| OPC-Neuronal | 0 60% | 1 00% | 9 50% | 4 80% |

**each condition so that Astro-Neuronal + OPC-Neuronal = 100%**

1c

| cell.type | Ascl1WT | Ascl1WT/Bcl2 | Ascl1SA6 | Ascl1SA6/Bcl2 | Mean across samples | SD (%) |
|---|---|---|---|---|---|---|
| Astro-Neuronal | 89 60% | 97 30% | 81 60% | 94 20% | 90 68% | 6 83% |
| OPC-Neuronal | 10 40% | 2 70% | 18 40% | 5 80% | 9 33% | 6 83% |

Table 2.

Ascl1_module for up and downregulted Ascl1 targets all genes combined

| Module | RV | group1 | group2 | statistic | p | p.signif |
|---|---|---|---|---|---|---|
| UP | Ascl1WT | O2N | A2N | 478510 | 0 522 | ns |
| UP | Ascl1WTBcl2 | O2N | A2N | 49952 | 4 09E-30 | **** |
| UP | Ascl1SA6 | O2N | A2N | 227106 | 7 18E-31 | **** |
| UP | Ascl1SA6Bcl2 | O2N | A2N | 29720 | 6 34E-22 | **** |
| DOWN | Ascl1WT | O2N | A2N | 690969 | 5 86E-68 | **** |
| DOWN | Ascl1WTBcl2 | O2N | A2N | 187075 | 4 01E-72 | **** |
| DOWN | Ascl1SA6 | O2N | A2N | 632504 | 2 31E-113 | **** |
| DOWN | Ascl1SA6Bcl2 | O2N | A2N | 138881 | 2 07E-48 | **** |

**Table 3.**

Module Score for up and downregulted Ascl1 targets per Clusters

| RV | Glia | mean | sd | n | p | stars | label | module |
|---|---|---|---|---|---|---|---|---|
| Control | O2N | -0 071388998 | 0 06162806 | 503 | 2 10E-151 | **** | **** 2.10e-151 | Ascl1_targets_cluster1_features |
| Control | A2N | -0 203414948 | 0 049496978 | 633 | 2 10E-151 | **** | **** 2.10e-151 | Ascl1_targets_cluster1_features |
| Ascl1WT | O2N | 0 002460935 | 0 090641616 | 700 | 1 11E-51 | **** | **** 1.11e-51 | Ascl1_targets_cluster1_features |
| Ascl1WT | A2N | -0 071159027 | 0 112632406 | 1339 | 1 11E-51 | **** | **** 1.11e-51 | Ascl1_targets_cluster1_features |
| Ascl1WTBcl2 | O2N | 0 023508157 | 0 091307902 | 192 | 0 220609648 | ns | ns 2.21e-01 | Ascl1_targets_cluster1_features |
| Ascl1WTBcl2 | A2N | 0 014616868 | 0 106321149 | 1068 | 0 220609648 | ns | ns 2.21e-01 | Ascl1_targets_cluster1_features |
| Ascl1SA6 | O2N | 0 105577858 | 0 122897721 | 375 | 1 55E-05 | **** | **** 1.55e-05 | Ascl1_targets_cluster1_features |
| Ascl1SA6 | A2N | 0 067073127 | 0 14206015 | 1821 | 1 55E-05 | **** | **** 1.55e-05 | Ascl1_targets_cluster1_features |
| Ascl1SA6Bcl2 | O2N | 0 150641959 | 0 109884426 | 93 | 0 31063847 | ns | ns 3.11e-01 | Ascl1_targets_cluster1_features |
| Ascl1SA6Bcl2 | A2N | 0 142873501 | 0 111671889 | 1423 | 0 31063847 | ns | ns 3.11e-01 | Ascl1_targets_cluster1_features |
| Control | O2N | -0 241527408 | 0 039779961 | 503 | 1 91E-137 | **** | **** 1.91e-137 | Ascl1_targets_cluster2_features |
| Control | A2N | -0 156918551 | 0 041809632 | 633 | 1 91E-137 | **** | **** 1.91e-137 | Ascl1_targets_cluster2_features |
| Ascl1WT | O2N | -0 196259248 | 0 064358577 | 700 | 6 75E-139 | **** | **** 6.75e-139 | Ascl1_targets_cluster2_features |
| Ascl1WT | A2N | -0 101032986 | 0 077403421 | 1339 | 6 75E-139 | **** | **** 6.75e-139 | Ascl1_targets_cluster2_features |
| Ascl1WTBcl2 | O2N | -0 172431938 | 0 068612569 | 192 | 4 94E-76 | **** | **** 4.94e-76 | Ascl1_targets_cluster2_features |
| Ascl1WTBcl2 | A2N | -0 001085033 | 0 097552468 | 1068 | 4 94E-76 | **** | **** 4.94e-76 | Ascl1_targets_cluster2_features |
| Ascl1SA6 | O2N | -0 03779686 | 0 118414461 | 375 | 2 96E-59 | **** | **** 2.96e-59 | Ascl1_targets_cluster2_features |
| Ascl1SA6 | A2N | 0 088965192 | 0 13456795 | 1821 | 2 96E-59 | **** | **** 2.96e-59 | Ascl1_targets_cluster2_features |
| Ascl1SA6Bcl2 | O2N | 0 037834704 | 0 109805859 | 93 | 1 85E-28 | **** | **** 1.85e-28 | Ascl1_targets_cluster2_features |
| Ascl1SA6Bcl2 | A2N | 0 179353804 | 0 103317994 | 1423 | 1 85E-28 | **** | **** 1.85e-28 | Ascl1_targets_cluster2_features |
| Control | O2N | -0 204722527 | 0 058332334 | 503 | 1 13E-23 | **** | **** 1.13e-23 | Ascl1_targets_cluster3_features |
| Control | A2N | -0 240105003 | 0 056521911 | 633 | 1 13E-23 | **** | **** 1.13e-23 | Ascl1_targets_cluster3_features |
| Ascl1WT | O2N | -0 115357657 | 0 098130025 | 700 | 2 14E-07 | **** | **** 2.14e-07 | Ascl1_targets_cluster3_features |
| Ascl1WT | A2N | -0 138153191 | 0 107266433 | 1339 | 2 14E-07 | **** | **** 2.14e-07 | Ascl1_targets_cluster3_features |
| Ascl1WTBcl2 | O2N | -0 086275287 | 0 102592703 | 192 | 1 52E-16 | **** | **** 1.52e-16 | Ascl1_targets_cluster3_features |
| Ascl1WTBcl2 | A2N | -0 007382705 | 0 127896778 | 1068 | 1 52E-16 | **** | **** 1.52e-16 | Ascl1_targets_cluster3_features |
| Ascl1SA6 | O2N | 0 031951228 | 0 136989922 | 375 | 9 30E-12 | **** | **** 9.30e-12 | Ascl1_targets_cluster3_features |
| Ascl1SA6 | A2N | 0 078460548 | 0 165721291 | 1821 | 9 30E-12 | **** | **** 9.30e-12 | Ascl1_targets_cluster3_features |
| Ascl1SA6Bcl2 | O2N | 0 120264676 | 0 143393184 | 93 | 5 73E-05 | **** | **** 5.73e-05 | Ascl1_targets_cluster3_features |
| Ascl1SA6Bcl2 | A2N | 0 176819924 | 0 104988223 | 1423 | 5 73E-05 | **** | **** 5.73e-05 | Ascl1_targets_cluster3_features |
| Control | O2N | -0 060704171 | 0 067877781 | 503 | 2 51E-11 | **** | **** 2.51e-11 | Ascl1_targets_cluster4_features |
| Control | A2N | -0 031118946 | 0 070660696 | 633 | 2 51E-11 | **** | **** 2.51e-11 | Ascl1_targets_cluster4_features |
| Ascl1WT | O2N | -0 040832749 | 0 089690359 | 700 | 9 27E-35 | **** | **** 9.27e-35 | Ascl1_targets_cluster4_features |
| Ascl1WT | A2N | -0 089652409 | 0 093607819 | 1339 | 9 27E-35 | **** | **** 9.27e-35 | Ascl1_targets_cluster4_features |
| Ascl1WTBcl2 | O2N | -0 035742698 | 0 094362999 | 192 | 0 475943201 | ns | ns 4.76e-01 | Ascl1_targets_cluster4_features |
| Ascl1WTBcl2 | A2N | -0 023862841 | 0 140958993 | 1068 | 0 475943201 | ns | ns 4.76e-01 | Ascl1_targets_cluster4_features |
| Ascl1SA6 | O2N | 0 035568869 | 0 109486373 | 375 | 0 046787593 | ** | ** 4.68e-02 | Ascl1_targets_cluster4_features |
| Ascl1SA6 | A2N | 0 030138555 | 0 157034602 | 1821 | 0 046787593 | ** | ** 4.68e-02 | Ascl1_targets_cluster4_features |
| Ascl1SA6Bcl2 | O2N | 0 084847505 | 0 098752292 | 93 | 0 204744878 | ns | ns 2.05e-01 | Ascl1_targets_cluster4_features |
| Ascl1SA6Bcl2 | A2N | 0 093281214 | 0 140634282 | 1423 | 0 204744878 | ns | ns 2.05e-01 | Ascl1_targets_cluster4_features |

| | | | | | | | | |
|---|---|---|---|---|---|---|---|---|
| Control | O2N | 0 001118201 | 0 046606526 | 503 | 7 55E-112 | **** | **** 7.55e-112 | Ascl1_targets_cluster5_features |
| Control | A2N | 0 090979514 | 0 058373994 | 633 | 7 55E-112 | **** | **** 7.55e-112 | Ascl1_targets_cluster5_features |
| Ascl1WT | O2N | -0 004608449 | 0 05022786 | 700 | 8 38E-69 | **** | **** 8.38e-69 | Ascl1_targets_cluster5_features |
| Ascl1WT | A2N | 0 049214758 | 0 068334276 | 1339 | 8 38E-69 | **** | **** 8.38e-69 | Ascl1_targets_cluster5_features |
| Ascl1WTBcl2 | O2N | -0 006141282 | 0 045889353 | 192 | 0 876397897 | ns | ns 8.76e-01 | Ascl1_targets_cluster5_features |
| Ascl1WTBcl2 | A2N | -0 002910302 | 0 063654713 | 1068 | 0 876397897 | ns | ns 8.76e-01 | Ascl1_targets_cluster5_features |
| Ascl1SA6 | O2N | 0 002221319 | 0 055468554 | 375 | 5 30E-17 | **** | **** 5.30e-17 | Ascl1_targets_cluster5_features |
| Ascl1SA6 | A2N | -0 022106435 | 0 069337553 | 1821 | 5 30E-17 | **** | **** 5.30e-17 | Ascl1_targets_cluster5_features |
| Ascl1SA6Bcl2 | O2N | -0 00423198 | 0 048685526 | 93 | 1 80E-17 | **** | **** 1.80e-17 | Ascl1_targets_cluster5_features |
| Ascl1SA6Bcl2 | A2N | -0 049200531 | 0 045622774 | 1423 | 1 80E-17 | **** | **** 1.80e-17 | Ascl1_targets_cluster5_features |
| Control | O2N | 0 047585388 | 0 073713333 | 503 | 6 89E-149 | **** | **** 6.89e-149 | Ascl1_targets_cluster6_features |
| Control | A2N | 0 255790949 | 0 100873757 | 633 | 6 89E-149 | **** | **** 6.89e-149 | Ascl1_targets_cluster6_features |
| Ascl1WT | O2N | 0 012447553 | 0 097911882 | 700 | 7 54E-54 | **** | **** 7.54e-54 | Ascl1_targets_cluster6_features |
| Ascl1WT | A2N | 0 104225916 | 0 129821171 | 1339 | 7 54E-54 | **** | **** 7.54e-54 | Ascl1_targets_cluster6_features |
| Ascl1WTBcl2 | O2N | -0 006947094 | 0 093520646 | 192 | 0 64510037 | ns | ns 6.45e-01 | Ascl1_targets_cluster6_features |
| Ascl1WTBcl2 | A2N | -0 009873143 | 0 106674002 | 1068 | 0 64510037 | ns | ns 6.45e-01 | Ascl1_targets_cluster6_features |
| Ascl1SA6 | O2N | -0 012924193 | 0 097387514 | 375 | 1 36E-09 | **** | **** 1.36e-09 | Ascl1_targets_cluster6_features |
| Ascl1SA6 | A2N | -0 03447834 | 0 13829932 | 1821 | 1 36E-09 | **** | **** 1.36e-09 | Ascl1_targets_cluster6_features |
| Ascl1SA6Bcl2 | O2N | -0 0525166 | 0 104727989 | 93 | 8 03E-07 | **** | **** 8.03e-07 | Ascl1_targets_cluster6_features |
| Ascl1SA6Bcl2 | A2N | -0 105147596 | 0 084046166 | 1423 | 8 03E-07 | **** | **** 8.03e-07 | Ascl1_targets_cluster6_features |
| Control | O2N | 0 028681724 | 0 051175993 | 503 | 4 01E-26 | **** | **** 4.01e-26 | Ascl1_targets_cluster7_features |
| Control | A2N | -0 005279466 | 0 051466947 | 633 | 4 01E-26 | **** | **** 4.01e-26 | Ascl1_targets_cluster7_features |
| Ascl1WT | O2N | 0 016671564 | 0 062041745 | 700 | 0 732683455 | ns | ns 7.33e-01 | Ascl1_targets_cluster7_features |
| Ascl1WT | A2N | 0 019209289 | 0 067873866 | 1339 | 0 732683455 | ns | ns 7.33e-01 | Ascl1_targets_cluster7_features |
| Ascl1WTBcl2 | O2N | -0 002254075 | 0 055101854 | 192 | 0 000185763 | **** | **** 1.86e-04 | Ascl1_targets_cluster7_features |
| Ascl1WTBcl2 | A2N | -0 021162533 | 0 080700699 | 1068 | 0 000185763 | **** | **** 1.86e-04 | Ascl1_targets_cluster7_features |
| Ascl1SA6 | O2N | -0 050896921 | 0 0800438 | 375 | 2 00E-09 | **** | **** 2.00e-09 | Ascl1_targets_cluster7_features |
| Ascl1SA6 | A2N | -0 02177464 | 0 07932025 | 1821 | 2 00E-09 | **** | **** 2.00e-09 | Ascl1_targets_cluster7_features |
| Ascl1SA6Bcl2 | O2N | -0 074932076 | 0 078785473 | 93 | 2 09E-05 | **** | **** 2.09e-05 | Ascl1_targets_cluster7_features |
| Ascl1SA6Bcl2 | A2N | -0 037024849 | 0 071405083 | 1423 | 2 09E-05 | **** | **** 2.09e-05 | Ascl1_targets_cluster7_features |

**Table 4.**

| Description | group | ID | GeneRatio | BgRatio | RichFactor | FoldEnrichment | zScore | pvalue | p.adjust | qvalue |
|---|---|---|---|---|---|---|---|---|---|---|
| gliogenesis | Control | GO:0042063 | 100/2648 | 412/28832 | 0 242718447 | 2 642771243 | 10 68032603 | 7 60E-20 | 1 56E-17 | 8 84E-18 |
| fatty acid metabolic process | Control | GO:0006631 | 128/2648 | 477/28832 | 0 268343816 | 2 921785834 | 13 45922951 | 1 79E-29 | 1 68E-26 | 9 49E-27 |
| Notch signaling pathway | Control | GO:0007219 | 44/2648 | 197/28832 | 0 223350254 | 2 431886147 | 6 413038603 | 2 32E-08 | 4 26E-07 | 2 41E-07 |
| glycolytic process | Control | GO:0006096 | 25/2648 | 113/28832 | 0 221238938 | 2 408897682 | 4 772048697 | 2 77E-05 | 0 000208337 | 0 00011786 |
| oxidative phosphorylation | Control | NA | NA | NA | NA | NA | NA | NA | 1 | NA |
| vesicle-mediated transport in synapse | Control | GO:0099003 | 63/2648 | 318/28832 | 0 198113208 | 2 157099698 | 6 598209714 | 4 11E-09 | 9 60E-08 | 5 43E-08 |
| regulation of synapse organization | Control | GO:0050807 | 94/2648 | 393/28832 | 0 239185751 | 2 604306481 | 10 18350544 | 2 78E-18 | 4 91E-16 | 2 78E-16 |
| regulation of axonogenesis | Control | GO:0050770 | 36/2648 | 175/28832 | 0 205714286 | 2 23986189 | 5 231760096 | 3 30E-06 | 3 37E-05 | 1 91E-05 |
| dendrite development | Control | GO:0016358 | 75/2648 | 332/28832 | 0 225903614 | 2 459687693 | 8 507019056 | 1 70E-13 | 1 25E-11 | 7 06E-12 |
| regulation of neurogenesis | Control | GO:0050767 | 96/2648 | 473/28832 | 0 202959831 | 2 209870787 | 8 4371341 | 8 49E-14 | 6 63E-12 | 3 75E-12 |
| gliogenesis | Ascl1WT | GO:0042063 | 86/2313 | 412/28832 | 0 208737864 | 2 601958537 | 9 672237239 | 1 44E-16 | 4 00E-14 | 2 42E-14 |
| fatty acid metabolic process | Ascl1WT | GO:0006631 | 90/2313 | 477/28832 | 0 188679245 | 2 351923908 | 8 7929724 | 1 78E-14 | 3 76E-12 | 2 28E-12 |
| Notch signaling pathway | Ascl1WT | GO:0007219 | 42/2313 | 197/28832 | 0 21319797 | 2 657554629 | 6 894320485 | 4 13E-09 | 2 18E-07 | 1 32E-07 |
| glycolytic process | Ascl1WT | NA | NA | NA | NA | NA | NA | NA | 1 | NA |
| oxidative phosphorylation | Ascl1WT | NA | NA | NA | NA | NA | NA | NA | 1 | NA |
| vesicle-mediated transport in synapse | Ascl1WT | GO:0099003 | 38/2313 | 318/28832 | 0 119496855 | 1 489551809 | 2 592520771 | 0 0089578 | 0 032408668 | 0 0196091 |
| regulation of synapse organization | Ascl1WT | GO:0050807 | 70/2313 | 393/28832 | 0 178117048 | 2 220264046 | 7 193363843 | 2 10E-10 | 1 65E-08 | 9 98E-09 |
| regulation of axonogenesis | Ascl1WT | GO:0050770 | 27/2313 | 175/28832 | 0 154285714 | 1 923201779 | 3 617757003 | 0 0007723 | 0 004993806 | 0 00302154 |
| dendrite development | Ascl1WT | GO:0016358 | 49/2313 | 332/28832 | 0 147590361 | 1 839742889 | 4 54498071 | 2 52E-05 | 0 00030034 | 0 00018172 |
| regulation of neurogenesis | Ascl1WT | GO:0050767 | 94/2313 | 473/28832 | 0 198731501 | 2 477227254 | 9 566906365 | 1 55E-16 | 4 14E-14 | 2 51E-14 |
| gliogenesis | Ascl1WTBcl2 | GO:0042063 | 45/1452 | 412/28832 | 0 109223301 | 2 168819706 | 5 502759545 | 1 02E-06 | 6 85E-05 | 5 30E-05 |
| fatty acid metabolic process | Ascl1WTBcl2 | NA | NA | NA | NA | NA | NA | NA | 1 | NA |
| Notch signaling pathway | Ascl1WTBcl2 | GO:0007219 | 23/1452 | 197/28832 | 0 116751269 | 2 318300681 | 4 275583921 | 0 0001588 | 0 003415458 | 0 00264187 |
| glycolytic process | Ascl1WTBcl2 | NA | NA | NA | NA | NA | NA | NA | 1 | NA |
| oxidative phosphorylation | Ascl1WTBcl2 | NA | NA | NA | NA | NA | NA | NA | 1 | NA |
| vesicle-mediated transport in synapse | Ascl1WTBcl2 | NA | NA | NA | NA | NA | NA | NA | 1 | NA |
| regulation of synapse organization | Ascl1WTBcl2 | GO:0050807 | 40/1452 | 393/28832 | 0 10178117 | 2 021043187 | 4 69331459 | 2 12E-05 | 0 000737207 | 0 00057023 |
| regulation of axonogenesis | Ascl1WTBcl2 | GO:0050770 | 21/1452 | 175/28832 | 0 12 | 2 382809917 | 4 225349013 | 0 0002055 | 0 00401603 | 0 00310641 |
| dendrite development | Ascl1WTBcl2 | GO:0016358 | 36/1452 | 332/28832 | 0 108433735 | 2 153141492 | 4 866599981 | 1 39E-05 | 0 000551204 | 0 00042636 |
| regulation of neurogenesis | Ascl1WTBcl2 | GO:0050767 | 49/1452 | 473/28832 | 0 10359408 | 2 057041683 | 5 337939744 | 1 60E-06 | 9 99E-05 | 7 73E-05 |
| gliogenesis | Ascl1SA6 | GO:0042063 | 38/958 | 412/28832 | 0 09223301 | 2 775847741 | 6 73062289 | 1 75E-08 | 1 19E-06 | 8 62E-07 |
| fatty acid metabolic process | Ascl1SA6 | NA | NA | NA | NA | NA | NA | NA | 1 | NA |
| Notch signaling pathway | Ascl1SA6 | GO:0007219 | 15/958 | 197/28832 | 0 076142132 | 2 291576147 | 3 372232387 | 0 0025367 | 0 018610843 | 0 0135385 |
| glycolytic process | Ascl1SA6 | NA | NA | NA | NA | NA | NA | NA | 1 | NA |
| oxidative phosphorylation | Ascl1SA6 | NA | NA | NA | NA | NA | NA | NA | 1 | NA |
| vesicle-mediated transport in synapse | Ascl1SA6 | GO:0099003 | 35/958 | 318/28832 | 0 110062893 | 3 312456507 | 7 687250267 | 6 64E-10 | 6 65E-08 | 4 84E-08 |
| regulation of synapse organization | Ascl1SA6 | GO:0050807 | 60/958 | 393/28832 | 0 152671756 | 4 59481426 | 13 30235489 | 4 46E-23 | 7 44E-20 | 5 42E-20 |
| regulation of axonogenesis | Ascl1SA6 | GO:0050770 | 42/958 | 175/28832 | 0 24 | 7 223048017 | 15 30804655 | 2 66E-24 | 6 67E-21 | 4 85E-21 |
| dendrite development | Ascl1SA6 | GO:0016358 | 49/958 | 332/28832 | 0 147590361 | 4 441884448 | 11 69380527 | 1 93E-18 | 9 68E-16 | 7 04E-16 |
| regulation of neurogenesis | Ascl1SA6 | GO:0050767 | 63/958 | 473/28832 | 0 133192389 | 4 008562589 | 12 23086365 | 5 65E-21 | 4 04E-18 | 2 94E-18 |
| gliogenesis | Ascl1SA6Bcl2 | GO:0042063 | 62/1839 | 412/28832 | 0 150485437 | 2 359323609 | 7 253616178 | 2 75E-10 | 1 20E-08 | 8 29E-09 |
| fatty acid metabolic process | Ascl1SA6Bcl2 | NA | NA | NA | NA | NA | NA | NA | 1 | NA |
| Notch signaling pathway | Ascl1SA6Bcl2 | GO:0007219 | 25/1839 | 197/28832 | 0 126903553 | 1 989604812 | 3 637822108 | 0 0008037 | 0 005609783 | 0 00388145 |
| glycolytic process | Ascl1SA6Bcl2 | NA | NA | NA | NA | NA | NA | NA | 1 | NA |
| oxidative phosphorylation | Ascl1SA6Bcl2 | GO:0006119 | 21/1839 | 154/28832 | 0 136363636 | 2 137920807 | 3 695669358 | 0 0008073 | 0 005625819 | 0 00389255 |
| vesicle-mediated transport in synapse | Ascl1SA6Bcl2 | GO:0099003 | 94/1839 | 318/28832 | 0 295597484 | 4 634402755 | 17 01032772 | 1 39E-37 | 4 00E-34 | 2 77E-34 |
| regulation of synapse organization | Ascl1SA6Bcl2 | GO:0050807 | 96/1839 | 393/28832 | 0 244274809 | 3 829761445 | 14 74293939 | 5 84E-31 | 6 33E-28 | 4 38E-28 |
| regulation of axonogenesis | Ascl1SA6Bcl2 | GO:0050770 | 54/1839 | 175/28832 | 0 308571429 | 4 837809368 | 13 29175114 | 4 07E-23 | 2 12E-20 | 1 47E-20 |
| dendrite development | Ascl1SA6Bcl2 | GO:0016358 | 98/1839 | 332/28832 | 0 295180723 | 4 627868734 | 17 35374921 | 4 42E-39 | 2 54E-35 | 1 76E-35 |
| regulation of neurogenesis | Ascl1SA6Bcl2 | GO:0050767 | 96/1839 | 473/28832 | 0 202959831 | 3 182021666 | 12 48933495 | 2 12E-24 | 1 35E-21 | 9 34E-22 |

| geneID | Count | log_p |
| --- | --- | --- |
| Daam2/Gli3/Atp1b2/Ptn/Plpp3/Kcnj10/Clu | 100 | 16 80628614 |
| Abhd3/Scd1/Acsbg1/Gstm1/Cyp4f15/Scd | 128 | 25 77551717 |
| Ttyh1/Tmem100/Notch3/Tgfb2/Timp4/Sox | 44 | 6 370635274 |
| Aldoc/Slc4a4/Eno1/Ppara/Ppargc1a/Pfkm | 25 | 3 681233507 |
| NA | 0 | 0 |
| Tspan7/Trim9/Nrxn1/Cspg5/Myo6/Pcdh17 | 63 | 7 017808478 |
| Dgkb/Sparcl1/Ptn/Slc7a11/Ncan/Nrxn1/Pt | 94 | 15 30918242 |
| Vegfa/Sema6d/Apoe/Ntrk3/Mt3/Lrp1/Dab | 36 | 4 472413205 |
| Prex2/Ptn/Ptprz1/Ngef/Apoe/Adgrb3/Myo6 | 75 | 10 9037327 |
| Daam2/Vegfa/Gli3/Ptn/Ptprz1/Gfap/Fgfr3/ | 96 | 11 1784035 |
| Olig1/Dbi/Egfr/Cd9/Ptprz1/Metrn/Plpp3/N | 86 | 13 39767543 |
| Dbi/Acot1/Acsbg1/Pdpn/Fads2/Scd2/Aca | 90 | 11 42466664 |
| Ttyh1/Tmem100/Lfng/Hes5/Sox2/Maml2/ | 42 | 6 660764958 |
| NA | 0 | 0 |
| NA | 0 | 0 |
| Cspg5/Pcdh17/Tspan7/Nlgn1/Unc13c/Syt | 38 | 1 489338822 |
| Apoe/Ntrk2/Sparcl1/Rhoa/Dgkb/Ptprt/Gric | 70 | 7 782525015 |
| Apoe/Metrn/Ntrk2/Lrrc4c/Draxin/Vim/Mt3/ | 27 | 2 301568341 |
| Apoe/Ptprz1/Epha5/Ntrk2/Nr3c1/Rhoa/Ptr | 49 | 3 522387008 |
| Apoe/Egfr/Ptprz1/Metrn/Ntrk2/Hes5/Serpi | 94 | 13 38276325 |
| Gstp1/Cdk6/Ascl1/Sox4/Lingo1/Dlx2/Myc | 45 | 4 164109806 |
| NA | 0 | 0 |
| Cdk6/Ascl1/Dlx2/Maml2/Zmiz1/Prag1/Dll | 23 | 2 466551014 |
| NA | 0 | 0 |
| NA | 0 | 0 |
| NA | 0 | 0 |
| Asic2/Lrrn1/Flrt2/Magi2/Mdga1/Nrp2/Prkc | 40 | 3 132410404 |
| Draxin/Zeb2/Sema5a/Map2/Pak3/Sipa1l1/ | 21 | 2 396203024 |
| Sez6/Map2/Pak3/Dynlt1b/Sipa1l1/Nsmf/S | 36 | 3 258687969 |
| Chd7/Btg2/Ascl1/Draxin/Lingo1/Dlx2/Myc | 49 | 4 000540913 |
| Sox11/Miat/Akt1/Nfix/Rtn4/B4galt5/Hdac1 | 38 | 5 926086615 |
| NA | 0 | 0 |
| Sorbs2/Cbfa2t2/Cntn6/Bmp7/Rbpj/Mfng/ | 15 | 1 730233954 |
| NA | 0 | 0 |
| NA | 0 | 0 |
| Syt6/Calm1/Arc/Cdk5r1/Akap5/Actb/Ap3b | 35 | 7 176985715 |
| Lrfn2/Ephb1/Dbn1/Flrt1/Akt1/Epha7/Lzts1 | 60 | 19 12820289 |
| Dbn1/Epha7/Rtn4/Brsk2/Gsk3b/Golga4/S | 42 | 20 17611786 |
| Ephb1/Dbn1/Flrt1/Lzts1/Arc/Gsk3b/Cdk5 | 49 | 15 01403881 |
| Sox11/Dbn1/Epha7/Rtn4/Gsk3b/Dpysl5/G | 63 | 17 3932599 |
| Nab2/Akt1/Sox11/Hdac11/Tiam1/Tuba1a/ | 62 | 7 921442946 |
| NA | 0 | 0 |
| Sorbs2/Bmp7/Gdpd5/Mfng/Neurl1a/Jag2/ | 25 | 2 251053953 |
| NA | 0 | 0 |
| Snca/Chchd2/Pde2a/Chchd10/mt-Nd4l/m | 21 | 2 24981427 |
| Snca/Calm2/Rapgef4/Calm1/Rph3al/Ap3b | 94 | 33 39820778 |
| Map1b/Dbn1/Snca/Akt1/Rapgef4/Dmpk/C | 96 | 27 19830248 |
| Map1b/Dbn1/Plxnd1/Golga4/Rufy3/Fstl4/ | 54 | 19 67272163 |
| Map1b/Dbn1/Dpysl5/Rapgef4/Rbfox2/Fstl | 98 | 34 59551012 |
| Map1b/Dbn1/Dpysl5/Plxnd1/Golga4/Sox1 | 96 | 20 86965823 |

Table 5.

a

| Ranked genes in cells moving towards alternative fate | | | | | |
|---|---|---|---|---|---|
| gene | score | logfoldchange | pval | pval_adj | group |
| Ptprz1 | 41 81870651 | 2 322840691 | 1 975E-181 | 2 9837E-178 | Away from Pseudotime End |
| Olig1 | 35 66394424 | 2 982382774 | 4 9784E-133 | 3 7605E-130 | Away from Pseudotime End |
| Pde4b | 34 13267517 | 2 674453974 | 1 5246E-133 | 1 1913E-130 | Away from Pseudotime End |
| Qk | 33 95996475 | 1 813656449 | 1 6371E-130 | 1 0599E-127 | Away from Pseudotime End |
| Erbb4 | 30 75954056 | 2 954482079 | 2 1815E-116 | 9 887E-114 | Away from Pseudotime End |
| Zfp36l1 | 29 32798576 | 2 983387709 | 9 732E-106 | 2 9802E-103 | Away from Pseudotime End |
| Dbi | 29 26375198 | 2 112361193 | 6 4854E-110 | 2 261E-107 | Away from Pseudotime End |
| Prss23 | 29 11261559 | 2 525969982 | 1 5503E-105 | 4 6226E-103 | Away from Pseudotime End |
| Myo10 | 28 58559227 | 2 058185816 | 1 295E-106 | 4 1923E-104 | Away from Pseudotime End |
| Maml2 | 28 53961182 | 2 062407255 | 1 8684E-106 | 5 8807E-104 | Away from Pseudotime End |
| Wscd1 | 27 71046257 | 1 851157069 | 1 2216E-101 | 3 3352E-99 | Away from Pseudotime End |
| Zcchc24 | 27 62107658 | 2 246401072 | 5 7493E-99 | 1 46387E-96 | Away from Pseudotime End |
| Sirt2 | 26 74124336 | 1 706843615 | 1 56457E-95 | 3 69319E-93 | Away from Pseudotime End |
| Luzp2 | 26 5007 | 3 016855478 | 5 45597E-92 | 1 22414E-89 | Away from Pseudotime End |
| Gm2a | 26 13518143 | 1 891812563 | 8 2185E-94 | 1 86239E-91 | Away from Pseudotime End |
| Exoc7 | 25 63935661 | 2 192194939 | 4 83469E-89 | 1 01444E-86 | Away from Pseudotime End |
| Asic2 | 25 35301208 | 2 646075964 | 2 81284E-89 | 5 95717E-87 | Away from Pseudotime End |
| Tmem132b | 24 24751091 | 2 328176975 | 1 9885E-83 | 3 69356E-81 | Away from Pseudotime End |
| Me3 | 23 94413948 | 1 714330673 | 3 72216E-84 | 7 14812E-82 | Away from Pseudotime End |
| Sema5a | 23 86820412 | 1 809063554 | 1 82034E-85 | 3 5561E-83 | Away from Pseudotime End |
| Epb41 | 23 76925468 | 1 756753445 | 6 29657E-81 | 1 1061E-78 | Away from Pseudotime End |
| Apoe | 23 74376488 | 2 422916889 | 5 83016E-83 | 1 05694E-80 | Away from Pseudotime End |
| Ednrb | 23 67308044 | 2 803675652 | 1 80325E-78 | 2 96112E-76 | Away from Pseudotime End |
| Tsc22d4 | 23 57584763 | 2 181234598 | 1 51656E-79 | 2 60354E-77 | Away from Pseudotime End |
| Olig2 | 21 7745533 | 1 484841347 | 4 70446E-73 | 6 74732E-71 | Away from Pseudotime End |
| Cadm2 | 21 41749763 | 1 407770514 | 4 77925E-72 | 6 64433E-70 | Away from Pseudotime End |
| Fabp7 | 21 09659576 | 2 099472046 | 4 25499E-69 | 5 50985E-67 | Away from Pseudotime End |
| Usp24 | 20 78937149 | 2 25931406 | 2 26878E-66 | 2 66387E-64 | Away from Pseudotime End |
| Lrch3 | 20 65449524 | 1 550593615 | 1 96684E-66 | 2 33353E-64 | Away from Pseudotime End |
| Cd9 | 20 47568512 | 1 33864069 | 8 47946E-68 | 1 06752E-65 | Away from Pseudotime End |
| Dock1 | 20 22120857 | 1 616512656 | 1 37297E-66 | 1 64619E-64 | Away from Pseudotime End |
| Plpp3 | 20 12818718 | 2 102138281 | 5 58661E-64 | 6 20579E-62 | Away from Pseudotime End |
| Xist | 19 95468903 | 2 166398525 | 4 72785E-65 | 5 38892E-63 | Away from Pseudotime End |
| Ntrk2 | 19 85322571 | 1 66197145 | 9 34399E-63 | 1 018E-60 | Away from Pseudotime End |
| Csmd2 | 19 78234482 | 2 174263954 | 9 55559E-62 | 1 0025E-59 | Away from Pseudotime End |
| Draxin | 19 76750946 | 2 800713778 | 3 86208E-61 | 3 97812E-59 | Away from Pseudotime End |
| Cst3 | 19 72037125 | 1 119672775 | 4 73126E-63 | 5 17947E-61 | Away from Pseudotime End |
| Slc24a3 | 19 43658447 | 2 353301287 | 3 82628E-60 | 3 73739E-58 | Away from Pseudotime End |
| Cacng4 | 19 41666794 | 1 690677166 | 1 29114E-60 | 1 29462E-58 | Away from Pseudotime End |
| Alcam | 19 39479065 | 1 412531018 | 4 05671E-62 | 4 31592E-60 | Away from Pseudotime End |
| Lrrn1 | 19 21965408 | 1 439261436 | 2 4848E-60 | 2 45887E-58 | Away from Pseudotime End |
| Xylt1 | 19 16338158 | 2 902636528 | 3 7328E-58 | 3 39715E-56 | Away from Pseudotime End |
| Mdga2 | 19 10343742 | 1 652858973 | 2 65969E-60 | 2 62049E-58 | Away from Pseudotime End |
| Gpm6b | 19 09468269 | 0 763910174 | 6 29456E-62 | 6 63446E-60 | Away from Pseudotime End |
| Nlgn1 | 18 79859924 | 1 356371284 | 3 41385E-60 | 3 34898E-58 | Away from Pseudotime End |
| Syn2 | 18 6969223 | 1 210040092 | 7 07763E-59 | 6 68276E-57 | Away from Pseudotime End |
| Cdh20 | 18 6762104 | 1 539964795 | 1 19156E-58 | 1 10212E-56 | Away from Pseudotime End |
| Sox6 | 18 49116516 | 1 48011744 | 7 62719E-57 | 6 75155E-55 | Away from Pseudotime End |
| Gpr37l1 | 18 4736557 | 1 307541132 | 3 22612E-57 | 2 87823E-55 | Away from Pseudotime End |
| Csmd1 | 18 00937843 | 2 166219234 | 4 74438E-54 | 3 82606E-52 | Away from Pseudotime End |
| Mafa | 47 06087112 | 5 80420208 | 0 | 0 | Towards Pseudotime End |
| Pkib | 45 48893738 | 3 794736147 | 3 3039E-240 | 1 0696E-236 | Towards Pseudotime End |
| Igfbpl1 | 44 71337891 | 4 467459679 | 1 259E-272 | 4 755E-269 | Towards Pseudotime End |
| Gm48893 | 42 08690643 | 4 732989311 | 2 6782E-279 | 1 5173E-275 | Towards Pseudotime End |
| Ttc28 | 41 91828156 | 2 865819216 | 1 1561E-199 | 2 1833E-196 | Towards Pseudotime End |
| Lhx3 | 40 36789322 | 5 90758276 | 1 483E-287 | 1 6803E-283 | Towards Pseudotime End |
| Slit3 | 40 36071396 | 5 557960987 | 4 5859E-285 | 3 464E-281 | Towards Pseudotime End |
| Map1b | 39 50159454 | 3 682989597 | 4 1562E-210 | 1 0465E-206 | Towards Pseudotime End |
| Wnk2 | 39 259655 | 3 065334082 | 4 6512E-200 | 9 5819E-197 | Towards Pseudotime End |
| Cib3 | 38 82245636 | 6 086823463 | 3 505E-274 | 1 5885E-270 | Towards Pseudotime End |
| Arhgap18 | 37 37925339 | 3 94168973 | 4 1098E-200 | 9 3132E-197 | Towards Pseudotime End |
| Nt5dc2 | 37 11937714 | 3 160920382 | 3 7772E-169 | 5 3496E-166 | Towards Pseudotime End |
| Ccdc88b | 35 77938461 | 5 355216026 | 8 9271E-238 | 2 5287E-234 | Towards Pseudotime End |
| Sel1l3 | 34 85439682 | 3 20741725 | 9 3703E-155 | 1 1797E-151 | Towards Pseudotime End |
| Prdx5 | 33 73621368 | 2 364691496 | 9 4492E-145 | 8 922E-142 | Towards Pseudotime End |
| Thsd7b | 32 89041901 | 2 917711973 | 3 8535E-137 | 3 3586E-134 | Towards Pseudotime End |
| Dpysl5 | 32 77949142 | 2 802078724 | 4 3476E-151 | 4 9261E-148 | Towards Pseudotime End |
| Rapgef4 | 32 77040863 | 4 290394306 | 1 3659E-196 | 2 381E-193 | Towards Pseudotime End |
| Srrm4 | 32 01224136 | 2 991588593 | 1 9041E-129 | 1 1662E-126 | Towards Pseudotime End |
| Kcnh3 | 31 92088699 | 5 428154945 | 2 9801E-195 | 4 8237E-192 | Towards Pseudotime End |
| Trpc4ap | 31 6861515 | 1 952864885 | 2 091E-132 | 1 4807E-129 | Towards Pseudotime End |
| Cbfa2t3 | 31 3387661 | 3 073669195 | 1 2615E-145 | 1 2994E-142 | Towards Pseudotime End |
| Olfm1 | 30 79904747 | 1 838173747 | 2 0291E-119 | 1 0218E-116 | Towards Pseudotime End |
| Gdpd5 | 30 72489357 | 3 221343994 | 5 7555E-143 | 5 2171E-140 | Towards Pseudotime End |
| Myl12b | 30 49261284 | 1 660172582 | 2 569E-118 | 1 2387E-115 | Towards Pseudotime End |
| Rbm38 | 30 21897125 | 2 377360344 | 1 3366E-129 | 8 4135E-127 | Towards Pseudotime End |
| Kif5c | 30 16583061 | 2 298593283 | 4 9579E-126 | 2 8808E-123 | Towards Pseudotime End |
| Trnp1 | 30 08090019 | 3 249686956 | 1 2547E-155 | 1 6725E-152 | Towards Pseudotime End |
| Cacna1c | 29 85645676 | 3 027718306 | 4 9855E-145 | 4 912E-142 | Towards Pseudotime End |
| Cotl1 | 29 49959373 | 1 895463705 | 1 6701E-109 | 5 7342E-107 | Towards Pseudotime End |
| Shf | 28 97579002 | 2 924389362 | 5 2604E-133 | 3 8454E-130 | Towards Pseudotime End |
| Pak7 | 28 95966148 | 2 486794949 | 1 2668E-113 | 5 4166E-111 | Towards Pseudotime End |

| | | | | | |
|---|---|---|---|---|---|
| Atp2b4 | 28 95912933 | 2 794488192 | 1 94E-135 | 1 5701E-132 | Towards Pseudotime End |
| Eepd1 | 28 93954659 | 2 399673462 | 1 0219E-112 | 4 2105E-110 | Towards Pseudotime End |
| Sox11 | 28 80961609 | 1 535788894 | 8 8718E-107 | 2 9137E-104 | Towards Pseudotime End |
| Clstn2 | 28 77786636 | 3 370522738 | 5 6176E-131 | 3 7442E-128 | Towards Pseudotime End |
| Basp1 | 28 75138664 | 2 936972857 | 1 0343E-123 | 5 7166E-121 | Towards Pseudotime End |
| Akt1 | 28 70598984 | 1 817057133 | 4 9087E-112 | 1 8853E-109 | Towards Pseudotime End |
| Tubb2a | 28 64892197 | 2 095999002 | 2 5849E-123 | 1 3947E-120 | Towards Pseudotime End |
| Rassf4 | 28 44004822 | 2 252565145 | 2 3839E-103 | 6 8827E-101 | Towards Pseudotime End |
| Mir124a-1hg | 28 34680748 | 4 047777653 | 3 9833E-148 | 4 2984E-145 | Towards Pseudotime End |
| Efhd2 | 28 17425728 | 2 541722059 | 2 4887E-111 | 8 9518E-109 | Towards Pseudotime End |
| Sult2b1 | 28 04395103 | 4 452260971 | 1 5061E-151 | 1 7963E-148 | Towards Pseudotime End |
| Cacna2d2 | 27 55907822 | 2 700126886 | 5 7631E-119 | 2 8391E-116 | Towards Pseudotime End |
| Vasp | 27 38487053 | 2 192193747 | 2 264E-114 | 1 006E-111 | Towards Pseudotime End |
| Plxnd1 | 27 22809792 | 3 274873734 | 2 6574E-131 | 1 8248E-128 | Towards Pseudotime End |
| Tecta | 27 08562851 | 2 658173084 | 2 9168E-112 | 1 1596E-109 | Towards Pseudotime End |
| Rbfox3 | 27 06219482 | 2 594800711 | 3 2272E-112 | 1 2609E-109 | Towards Pseudotime End |
| Nppc | 26 96769142 | 4 223303318 | 1 5945E-136 | 1 3382E-133 | Towards Pseudotime End |
| Hpcal1 | 26 81870079 | 2 21947217 | 7 6704E-104 | 2 2574E-101 | Towards Pseudotime End |

Percentage of cells heading towards different fates, RNA Velocity analysis

**b**

| **RV** | **Towards Astrocyt** | **Towards Neurona** | **Towards Alternative Fate (%)** |
|---|---|---|---|
| Ascl1WT | 70 9 | 14 2 | 14 9 |
| Ascl1WTBcl2 | 22 6 | 57 4 | 20 |
| Ascl1SA6 | 12 4 | 85 | 2 6 |
| Ascl1SA6Bcl2 | 1 5 | 97 9 | 0 7 |

Table 6.

Pvalb RNAscope dot counts / cell

| Ascl1WT | Ascl1WT+Bcl2 | Ascl1SA6 | Ascl1SA6+Bcl2 |
|---|---|---|---|
| 2 | 2 | 3 | 3 |
| 0 | 16 | 6 | 4 |
| 3 | 2 | 6 | 9 |
| 0 | 1 | 8 | 8 |
| 1 | 1 | 6 | 1 |
| 3 | 13 | 5 | 1 |
| 1 | 0 | 12 | 8 |
| 1 | 6 | 15 | 21 |
| 1 | 1 | 6 | 13 |
| 2 | 2 | 1 | 6 |
| 3 | 2 | 28 | 12 |
| 2 | 5 | 10 | 6 |
| 2 | 3 | 6 | 1 |
| 2 | 1 | 20 | 4 |
| 2 | 1 | 1 | 2 |
| 1 | 1 | 6 | 11 |
| 1 | 3 | 1 | 48 |
| 1 | 3 | 10 | 9 |
| 5 | | 13 | 3 |
| 1 | | 2 | 3 |
| 1 | | 11 | 3 |
| | | 2 | 2 |
| | | 26 | 6 |
| | | | 6 |
| | | | 6 |
| | | | 21 |
| | | | 27 |
| | | | 5 |
| | | | 7 |
| | | | 3 |
| | | | 8 |
| | | | 9 |
| | | | 4 |
| | | | 2 |
| | | | 20 |
| | | | 1 |
| | | | 8 |
| | | | 9 |
| | | | 5 |
| | | | 10 |

| Stats | Ascl1WT | Ascl1WT+Bcl2 | Ascl1SA6 | Ascl1SA6+Bcl2 |
|---|---|---|---|---|
| Number of values | 21 | 18 | 23 | 40 |
| Minimum | 0 | 0 | 1 | 1 |
| Maximum | 5 | 16 | 28 | 48 |
| Range | 5 | 16 | 27 | 47 |
| Mean | 1 667 | 3 5 | 8 87 | 8 375 |
| Std. Deviation | 1 155 | 4 301 | 7 491 | 8 787 |
| Std. Error of Mean | 0 252 | 1 014 | 1 562 | 1 389 |
| Sum | 35 | 63 | 204 | 335 |

| | |
|---|---|
| Number of families | 1 |
| Number of comparisons per fami | 6 |
| Alpha | 0 05 |

| Tukey's multiple comparisons te | Mean Diff. | 95.00% CI of diff. | Below threshold | Summary | Adjusted P Value | |
|---|---|---|---|---|---|---|
| Ascl1WT vs. Ascl1WT+Bcl2 | -1 833 | -7.577 to 3.910 | No | ns | 0 838 | A-B |
| Ascl1WT vs. Ascl1SA6 | -7 203 | -12.60 to -1.806 | Yes | ** | 0 004 | A-C |
| Ascl1WT vs. Ascl1SA6+Bcl2 | -6 708 | -11.53 to -1.890 | Yes | ** | 0 0024 | A-D |
| Ascl1WT+Bcl2 vs. Ascl1SA6 | -5 37 | -11.00 to 0.2577 | No | ns | 0 0671 | B-C |
| Ascl1WT+Bcl2 vs. Ascl1SA6+B | -4 875 | -9.950 to 0.2002 | No | ns | 0 0645 | B-D |
| Ascl1SA6 vs. Ascl1SA6+Bcl2 | 0 4946 | -4.185 to 5.174 | No | ns | 0 9926 | C-D |

| Test details | Mean 1 | Mean 2 | Mean Diff. | SE of diff. | n1 | n2 | q | DF |
|---|---|---|---|---|---|---|---|---|
| Ascl1WT vs. Ascl1WT+Bcl2 | 1 667 | 3 5 | -1 833 | 2 198 | 21 | 18 | 1 18 | 98 |
| Ascl1WT vs. Ascl1SA6 | 1 667 | 8 87 | -7 203 | 2 065 | 21 | 23 | 4 933 | 98 |
| Ascl1WT vs. Ascl1SA6+Bcl2 | 1 667 | 8 375 | -6 708 | 1 844 | 21 | 40 | 5 146 | 98 |
| Ascl1WT+Bcl2 vs. Ascl1SA6 | 3 5 | 8 87 | -5 37 | 2 153 | 18 | 23 | 3 527 | 98 |
| Ascl1WT+Bcl2 vs. Ascl1SA6+B | 3 5 | 8 375 | -4 875 | 1 942 | 18 | 40 | 3 55 | 98 |
| Ascl1SA6 vs. Ascl1SA6+Bcl2 | 8 87 | 8 375 | 0 4946 | 1 79 | 23 | 40 | 0 3907 | 98 |

Table 7.

Dlx2 RNAscope dot counts / cell

| Ascl1WT | Ascl1WT+Bcl2 | Ascl1SA6 | Ascl1SA6+Bcl2 |
|---|---|---|---|
| 0 | 34 | 7 | 1 |
| 0 | 23 | 3 | 1 |
| 3 | 7 | 0 | 1 |
| 3 | 6 | 14 | 0 |
| 1 | 0 | 0 | 4 |
| 2 | 9 | 2 | 0 |
| 0 | 12 | 8 | 6 |
| 0 | 0 | 9 | 9 |
| 1 | 14 | 1 | 4 |
| 1 | 0 | 10 | 0 |
| 0 | 2 | 1 | 9 |
| 0 | 16 | 2 | 6 |
| 1 | 9 | 2 | 1 |
| 2 | 0 | 0 | 1 |
| 0 | 0 | 0 | 2 |
| 0 | 0 | 5 | 1 |
| 4 | 2 | 5 | 2 |
| 2 | 11 | 5 | 3 |
| 1 | | 4 | 5 |
| 0 | | 2 | 4 |
| | | 1 | 6 |
| | | 0 | 1 |
| | | 1 | 2 |
| | | 1 | 2 |
| | | 0 | 3 |
| | | | 4 |
| | | | 3 |
| | | | 5 |
| | | | 1 |
| | | | 3 |
| | | | 1 |
| | | | 0 |
| | | | 2 |
| | | | 2 |
| | | | 1 |
| | | | 1 |
| | | | 0 |
| | | | 3 |
| | | | 1 |

| | Ascl1WT | Ascl1WT+Bcl2 | Ascl1SA6 | Ascl1SA6+Bcl2 |
|---|---|---|---|---|
| Number of values | 20 | 18 | 25 | 39 |
| Minimum | 0 | 0 | 0 | 0 |
| Maximum | 4 | 34 | 14 | 9 |
| Range | 4 | 34 | 14 | 9 |
| Mean | 1 05 | 8 056 | 3 32 | 2 59 |
| Std. Deviation | 1 234 | 9 359 | 3 739 | 2 314 |
| Std. Error of Mean | 0 276 | 2 206 | 0 7477 | 0 3705 |
| Sum | 21 | 145 | 83 | 101 |

| | |
|---|---|
| Number of families | 1 |
| Number of comparisons per family | 6 |
| Alpha | 0 05 |

| Tukey's multiple comparisons test | Mean Diff. | 95.00% CI of diff. | Below thresho | Summary | Adjusted P Value | |
|---|---|---|---|---|---|---|
| Ascl1WT vs. Ascl1WT+Bcl2 | -7 006 | -10.90 to -3.115 | Yes | **** | <0.0001 | A-B |
| Ascl1WT vs. Ascl1SA6 | -2 27 | -5.862 to 1.322 | No | ns | 0 3549 | A-C |
| Ascl1WT vs. Ascl1SA6+Bcl2 | -1 54 | -4.833 to 1.753 | No | ns | 0 6144 | A-D |
| Ascl1WT+Bcl2 vs. Ascl1SA6 | 4 736 | 1.034 to 8.437 | Yes | ** | 0 0063 | B-C |
| Ascl1WT+Bcl2 vs. Ascl1SA6+Bcl2 | 5 466 | 2.054 to 8.878 | Yes | *** | 0 0004 | B-D |
| Ascl1SA6 vs. Ascl1SA6+Bcl2 | 0 7303 | -2.337 to 3.798 | No | ns | 0 9247 | C-D |

| Test details | Mean 1 | Mean 2 | Mean Diff. | SE of diff. | n1 | n2 | q | DF |
|---|---|---|---|---|---|---|---|---|
| Ascl1WT vs. Ascl1WT+Bcl2 | 1 05 | 8 056 | -7 006 | 1 488 | 20 | 18 | 6 656 | 98 |
| Ascl1WT vs. Ascl1SA6 | 1 05 | 3 32 | -2 27 | 1 374 | 20 | 25 | 2 336 | 98 |
| Ascl1WT vs. Ascl1SA6+Bcl2 | 1 05 | 2 59 | -1 54 | 1 26 | 20 | 39 | 1 728 | 98 |
| Ascl1WT+Bcl2 vs. Ascl1SA6 | 8 056 | 3 32 | 4 736 | 1 416 | 18 | 25 | 4 729 | 98 |
| Ascl1WT+Bcl2 vs. Ascl1SA6+Bcl2 | 8 056 | 2 59 | 5 466 | 1 305 | 18 | 39 | 5 921 | 98 |
| Ascl1SA6 vs. Ascl1SA6+Bcl2 | 3 32 | 2 59 | 0 7303 | 1 174 | 25 | 39 | 0 88 | 98 |

Table 8. MGE versus CGE Transcription factors expressed in induced Neurons

| TFs | p_val | avg_log2FC | pct.1 | pct.2 | p_val_adj | up_in | |
|---|---|---|---|---|---|---|---|
| Nfib | 3 65E-242 | -4 306874704 | 0 208 | 0 935 | 9 63E-238 | Up_in_CGE | CGE derived cells Lamp5, Vip |
| Zbtb16 | 1 91E-140 | -2 693284181 | 0 379 | 0 814 | 5 03E-136 | Up_in_CGE | |
| Prox1 | 1 32E-120 | -5 673897651 | 0 024 | 0 528 | 3 49E-116 | Up_in_CGE | |
| Zfp536 | 1 88E-116 | -1 468713173 | 0 772 | 0 969 | 4 96E-112 | Up_in_CGE | |
| Nr3c2 | 1 61E-104 | -2 038250601 | 0 549 | 0 842 | 4 24E-100 | Up_in_CGE | |
| Nfix | 1 05E-79 | -3 74965403 | 0 062 | 0 448 | 2 78E-75 | Up_in_CGE | |
| Tcf4 | 8 51E-65 | -0 452216363 | 1 | 1 | 2 25E-60 | Up_in_CGE | |
| Nfia | 4 51E-57 | -3 83254793 | 0 053 | 0 355 | 1 19E-52 | Up_in_CGE | |
| Nr2f2 | 1 14E-50 | -2 587902971 | 0 095 | 0 398 | 3 02E-46 | Up_in_CGE | |
| Pbx3 | 2 90E-33 | -3 776227551 | 0 087 | 0 3 | 7 65E-29 | Up_in_CGE | |
| Tcf12 | 3 28E-28 | -0 842921168 | 0 73 | 0 79 | 8 65E-24 | Up_in_CGE | |
| Myt1l | 6 07E-25 | -0 571765931 | 0 976 | 0 957 | 1 60E-20 | Up_in_CGE | |
| Klf8 | 6 66E-25 | -1 75280511 | 0 131 | 0 329 | 1 76E-20 | Up_in_CGE | |
| Id2 | 1 81E-24 | -2 987606493 | 0 036 | 0 19 | 4 77E-20 | Up_in_CGE | |
| Foxp1 | 1 62E-23 | -0 915650913 | 0 59 | 0 685 | 4 27E-19 | Up_in_CGE | |
| Nr2e1 | 2 25E-23 | -4 845578539 | 0 003 | 0 119 | 5 93E-19 | Up_in_CGE | |
| Zeb1 | 1 61E-22 | -0 975513697 | 0 54 | 0 66 | 4 24E-18 | Up_in_CGE | |
| Zfp57 | 2 02E-19 | -1 729665679 | 0 1 | 0 26 | 5 32E-15 | Up_in_CGE | |
| Nr2f1 | 3 82E-16 | -1 15229298 | 0 247 | 0 402 | 1 01E-11 | Up_in_CGE | |
| Aff2 | 8 21E-15 | -0 622281647 | 0 705 | 0 787 | 2 17E-10 | Up_in_CGE | |
| Bach2 | 8 71E-15 | -0 522945364 | 0 781 | 0 793 | 2 30E-10 | Up_in_CGE | |
| Foxn3 | 1 90E-14 | -0 467436723 | 0 778 | 0 826 | 5 01E-10 | Up_in_CGE | |
| Rreb1 | 2 12E-14 | -2 410137662 | 0 03 | 0 129 | 5 60E-10 | Up_in_CGE | |
| Lcorl | 8 25E-14 | -0 832353657 | 0 449 | 0 557 | 2 18E-09 | Up_in_CGE | |
| Tshz1 | 1 15E-13 | -1 877066715 | 0 092 | 0 212 | 3 03E-09 | Up_in_CGE | |
| Arid1b | 2 05E-12 | -0 564332446 | 0 714 | 0 741 | 5 41E-08 | Up_in_CGE | |
| Zkscan16 | 4 48E-12 | -1 379775671 | 0 133 | 0 252 | 1 18E-07 | Up_in_CGE | |
| Smad3 | 5 48E-12 | -2 666897502 | 0 031 | 0 116 | 1 45E-07 | Up_in_CGE | |
| Zfp40 | 1 18E-11 | -1 399841328 | 0 121 | 0 237 | 3 12E-07 | Up_in_CGE | |
| Zbtb20 | 4 78E-11 | -0 912539721 | 0 559 | 0 664 | 1 26E-06 | Up_in_CGE | |
| Cers5 | 2 83E-10 | -0 727999838 | 0 48 | 0 542 | 7 48E-06 | Up_in_CGE | |
| Nr4a1 | 2 12E-09 | -1 182700623 | 0 156 | 0 259 | 5 59E-05 | Up_in_CGE | |
| Egr1 | 2 57E-09 | -2 383577933 | 0 038 | 0 112 | 6 79E-05 | Up_in_CGE | |
| Pou2f1 | 3 56E-09 | -0 714301267 | 0 462 | 0 515 | 9 40E-05 | Up_in_CGE | |
| Zmiz1 | 6 30E-08 | -1 027076347 | 0 135 | 0 227 | 0 00166212 | Up_in_CGE | |
| Tead1 | 8 05E-08 | -1 288196937 | 0 116 | 0 202 | 0 0021252 | Up_in_CGE | |
| Dlx1 | 1 06E-07 | -1 091491846 | 0 236 | 0 321 | 0 00279227 | Up_in_CGE | |
| Zhx2 | 1 28E-07 | -1 087172551 | 0 122 | 0 212 | 0 00338295 | Up_in_CGE | |
| Zeb2 | 2 04E-07 | -0 329758147 | 0 955 | 0 966 | 0 00538901 | Up_in_CGE | |
| Rora | 2 83E-07 | -0 338915553 | 0 898 | 0 93 | 0 00748067 | Up_in_CGE | |
| Srebf2 | 5 05E-07 | -0 897440402 | 0 25 | 0 33 | 0 01334106 | Up_in_CGE | |
| Klf7 | 1 07E-06 | -0 996517134 | 0 243 | 0 319 | 0 02817066 | Up_in_CGE | |
| Dnajc1 | 1 39E-06 | -0 408808458 | 0 679 | 0 693 | 0 03667845 | Up_in_CGE | |
| Zfp46 | 1 85E-06 | -1 490541628 | 0 063 | 0 128 | 0 04870651 | Up_in_CGE | |
| L3mbtl1 | 1 34E-05 | -1 304801117 | 0 083 | 0 144 | 0 3525515 | Up_in_CGE | |
| Klf13 | 6 32E-05 | -0 944313427 | 0 142 | 0 206 | 1 | Up_in_CGE | |
| Npas2 | 6 72E-05 | -0 661046229 | 0 3 | 0 36 | 1 | Up_in_CGE | |
| Ar | 6 90E-05 | -0 82217665 | 0 138 | 0 206 | 1 | Up_in_CGE | |
| Dlx2 | 7 31E-05 | -1 102114392 | 0 098 | 0 156 | 1 | Up_in_CGE | |
| Mxd4 | 9 15E-05 | -1 061271495 | 0 092 | 0 149 | 1 | Up_in_CGE | |
| Nr2c2 | 0 000129487 | -0 388728882 | 0 538 | 0 548 | 1 | Up_in_CGE | |
| Arid5b | 0 000333835 | -0 618223838 | 0 405 | 0 436 | 1 | Up_in_CGE | |
| Hivep3 | 0 001025204 | -0 349787728 | 0 803 | 0 728 | 1 | Up_in_CGE | |
| Zfp407 | 0 002045497 | -0 430857695 | 0 431 | 0 455 | 1 | Up_in_CGE | |
| Tef | 0 002559595 | -0 483539727 | 0 358 | 0 394 | 1 | Up_in_CGE | |
| Zfp618 | 0 003063715 | -0 686139993 | 0 243 | 0 283 | 1 | Up_in_CGE | |
| Pou2f2 | 0 004273463 | -0 637250564 | 0 167 | 0 216 | 1 | Up_in_CGE | |
| Pias1 | 0 00449948 | -0 360208435 | 0 526 | 0 528 | 1 | Up_in_CGE | |
| Zfp944 | 0 005658692 | -0 90373098 | 0 077 | 0 114 | 1 | Up_in_CGE | |
| Thra | 0 008667688 | -0 463233989 | 0 44 | 0 443 | 1 | Up_in_CGE | |
| Jarid2 | 0 009671823 | -0 44125053 | 0 47 | 0 465 | 1 | Up_in_CGE | |
| A430033K04Rik | 0 026726843 | -0 685436046 | 0 164 | 0 193 | 1 | Up_in_CGE | |
| Bbx | 0 028853447 | -0 553472483 | 0 28 | 0 303 | 1 | Up_in_CGE | |
| Junb | 0 030484989 | -0 978427199 | 0 193 | 0 219 | 1 | Up_in_CGE | |
| Pou3f3 | 0 032204523 | -0 545033344 | 0 169 | 0 202 | 1 | Up_in_CGE | |
| Zhx3 | 0 032623788 | -0 465445825 | 0 469 | 0 452 | 1 | Up_in_CGE | |
| Foxg1 | 0 038418082 | -0 624639686 | 0 236 | 0 252 | 1 | Up_in_CGE | |
| Pknox2 | 0 041731136 | -0 305167865 | 0 666 | 0 638 | 1 | Up_in_CGE | |
| Zfp362 | 0 055162177 | -0 437276533 | 0 174 | 0 203 | 1 | Up_in_CGE | |
| Mier3 | 0 060585959 | -0 491189018 | 0 098 | 0 124 | 1 | Up_in_CGE | |
| Sox2 | 0 067225662 | -0 469051552 | 0 207 | 0 23 | 1 | Up_in_CGE | |
| Zfp560 | 0 076666354 | -0 480226323 | 0 327 | 0 327 | 1 | Up_in_CGE | |
| Dmtf1 | 0 076872799 | -0 470414992 | 0 294 | 0 305 | 1 | Up_in_CGE | |
| Arid2 | 0 083225851 | -0 411584871 | 0 336 | 0 345 | 1 | Up_in_CGE | |
| Ikzf4 | 0 084434858 | -0 388245811 | 0 23 | 0 253 | 1 | Up_in_CGE | |
| Zfp141 | 0 085418425 | -0 449022483 | 0 264 | 0 285 | 1 | Up_in_CGE | |
| Gtf2i | 0 08963951 | -0 343544018 | 0 528 | 0 51 | 1 | Up_in_CGE | |
| Hmgb1 | 0 098966451 | -0 381370393 | 0 349 | 0 352 | 1 | Up_in_CGE | |
| Mlxip | 0 101132826 | -0 427115778 | 0 224 | 0 245 | 1 | Up_in_CGE | |
| Deaf1 | 0 112255729 | -0 506335633 | 0 165 | 0 186 | 1 | Up_in_CGE | |
| Hmbox1 | 0 120552675 | -0 281330619 | 0 512 | 0 487 | 1 | Up_in_CGE | |
| Terf1 | 0 129814981 | -0 321350002 | 0 41 | 0 403 | 1 | Up_in_CGE | |
| Sp4 | 0 152616903 | -0 371699629 | 0 313 | 0 317 | 1 | Up_in_CGE | |

| | | | | | | | |
|---|---|---|---|---|---|---|---|
| 2610021A01Rik | 0 163017004 | -0 482126272 | 0 197 | 0 209 | 1 | Up_in_CGE | |
| Atf7 | 0 193097247 | -0 306866191 | 0 472 | 0 452 | 1 | Up_in_CGE | |
| Sp3 | 0 22095458 | -0 471917094 | 0 224 | 0 234 | 1 | Up_in_CGE | |
| Zfp68 | 0 236338527 | -0 423009056 | 0 173 | 0 186 | 1 | Up_in_CGE | |
| Sox11 | 0 250109243 | -0 600874295 | 0 114 | 0 127 | 1 | Up_in_CGE | |
| Ubtf | 0 281310671 | -0 500528613 | 0 116 | 0 129 | 1 | Up_in_CGE | |
| Zfp710 | 0 283964019 | -0 365239133 | 0 131 | 0 145 | 1 | Up_in_CGE | |
| Bcl11b | 0 284592422 | -0 326273928 | 0 533 | 0 414 | 1 | Up_in_CGE | |
| Zfhx2 | 0 293230585 | -0 403912441 | 0 343 | 0 328 | 1 | Up_in_CGE | |
| Zfp763 | 0 299606605 | -0 472483849 | 0 143 | 0 153 | 1 | Up_in_CGE | |
| Zfp566 | 0 306658646 | -0 301211587 | 0 134 | 0 112 | 1 | Up_in_CGE | |
| Arntl | 0 31646756 | -0 484831695 | 0 298 | 0 291 | 1 | Up_in_CGE | |
| Zfp53 | 0 343790344 | -0 352084171 | 0 166 | 0 177 | 1 | Up_in_CGE | |
| Snapc4 | 0 344453248 | -0 327531214 | 0 127 | 0 138 | 1 | Up_in_CGE | |
| Csdc2 | 0 346372533 | -0 467967013 | 0 233 | 0 231 | 1 | Up_in_CGE | |
| Pias4 | 0 364429136 | -0 590659574 | 0 11 | 0 119 | 1 | Up_in_CGE | |
| Ttf1 | 0 402868814 | -0 376159531 | 0 21 | 0 212 | 1 | Up_in_CGE | |
| Zscan21 | 0 403912479 | -0 434723149 | 0 166 | 0 173 | 1 | Up_in_CGE | |
| Zfp386 | 0 421661891 | -0 519585137 | 0 128 | 0 135 | 1 | Up_in_CGE | |
| Zfp651 | 0 451993778 | -0 306113373 | 0 16 | 0 141 | 1 | Up_in_CGE | |
| Zbtb25 | 0 455188358 | -0 35054041 | 0 143 | 0 15 | 1 | Up_in_CGE | |
| E2f5 | 0 488401338 | -0 358886203 | 0 103 | 0 112 | 1 | Up_in_CGE | |
| Stat3 | 0 499526031 | -0 266253277 | 0 198 | 0 173 | 1 | Up_in_CGE | |
| Hbp1 | 0 538426617 | -0 295781954 | 0 162 | 0 142 | 1 | Up_in_CGE | |
| Zfp52 | 0 576741726 | -0 36337896 | 0 102 | 0 108 | 1 | Up_in_CGE | |
| Tub | 0 587629658 | -0 305947455 | 0 23 | 0 227 | 1 | Up_in_CGE | |
| Atf4 | 0 589436173 | -0 394657499 | 0 152 | 0 153 | 1 | Up_in_CGE | |
| Zfp458 | 0 630208221 | -0 338262173 | 0 131 | 0 134 | 1 | Up_in_CGE | |
| Stat2 | 0 64260976 | -0 441788878 | 0 197 | 0 192 | 1 | Up_in_CGE | |
| Prdm5 | 0 654652555 | -0 356639234 | 0 113 | 0 102 | 1 | Up_in_CGE | |
| Sub1 | 0 674653237 | -0 391029832 | 0 185 | 0 181 | 1 | Up_in_CGE | |
| Zfp507 | 0 682370547 | -0 277477289 | 0 26 | 0 248 | 1 | Up_in_CGE | |
| Zfp605 | 0 717666827 | -0 378305531 | 0 109 | 0 112 | 1 | Up_in_CGE | |
| Vezf1 | 0 726295991 | -0 310890449 | 0 271 | 0 257 | 1 | Up_in_CGE | |
| Zfp653 | 0 743483626 | -0 268583729 | 0 112 | 0 103 | 1 | Up_in_CGE | |
| Zfp711 | 0 752910524 | -0 268735863 | 0 24 | 0 233 | 1 | Up_in_CGE | |
| Foxo3 | 0 767899294 | -0 359529742 | 0 219 | 0 195 | 1 | Up_in_CGE | |
| Zfp867 | 0 771186468 | -0 316058852 | 0 134 | 0 123 | 1 | Up_in_CGE | |
| Zfp341 | 0 78847352 | -0 362257502 | 0 171 | 0 158 | 1 | Up_in_CGE | |
| Klf9 | 0 830791877 | -0 374932803 | 0 217 | 0 208 | 1 | Up_in_CGE | |
| Zbtb24 | 0 864440114 | -0 414434041 | 0 143 | 0 138 | 1 | Up_in_CGE | |
| Hivep1 | 0 870010088 | -0 366195301 | 0 28 | 0 26 | 1 | Up_in_CGE | |
| 2700081O15Rik | 0 917596329 | -0 38999958 | 0 138 | 0 134 | 1 | Up_in_CGE | |
| Zfp707 | 0 921338044 | -0 327464477 | 0 122 | 0 117 | 1 | Up_in_CGE | |
| Zfp516 | 0 921644246 | -0 327442925 | 0 137 | 0 128 | 1 | Up_in_CGE | |
| Foxj2 | 0 949712313 | -0 334659141 | 0 121 | 0 117 | 1 | Up_in_CGE | |
| Nfe2l3 | 0 992484774 | -0 566150545 | 0 102 | 0 098 | 1 | Up_in_CGE | |
| Sox6 | 9 32E-293 | 5 443824872 | 0 98 | 0 079 | 2 46E-288 | Up_in_MGE | MGE derived cells Pvalb and Sst |
| Satb1 | 2 84E-183 | 3 304169082 | 0 858 | 0 174 | 7 50E-179 | Up_in_MGE | |
| Tshz3 | 5 79E-103 | 2 181106884 | 0 714 | 0 183 | 1 53E-98 | Up_in_MGE | |
| Cers6 | 8 28E-96 | 1 4364747 | 0 916 | 0 534 | 2 19E-91 | Up_in_MGE | |
| Lhx6 | 2 84E-75 | 4 705253349 | 0 367 | 0 017 | 7 50E-71 | Up_in_MGE | |
| Klf5 | 3 28E-74 | 3 229944375 | 0 43 | 0 05 | 8 65E-70 | Up_in_MGE | |
| Etv6 | 1 60E-66 | 2 172219989 | 0 587 | 0 198 | 4 22E-62 | Up_in_MGE | |
| Mafb | 1 68E-65 | 2 193247031 | 0 508 | 0 116 | 4 43E-61 | Up_in_MGE | |
| Mitf | 5 45E-65 | 2 263252661 | 0 557 | 0 188 | 1 44E-60 | Up_in_MGE | |
| Ahr | 6 02E-61 | 1 976944724 | 0 547 | 0 163 | 1 59E-56 | Up_in_MGE | |
| Esrrg | 6 42E-59 | 1 428910188 | 0 849 | 0 567 | 1 69E-54 | Up_in_MGE | |
| Cux2 | 1 23E-54 | 1 29444685 | 0 713 | 0 324 | 3 25E-50 | Up_in_MGE | |
| Klf12 | 1 10E-49 | 0 823599824 | 0 957 | 0 822 | 2 91E-45 | Up_in_MGE | |
| Runx2 | 7 13E-48 | 1 33616659 | 0 753 | 0 436 | 1 88E-43 | Up_in_MGE | |
| Creb3l2 | 1 73E-47 | 3 688577498 | 0 277 | 0 027 | 4 56E-43 | Up_in_MGE | |
| Camta1 | 7 05E-47 | 0 789825964 | 0 926 | 0 74 | 1 86E-42 | Up_in_MGE | |
| Peg3 | 2 56E-43 | 0 695538924 | 0 936 | 0 771 | 6 75E-39 | Up_in_MGE | |
| Mef2c | 4 17E-34 | 0 735773994 | 0 948 | 0 921 | 1 10E-29 | Up_in_MGE | |
| Klf3 | 1 46E-28 | 1 459388084 | 0 373 | 0 134 | 3 86E-24 | Up_in_MGE | |
| Rarb | 3 30E-28 | 1 817244926 | 0 278 | 0 071 | 8 71E-24 | Up_in_MGE | |
| Cux1 | 3 36E-25 | 0 975894866 | 0 551 | 0 297 | 8 86E-21 | Up_in_MGE | |
| Pparg | 6 54E-23 | 1 017111712 | 0 436 | 0 198 | 1 72E-18 | Up_in_MGE | |
| Trerf1 | 5 74E-22 | 0 569705871 | 0 814 | 0 579 | 1 52E-17 | Up_in_MGE | |
| Zfp398 | 7 79E-22 | 0 593783013 | 0 807 | 0 584 | 2 06E-17 | Up_in_MGE | |
| Bcl11a | 1 21E-21 | 0 671789405 | 0 687 | 0 422 | 3 18E-17 | Up_in_MGE | |
| Etv1 | 3 88E-19 | 2 36695352 | 0 165 | 0 036 | 1 02E-14 | Up_in_MGE | |
| Bach1 | 5 60E-18 | 1 270740946 | 0 26 | 0 095 | 1 48E-13 | Up_in_MGE | |
| Zfp30 | 2 32E-17 | 0 592052595 | 0 664 | 0 424 | 6 13E-13 | Up_in_MGE | |
| Prdm1 | 3 48E-17 | 3 283595617 | 0 108 | 0 012 | 9 17E-13 | Up_in_MGE | |
| Zfp169 | 4 58E-17 | 0 932849757 | 0 391 | 0 194 | 1 21E-12 | Up_in_MGE | |
| Thrb | 9 24E-17 | 0 306698508 | 0 973 | 0 912 | 2 44E-12 | Up_in_MGE | |
| Tox | 1 61E-16 | 1 557774117 | 0 43 | 0 277 | 4 24E-12 | Up_in_MGE | |
| Barx2 | 4 23E-16 | 2 117414573 | 0 119 | 0 019 | 1 12E-11 | Up_in_MGE | |
| Arnt2 | 5 33E-16 | 0 443810893 | 0 864 | 0 7 | 1 41E-11 | Up_in_MGE | |
| Ets1 | 1 08E-14 | 2 009966491 | 0 127 | 0 027 | 2 84E-10 | Up_in_MGE | |
| Arx | 2 30E-13 | 0 763779033 | 0 371 | 0 198 | 6 07E-09 | Up_in_MGE | |
| Zfp260 | 6 08E-13 | 0 616688621 | 0 485 | 0 288 | 1 60E-08 | Up_in_MGE | |
| Cbfb | 1 10E-10 | 1 229293558 | 0 156 | 0 058 | 2 92E-06 | Up_in_MGE | |
| Aff1 | 1 13E-10 | 1 640399838 | 0 105 | 0 027 | 2 99E-06 | Up_in_MGE | |

| | | | | | | |
|---|---|---|---|---|---|---|
| Pou6f2 | 1 26E-10 | 1 894131212 | 0 126 | 0 041 | 3 33E-06 | Up_in_MGE |
| Hes1 | 1 84E-10 | 1 415345284 | 0 143 | 0 051 | 4 85E-06 | Up_in_MGE |
| Zfp827 | 2 15E-10 | 0 556083915 | 0 388 | 0 227 | 5 67E-06 | Up_in_MGE |
| Zfp941 | 2 70E-10 | 0 480071835 | 0 535 | 0 345 | 7 12E-06 | Up_in_MGE |
| Trps1 | 1 22E-09 | 0 959975142 | 0 564 | 0 458 | 3 21E-05 | Up_in_MGE |
| Nfkb1 | 2 10E-09 | 0 482398815 | 0 52 | 0 35 | 5 54E-05 | Up_in_MGE |
| Zfp641 | 2 65E-09 | 0 378584564 | 0 667 | 0 493 | 7 00E-05 | Up_in_MGE |
| Zkscan3 | 4 91E-09 | 0 431138233 | 0 486 | 0 32 | 0 00012961 | Up_in_MGE |
| Crebzf | 6 58E-09 | 0 32980222 | 0 844 | 0 681 | 0 00017356 | Up_in_MGE |
| Smad9 | 2 08E-08 | 0 725176231 | 0 171 | 0 078 | 0 00054774 | Up_in_MGE |
| Tfdp2 | 3 30E-08 | 0 411208521 | 0 467 | 0 306 | 0 00087112 | Up_in_MGE |
| Pbx4 | 5 34E-08 | 0 674497822 | 0 26 | 0 149 | 0 00140911 | Up_in_MGE |
| Zfp90 | 6 71E-08 | 0 74318269 | 0 183 | 0 088 | 0 00177126 | Up_in_MGE |
| Zfp27 | 2 52E-07 | 0 583597992 | 0 23 | 0 128 | 0 00663861 | Up_in_MGE |
| Gm14399 | 2 75E-07 | 0 723389802 | 0 184 | 0 094 | 0 00725216 | Up_in_MGE |
| Zbtb7a | 2 91E-07 | 0 600201199 | 0 324 | 0 206 | 0 00769076 | Up_in_MGE |
| Zfp408 | 1 14E-06 | 0 528392987 | 0 299 | 0 184 | 0 03013551 | Up_in_MGE |
| Gm14410 | 1 71E-06 | 0 403924528 | 0 416 | 0 291 | 0 045171 | Up_in_MGE |
| Zfp719 | 1 89E-06 | 0 450203593 | 0 257 | 0 152 | 0 0499051 | Up_in_MGE |
| Zfpm2 | 3 78E-06 | 0 576589214 | 0 369 | 0 265 | 0 09978274 | Up_in_MGE |
| Zbtb14 | 4 74E-06 | 0 989207052 | 0 114 | 0 052 | 0 12496648 | Up_in_MGE |
| Hmg20b | 4 96E-06 | 0 628520771 | 0 105 | 0 044 | 0 13101523 | Up_in_MGE |
| Ascl4 | 5 05E-06 | 0 787409758 | 0 126 | 0 059 | 0 13321895 | Up_in_MGE |
| Zfp955a | 5 06E-06 | 0 340043093 | 0 292 | 0 185 | 0 13353889 | Up_in_MGE |
| Zfp934 | 6 14E-06 | 0 75326714 | 0 14 | 0 071 | 0 1620959 | Up_in_MGE |
| Rfx3 | 8 30E-06 | 0 258758258 | 0 595 | 0 443 | 0 21916302 | Up_in_MGE |
| Zfp949 | 9 20E-06 | 0 368639488 | 0 38 | 0 262 | 0 2426869 | Up_in_MGE |
| Arntl2 | 9 58E-06 | 0 341834473 | 0 286 | 0 183 | 0 25294996 | Up_in_MGE |
| Ncor2 | 1 00E-05 | 0 365713989 | 0 302 | 0 198 | 0 2645042 | Up_in_MGE |
| Zfp961 | 1 21E-05 | 0 378203578 | 0 41 | 0 287 | 0 32052516 | Up_in_MGE |
| Zfp770 | 1 76E-05 | 0 289852807 | 0 317 | 0 212 | 0 46329822 | Up_in_MGE |
| Zkscan2 | 1 80E-05 | 0 293261694 | 0 419 | 0 295 | 0 47565477 | Up_in_MGE |
| Etv5 | 2 64E-05 | 0 413757822 | 0 251 | 0 159 | 0 69587858 | Up_in_MGE |
| Zfp281 | 2 83E-05 | 0 403865875 | 0 155 | 0 085 | 0 74561333 | Up_in_MGE |
| Hsf2 | 2 86E-05 | 0 42305385 | 0 274 | 0 178 | 0 75457254 | Up_in_MGE |
| Nfyc | 2 88E-05 | 0 294132524 | 0 597 | 0 473 | 0 76018148 | Up_in_MGE |
| Cdc5l | 5 62E-05 | 0 362802871 | 0 281 | 0 187 | 1 | Up_in_MGE |
| Zscan22 | 5 90E-05 | 0 31273012 | 0 303 | 0 206 | 1 | Up_in_MGE |
| Esrra | 6 46E-05 | 0 413019557 | 0 284 | 0 19 | 1 | Up_in_MGE |
| Zfp777 | 6 81E-05 | 0 389083772 | 0 186 | 0 112 | 1 | Up_in_MGE |
| Gm14401 | 8 12E-05 | 0 658759603 | 0 116 | 0 06 | 1 | Up_in_MGE |
| Smad7 | 8 45E-05 | 0 361484938 | 0 221 | 0 14 | 1 | Up_in_MGE |
| Zbtb37 | 0 000116915 | 0 423821432 | 0 198 | 0 123 | 1 | Up_in_MGE |
| Zbtb26 | 0 000192202 | 0 370000839 | 0 178 | 0 109 | 1 | Up_in_MGE |
| Zfp775 | 0 000230842 | 0 628625294 | 0 109 | 0 058 | 1 | Up_in_MGE |
| Zfp444 | 0 000244895 | 0 365946613 | 0 243 | 0 163 | 1 | Up_in_MGE |
| Zfp956 | 0 000274912 | 0 312236074 | 0 302 | 0 212 | 1 | Up_in_MGE |
| Zfp943 | 0 000320454 | 0 286175347 | 0 22 | 0 143 | 1 | Up_in_MGE |
| Zbtb17 | 0 000433868 | 0 377941111 | 0 197 | 0 127 | 1 | Up_in_MGE |
| Klf6 | 0 000463004 | 0 52438099 | 0 145 | 0 088 | 1 | Up_in_MGE |
| Zfp128 | 0 000531309 | 0 563688846 | 0 138 | 0 084 | 1 | Up_in_MGE |
| Ybx3 | 0 000558814 | 0 352392965 | 0 252 | 0 176 | 1 | Up_in_MGE |
| Gm14325 | 0 00057647 | 0 273517963 | 0 227 | 0 15 | 1 | Up_in_MGE |
| Sp1 | 0 000721395 | 0 364505948 | 0 151 | 0 094 | 1 | Up_in_MGE |
| Smad1 | 0 000762171 | 0 252967404 | 0 264 | 0 185 | 1 | Up_in_MGE |
| Zfp790 | 0 000814847 | 0 38187448 | 0 105 | 0 058 | 1 | Up_in_MGE |
| Zfp866 | 0 000826337 | 0 288020229 | 0 256 | 0 179 | 1 | Up_in_MGE |
| Zfp868 | 0 001357861 | 0 320483545 | 0 142 | 0 088 | 1 | Up_in_MGE |
| Zfp617 | 0 001367753 | 0 415401824 | 0 117 | 0 07 | 1 | Up_in_MGE |
| Sox5 | 0 001526524 | 0 404228848 | 0 628 | 0 622 | 1 | Up_in_MGE |
| Zfp239 | 0 001957467 | 0 263890998 | 0 241 | 0 171 | 1 | Up_in_MGE |
| Zbtb40 | 0 002006609 | 0 414791809 | 0 134 | 0 085 | 1 | Up_in_MGE |
| Sp9 | 0 002652481 | 0 271331106 | 0 126 | 0 078 | 1 | Up_in_MGE |
| Zfp410 | 0 003036673 | 0 265929743 | 0 191 | 0 133 | 1 | Up_in_MGE |
| Zfp871 | 0 003259777 | 0 28527266 | 0 256 | 0 186 | 1 | Up_in_MGE |
| Zbtb21 | 0 005163403 | 0 339703934 | 0 101 | 0 063 | 1 | Up_in_MGE |
| Zfp93 | 0 00596531 | 0 357265339 | 0 117 | 0 076 | 1 | Up_in_MGE |
| Nfyb | 0 006188828 | 0 337249953 | 0 144 | 0 098 | 1 | Up_in_MGE |
| Zfp276 | 0 007987465 | 0 261574043 | 0 138 | 0 094 | 1 | Up_in_MGE |
| Zbtb1 | 0 008992759 | 0 294630485 | 0 114 | 0 074 | 1 | Up_in_MGE |
| Zfp316 | 0 009824523 | 0 422719082 | 0 123 | 0 084 | 1 | Up_in_MGE |
| Mtf1 | 0 010250132 | 0 255811653 | 0 162 | 0 115 | 1 | Up_in_MGE |
| Zfp94 | 0 012188421 | 0 274682848 | 0 135 | 0 093 | 1 | Up_in_MGE |
| Arid3b | 0 032463367 | 0 310591473 | 0 144 | 0 109 | 1 | Up_in_MGE |

Table 9. Electrphysiological recording parameters and statistical analysis

| Max firing freq (Hz) | | AP amp (mV) | | AP Half W (ms) | |
|---|---|---|---|---|---|
| Ascl1SA6+Bcl2 | Ascl1SA6+Dlx2 | Ascl1SA6+Bcl2 | Ascl1SA6+Dlx2 | Ascl1SA6+Bcl2 | Ascl1SA6+Dlx2 |
| 28 | 25 | 53 7 | 37 26 | 1 86 | 1 14 |
| 30 | 72 | 77 09 | 23 7 | 1 26 | 1 62 |
| 136 | 108 | 89 78 | 20 1 | 0 92 | 1 15 |
| 36 | 62 | 74 58 | 33 7 | 1 47 | 1 12 |
| 134 | 62 | 70 19 | 47 71 | 1 39 | 1 45 |
| 20 | 80 | 57 46 | 25 | 1 89 | 1 59 |
| 160 | 42 | 35 3 | 15 1 | 1 35 | 1 19 |
| 160 | 94 | 46 17 | 39 8 | 0 93 | 1 26 |
| 126 | 142 | 72 34 | 38 67 | 0 84 | 0 97 |
| 174 | 34 | 74 34 | 19 13 | 0 67 | 1 52 |
| 30 | | 51 9 | | 0 83 | |
| 98 | | 46 | | 1 48 | |
| 160 | | 24 4 | | 0 87 | |
| 38 | | 44 | | 0 9 | |

| AHP (mV) | | IR (Mohms) | | RMP (mV) | | Cm (pF) | |
|---|---|---|---|---|---|---|---|
| Ascl1SA6+Bcl2 | Ascl1SA6+Dlx2 | Ascl1SA6+Bcl2 | Ascl1SA6+Dlx2 | Ascl1SA6+Bcl2 | Ascl1SA6+Dlx2 | Ascl1SA6+Bcl2 | Ascl1SA6+Dlx2 |
| 30 55 | 12 8 | 1600 | 294 | -45 | -73 | 11 4 | 14 9 |
| 63 49 | 4 33 | 2000 | 2000 | -53 | -47 | 9 05 | 15 7 |
| 78 73 | 16 39 | 978 | 1900 | -40 | -49 | 8 77 | 20 84 |
| 46 55 | 10 9 | 1140 | 2153 | -32 | -55 | 9 28 | 20 2 |
| 98 8 | 3 39 | 1820 | 590 | -44 | -30 | 2 26 | 24 7 |
| 54 6 | 6 68 | 2010 | 1070 | -45 | -41 | 5 22 | 18 4 |
| 32 1 | 17 12 | 1373 | 1521 | -65 | -60 | 4 2 | 41 3 |
| 43 21 | 6 37 | 830 | 716 | -45 | -43 | 6 28 | 21 9 |
| 46 85 | 11 26 | 879 | 1442 | -45 | -48 | 7 12 | 18 9 |
| 60 56 | 18 9 | 1037 | 904 | -43 | -36 | 10 9 | 18 |
| 31 7 | | 1685 | | -77 | | 13 1 | |
| 12 72 | | 1300 | | -51 | | 22 3 | |
| 5 49 | | 2400 | | -57 | | 33 5 | |
| 19 9 | | 1630 | | -40 | | 15 4 | |

| Table Analyzed | Max firing freq (Hz) | AHP (mV) | Cm (pF) | IR (Mohms) | RMP (mV) | AP Half W (ms) | AP amp (mV) |
|---|---|---|---|---|---|---|---|
| Column B | Ascl1SA6+Dlx2 | Ascl1SA6+Dlx2 | Ascl1SA6+Dlx2 | Ascl1SA6+Dlx2 | Ascl1SA6+Dlx2 | Ascl1SA6+Dlx2 | Ascl1SA6+Dlx2 |
| vs. | vs. | vs. | vs. | vs. | vs. | vs. | vs. |
| Column A | Ascl1SA6+Bcl2 | Ascl1SA6+Bcl2 | Ascl1SA6+Bcl2 | Ascl1SA6+Bcl2 | Ascl1SA6+Bcl2 | Ascl1SA6+Bcl2 | Ascl1SA6+Bcl2 |
| paired t test with Welch's correction | | | | | | | |
| P value | 0 2611 | 0 0002 | 0 005 | 0 3761 | 0 9183 | 0 3932 | 0 0001 |
| P value summary | ns | *** | ** | ns | ns | ns | *** |
| Significantly different (P < 0.05 | No | Yes | Yes | No | No | No | Yes |
| One- or two-tailed P value? | Two-tailed | Two-tailed | Two-tailed | Two-tailed | Two-tailed | Two-tailed | Two-tailed |
| Welch-corrected t, df | t=1.154, df=21.37 | t=4.798, df=14.66 | t=3.145, df=20.44 | t=0.9106, df=15.87 | t=0.1040, df=18.62 | t=0.8715, df=21.23 | t=4.716, df=21.36 |
| How big is the difference? | | | | | | | |
| Mean of column A | 95 | 44 66 | 11 34 | 1477 | -48 71 | 1 19 | 58 38 |
| Mean of column B | 72 1 | 10 81 | 21 48 | 1259 | -48 2 | 1 301 | 30 02 |
| ference between means (B - A) ± | -22.90 ± 19.84 | -33.85 ± 7.054 | 10.14 ± 3.225 | -218.3 ± 239.7 | 0.5143 ± 4.946 | 0.1110 ± 0.1274 | -28.36 ± 6.013 |
| 95% confidence interval | -64.11 to 18.31 | -48.91 to -18.78 | 3.424 to 16.86 | -726.8 to 290.3 | -9.852 to 10.88 | -0.1537 to 0.3757 | -40.85 to -15.87 |
| R squared (eta squared) | 0 05869 | 0 6109 | 0 3261 | 0 04966 | 0 0005802 | 0 03454 | 0 5101 |
| F test to compare variances | | | | | | | |
| F, DFn, Dfd | 2.913, 13, 9 | 21.45, 13, 9 | 1.168, 13, 9 | 1.790, 9, 13 | 1.158, 9, 13 | 3.039, 13, 9 | 2.922, 13, 9 |
| P value | 0 1138 | <0.0001 | 0 8344 | 0 3295 | 0 7857 | 0 1008 | 0 1128 |
| P value summary | ns | **** | ns | ns | ns | ns | ns |
| Significantly different (P < 0.05 | No | Yes | No | No | No | No | No |
| Data analyzed | | | | | | | |
| Sample size, column A | 14 | 14 | 14 | 14 | 14 | 14 | 14 |
| Sample size, column B | 10 | 10 | 10 | 10 | 10 | 10 | 10 |

Table 10.

a

Soma size and primary processes of induced neurons SA6/Bcl2 versus SA6/Dlx2 at 4 wpi

| Bcl2 | Dlx2 | Bcl2 | Dlx2 |
|---|---|---|---|
| 101 | 92 | 59 | 67 |
| 84 | 79 | 52 | 84 |
| 87 | 181 | 44 | 118 |
| 99 | 69 | 39 | 56 |
| 77 | 66 | 41 | 90 |
| 82 | 66 | 57 | 69 |
| 78 | 81 | 45 | 77 |
| 113 | 104 | 35 | 92 |
| 86 | 100 | 41 | 60 |
| 69 | 100 | 52 | 75 |
| 59 | 114 | 28 | 37 |
| 62 | 104 | 43 | 90 |
| 88 | 104 | 35 | 79 |
| 65 | 101 | 48 | 66 |
| 71 | 171 | 36 | 77 |
| 59 | 107 | 54 | 57 |
| 66 | 109 | 41 | 53 |
| 56 | 86 | 42 | 65 |
| 59 | 104 | 52 | 60 |
| 83 | 164 | 49 | 52 |
| 82 | 59 | 50 | 83 |
| 67 | 122 | 50 | 61 |
| 86 | 81 | 49 | 95 |
| 66 | 99 | 49 | 78 |
| 72 | 69 | 59 | 88 |
| 122 | 93 | 100 | 93 |
| 86 | 105 | 92 | 86 |
| 75 | 94 | 38 | 76 |
| 52 | 93 | 35 | 96 |
| 72 | 76 | 39 | 105 |
| 78 | 100 | 33 | 95 |
| 63 | 98 | 49 | 72 |
| 69 | 76 | 41 | 95 |
| 88 | 117 | 48 | 109 |
| 114 | 84 | 35 | 133 |
| 104 | 75 | 86 | 102 |
| 88 | 77 | 74 | 85 |
| 109 | 140 | 113 | 93 |
| 74 | 105 | 69 | 99 |
| 72 | 83 | 93 | 116 |
| 67 | 67 | 78 | 88 |
| 91 | 58 | 59 | 81 |
| 72 | 65 | 56 | 69 |
| 75 | 59 | 64 | 93 |
| 71 | 71 | 62 | 74 |
| 84 | 107 | 51 | 109 |
| 49 | 119 | 50 | 101 |
| 71 | 74 | 58 | 88 |
| 71 | 74 | 83 | 75 |
| 66 | 60 | 66 | 90 |
| 82 | 59 | 51 | 75 |
| 89 | 90 | 45 | 78 |
| 85 | 79 | 54 | 106 |
| 79 | 85 | 80 | 69 |
| 101 | 93 | 65 | |
| 82 | 60 | 93 | |
| 103 | 118 | 82 | |
| 70 | 77 | 59 | |
| 47 | 93 | 83 | |
| 42 | 90 | 69 | |
| 67 | 81 | 77 | |
| 69 | 102 | 75 | |
| 70 | 100 | 61 | |
| 44 | 92 | 49 | |
| 64 | 120 | 82 | |
| 55 | 88 | 75 | |
| 94 | 88 | 85 | |
| 70 | 102 | 52 | |
| 66 | 90 | 65 | |
| 90 | 91 | 66 | |
| 69 | 113 | 50 | |
| 86 | 96 | 49 | |
| 56 | 98 | 47 | |
| 80 | 81 | 42 | |
| 58 | | 45 | |
| 84 | | 52 | |
| 65 | | 45 | |
| 62 | | 43 | |
| 89 | | 52 | |
| | | 49 | |

b

Primary Pocesses Number

| Bc2 | Dlx2 |
|---|---|
| 2 | 4 |
| 2 | 3 |
| 2 | 3 |
| 2 | 2 |
| 1 | 4 |
| 1 | 4 |
| 2 | 3 |
| 2 | 3 |
| 2 | 3 |
| 2 | 4 |
| 2 | 4 |
| 2 | 4 |
| 2 | 2 |
| 1 | 4 |
| 4 | 1 |
| 2 | 3 |
| 2 | 3 |
| 2 | 1 |
| 2 | 1 |
| 3 | 3 |
| 3 | 4 |
| 3 | 5 |
| 5 | 2 |
| 1 | 4 |
| 3 | 4 |
| 2 | 1 |
| 2 | 4 |
| 2 | 2 |
| 1 | 3 |
| 3 | 3 |
| 2 | 2 |
| 2 | 4 |
| 5 | 3 |
| 5 | 5 |
| 2 | 4 |
| 2 | 4 |
| 2 | 3 |
| 2 | 2 |
| 3 | 3 |
| 3 | 4 |
| 2 | 4 |
| 3 | 3 |
| 4 | 3 |
| 3 | 5 |
| 2 | 1 |
| 3 | 2 |
| 2 | 1 |
| 2 | |
| 1 | |

c Stastists

| Table Analyzed | Soma size 4 wpi | rimary proccesses 4 w |
|---|---|---|
| Column B | SA6-Dlx2 | SA6-Dlx2 |
| vs. | vs. | vs. |
| Column A | SA6-Bcl2 | SA6-Bcl2 |
| Unpaired t test with Welch's correction | | |
| P value | <0.0001 | <0.0001 |
| P value summary | **** | **** |
| Significantly different (P < 0.05)? | Yes | Yes |
| One- or two-tailed P value? | Two-tailed | Two-tailed |
| Welch-corrected t, df | t=8.080, df=109.8 | t=11.85, df=138.1 |
| How big is the difference? | | |
| Mean of column A | 57 05 | 1 638 |
| Mean of column B | 82 96 | 2 736 |
| Difference between means (B - A) ± SE | 25.91 ± 3.207 | 1.098 ± 0.09267 |
| 95% confidence interval | 19.56 to 32.27 | 0.9150 to 1.282 |
| R squared (eta squared) | 0 3728 | 0 5043 |
| F test to compare variances | | |
| F, DFn, Dfd | 1.110, 53, 79 | 1.484, 109, 582 |
| P value | 0 6661 | 0 0047 |
| P value summary | ns | ** |
| Significantly different (P < 0.05)? | No | Yes |
| Data analyzed | | |
| Sample size, column A | 80 | 583 |
| Sample size, column B | 54 | 110 |

Layer quantification of Induced Neurons at 2 and 4 wpi

Table 11.

| 2wpi | | | |
|---|---|---|---|
| Bcl2 | 1 | 2,3 | >4 |
| rep1 | 49 73 | 26 89 | 23 38 |
| rep2 | 57 71 | 23 03 | 19 26 |
| rep3 | 58 40 | 18 82 | 22 78 |
| average | **55 28** | **22 91** | **21 81** |

| 2wpi | | | |
|---|---|---|---|
| Dlx2 | 1 | 2,3 | >4 |
| rep1 | 15 62 | 56 64 | 27 74 |
| rep2 | 30 48 | 49 81 | 19 71 |
| rep3 | 28 11 | 52 10 | 19 80 |
| rep4 | 31 20 | 42 73 | 26 07 |
| average | **26 35** | **50 32** | **23 33** |

| 4wpi | | | |
|---|---|---|---|
| Bcl2 | 1 | 2,3 | >4 |
| rep1 | 62 40 | 26 17 | 11 43 |
| rep2 | 62 30 | 19 28 | 18 42 |
| rep3 | 53 61 | 22 36 | 24 04 |
| rep4 | 50 66 | 18 84 | 30 50 |
| average | **57 24** | **21 66** | **21 10** |

| 4wpi | | | |
|---|---|---|---|
| Dlx2 | 1 | 2,3 | >4 |
| rep1 | 28 59 | 52 78 | 18 63 |
| rep2 | 28 93 | 46 31 | 24 76 |
| rep3 | 24 28 | 52 44 | 23 28 |
| rep4 | 22 56 | 59 73 | 17 72 |
| average | **26 09** | **52 81** | **21 10** |

| mouse | total number of iNs/ layer | | | | percentage of iNs/ layer | | | | summary | | |
|---|---|---|---|---|---|---|---|---|---|---|---|
| Bcl2 2wpi | I | II/III | >IV | | I | II/III | >IV | | I | II/III | >IV |
| 1a | 122 | 60 | 33 | 215 | 56 74418605 | 27 90697674 | 15 34883721 | | 49 72732916 | 26 88983334 | 23 3828375 |
| 1b | 208 | 126 | 153 | 487 | 42 71047228 | 25 87268994 | 31 41683778 | | 57 71069281 | 23 03288292 | 19 25642428 |
| | | | | | **49 72732916** | **26 88983334** | **23 3828375** | | 58 40462323 | 18 81598107 | 22 7793957 |
| 2a | 121 | 47 | 49 | 217 | 55 76036866 | 21 65898618 | 22 58064516 | | **55 28088173** | **22 91289911** | **21 80621916** |
| 2b | 176 | 72 | 47 | 295 | 59 66101695 | 24 40677966 | 15 93220339 | | | | |
| | | | | | **57 71069281** | **23 03288292** | **19 25642428** | | | | |
| 3a | 111 | 45 | 45 | 201 | 55 2238806 | 22 3880597 | 22 3880597 | | | | |
| 3b | 101 | 25 | 38 | 164 | 61 58536585 | 15 24390244 | 23 17073171 | | | | |
| | | | | | **58 40462323** | **18 81598107** | **22 7793957** | | | | |
| Dlx2 2wpi | I | II/III | >IV | | I | II/III | >IV | | I | II/III | >IV |
| 4a | 12 | 50 | 29 | 91 | 13 18681319 | 54 94505495 | 31 86813187 | | 15 62118437 | 56 63919414 | 27 73962149 |
| 4b | 13 | 42 | 17 | 72 | 18 05555556 | 58 33333333 | 23 61111111 | | 30 47603902 | 49 80916031 | 19 71480068 |
| | | | | | **15 62118437** | **56 63919414** | **27 73962149** | | 28 10728745 | 52 0951417 | 19 79757085 |
| 5a | 38 | 65 | 28 | 131 | 29 00763359 | 49 61832061 | 21 3740458 | | 31 20155039 | 42 72609819 | 26 07235142 |
| 5b | 46 | 72 | 26 | 144 | 31 94444444 | 50 | 18 05555556 | | **26 35151531** | **50 31739858** | **23 33108611** |
| | | | | | **30 47603902** | **49 80916031** | **19 71480068** | | | | |
| 6a | 30 | 58 | 16 | 104 | 28 84615385 | 55 76923077 | 15 38461538 | | | | |
| 6b | 26 | 46 | 23 | 95 | 27 36842105 | 48 42105263 | 24 21052632 | | | | |
| | | | | | **28 10728745** | **52 0951417** | **19 79757085** | | | | |
| 7a | 30 | 34 | 26 | 90 | 33 33333333 | 37 77777778 | 28 88888889 | | | | |
| 7b | 25 | 41 | 20 | 86 | 29 06976744 | 47 6744186 | 23 25581395 | | | | |
| | | | | | **31 20155039** | **42 72609819** | **26 07235142** | | | | |
| Bcl2 4wpi | I | II/III | >IV | | I | II/III | >IV | | I | II/III | >IV |
| 12a | 24 | 10 | 4 | 38 | 63 15789474 | 26 31578947 | 10 52631579 | | 62 40086518 | 26 17159337 | 11 42754146 |
| 12b | 45 | 19 | 9 | 73 | 61 64383562 | 26 02739726 | 12 32876712 | | 62 29909154 | 19 28138831 | 18 41952015 |
| | | | | | **62 40086518** | **26 17159337** | **11 42754146** | | 53 60576923 | 22 35576923 | 24 03846154 |
| 13a | 98 | 25 | 36 | 159 | 61 63522013 | 15 72327044 | 22 64150943 | | 50 66365627 | 18 83552271 | 30 50082102 |
| 13b | 102 | 37 | 23 | 162 | 62 96296296 | 22 83950617 | 14 19753086 | | **57 24234555** | **21 6610684** | **21 09658604** |
| | | | | | **62 29909154** | **19 28138831** | **18 41952015** | | | | |
| 14a | 87 | 37 | 84 | 208 | 41 82692308 | 17 78846154 | 40 38461538 | | | | |
| 14b | 68 | 28 | 8 | 104 | 65 38461538 | 26 92307692 | 7 692307692 | | | | |
| | | | | | **53 60576923** | **22 35576923** | **24 03846154** | | | | |
| 15a | 90 | 40 | 44 | 174 | 51 72413793 | 22 98850575 | 25 28735632 | | | | |
| 15b | 125 | 37 | 90 | 252 | 49 6031746 | 14 68253968 | 35 71428571 | | | | |
| | | | | | **50 66365627** | **18 83552271** | **30 50082102** | | | | |
| Dlx2 4wpi | I | II/III | >IV | | | | | | I | II/III | >IV |
| 8a | 10 | 20 | 6 | 36 | 27 77777778 | 55 55555556 | 16 66666667 | | 28 59477124 | 52 77777778 | 18 62745098 |
| 8b | 10 | 17 | 7 | 34 | 29 41176471 | 50 | 20 58823529 | | 28 92857143 | 46 30952381 | 24 76190476 |
| | | | | | **28 59477124** | **52 77777778** | **18 62745098** | | 24 28070175 | 52 43859649 | 23 28070175 |
| 9a | 19 | 27 | 14 | 60 | 31 66666667 | 45 | 23 33333333 | | 22 55639098 | 59 72744361 | 17 71616541 |
| 9b | 11 | 20 | 11 | 42 | 26 19047619 | 47 61904762 | 26 19047619 | | **26 09010885** | **52 81333542** | **21 09655573** |
| | | | | 0 | **28 92857143** | **46 30952381** | **24 76190476** | | | | |
| 10a | 12 | 27 | 11 | 50 | 24 | 54 | 22 | | | | |
| 10b | 14 | 29 | 14 | 57 | 24 56140351 | 50 87719298 | 24 56140351 | | | | |
| | | | | 0 | **24 28070175** | **52 43859649** | **23 28070175** | | | | |
| 11a | 9 | 23 | 6 | 38 | 23 68421053 | 60 52631579 | 15 78947368 | | | | |
| 11b | 12 | 33 | 11 | 56 | 21 42857143 | 58 92857143 | 19 64285714 | | | | |
| | | | | | **22 55639098** | **59 72744361** | **17 71616541** | | | | |

| Layer | Timepoint | BCL2 mean (% | Dlx2 mean (%) | t-value | df | p-value | Significance |
|---|---|---|---|---|---|---|---|
| Layer I | 2 wpi | 55 3 | 26 4 | 6 32 | 4 98 | 0 00149 | ** |
| Layer I | 4 wpi | 57 2 | 26 1 | 9 16 | 4 54 | 0 000426 | *** |
| Layer 2/3 | 2 wpi | 22 9 | 50 3 | -7 37 | 5 | 0 000725 | *** |
| Layer 2/3 | 4 wpi | 21 7 | 52 8 | -9 66 | 5 | 0 000202 | *** |
| Layer 4/5/6 | 2 wpi | 21 8 | 23 3 | -0 62 | 4 69 | 0 564 | n.s. |
| Layer 4/5/6 | 4 wpi | 21 1 | 21 1 | 0 | 4 05 | 1 | n.s. |
